# A Generalizable Nanopore Sensor for Highly Specific Protein Detection at Single-Molecule Precision

**DOI:** 10.1101/2022.10.12.511930

**Authors:** Mohammad Ahmad, Jeung-Hoi Ha, Lauren A. Mayse, Maria F. Presti, Aaron J. Wolfe, Kelsey J. Moody, Stewart N. Loh, Liviu Movileanu

**Affiliations:** Department of Physics, Syracuse University, 201 Physics Building, Syracuse, New York 13244-1130, USA; Department of Biochemistry and Molecular Biology, State University of New York - Upstate Medical University, 4249 Weiskotten Hall, 766 Irving Avenue, Syracuse, New York 13210, USA; Ichor Life Sciences, Inc., 2651 US Route 11, LaFayette, New York 13084, USA; Lewis School of Health Sciences, Clarkson University, 8 Clarkson Avenue, Potsdam, New York 13699, USA; Department of Chemistry, State University of New York, College of Environmental Science and Forestry, 1 Forestry Dr., Syracuse, New York 13210, USA; Department of Biomedical and Chemical Engineering, Syracuse University, 329 Link Hall, Syracuse, New York 13244, USA; The BioInspired Institute, Syracuse University, Syracuse, New York 13244, USA

**Keywords:** Protein Biomarker, Ion channel, Protein dynamics, Single-molecule electrophysiology, Protein engineering, Cancer, Monobody

## Abstract

Protein detection and biomarker profiling have wide-ranging implications in many areas of basic research and molecular diagnostics. Substantial progress has been made in protein analytics using nanopores and the resistive-pulse technique. Yet, a long-standing challenge is implementing specific binding interfaces for detecting proteins without the steric hindrance of the pore interior. To overcome this technological difficulty, we formulate a new class of sensing elements made of a programmable antibody-mimetic binder fused to a monomeric protein nanopore. This way, such a modular design significantly expands the utility of nanopore sensors to numerous proteins while preserving their architecture, specificity, and sensitivity. We prove the power of this approach by developing and validating nanopore sensors for protein analytes that drastically vary in size, charge, and structural complexity. These analytes produce unique electrical signatures that depend on their identity and quantity and the binder-analyte assembly at the nanopore tip. From a practical point of view, our sensors unambiguously probe protein recognition events without the necessity of using any additional exogenous tag. The outcomes of this work will impact biomedical diagnostics by providing a fundamental basis and tools for protein biomarker detection in biofluids.

## Introduction

Identifying and quantifying protein biomarkers is a pressing demand in precision and personalized medicine.^1, 2^ Recent advancements in functional proteomics indicate that there are yet numerous unexplored proteins with potential implications for the progression of pathological conditions.^3^ Therefore, there is an increasing need to create highly specific and sensitive protein sensing approaches that employ rapid signal responses to various biochemical stimuli.^4^ Molecular details of protein detection are illuminated using single-molecule methods.^5-7^ In particular, single-molecule sensing with nanopores^8-14^ using the resistive-pulse technique^15^ is adaptable to parallel recording technologies.^16, 17^ Despite such a significant benefit, this approach usually requires the targeted proteins to partition into the nanopore interior. Hence, the detection is conducted under steric restrictions of the nanopore confinement, potentially impairing the strength of specific interactions.

Detecting single proteins outside the nanopore is a practical alternative to sampling the complexity of protein recognition events.^18-21^ This task would necessitate an external protein binder (e.g., receptor) covalently attached to a nanopore. However, a transducing mechanism is needed to convert the physical captures and releases of a protein analyte (e.g., its ligand) into a specific electrical signature of the sensor. In addition, changing the system to a new binder-analyte pair requires a lengthy and tedious optimization process that includes amplified difficulties. The heterogeneous architecture, size, charge, and other traits of different binders need extensive protein engineering. This prerequisite is critical for each sensor for a given protein analyte. Earlier studies have suggested that these protein sensors may be limited to established protein fragments of ∼100 residues.^22, 23^ For example, large protein binders likely induce additional steric constraints, precluding the clearance of the space around the pore opening. Moreover, the interaction interface of the binder must be fully accessible to the protein analyte.

To address these shortcomings, we propose a new class of sensing elements for probing proteins at a single-detector precision. These sensors will have an antibody-mimetic protein binder engineered on the tFhuA nanopore,^22^ a monomeric β-barrel scaffold, via a flexible tether. This strategy maintains the sensor’s architecture, high sensitivity, and specificity while featuring its generalization to numerous protein analytes. We demonstrate that by changing only the binding interface, a novel binder-containing nanopore sensor can be obtained and readily implemented into the detection of a specific protein (**Fig. 1**). Here, the binder is a monobody,^24-28^ a recombinant protein based on the 94-residue fibronectin type III (FN3) domain.^29^ Using the monobody-based nanopore sensors with varying binding interfaces, it is possible to detect different proteins that vary substantially in their structural and functional properties. When subjected to a biofluid, this class of sensing elements can report the presence of a protein biomarker at a single-molecule level. Finally, this tactic will not only enable overcoming the abovementioned challenges but will also motivate the widespread applications of these sensors.

**Figure 1.**
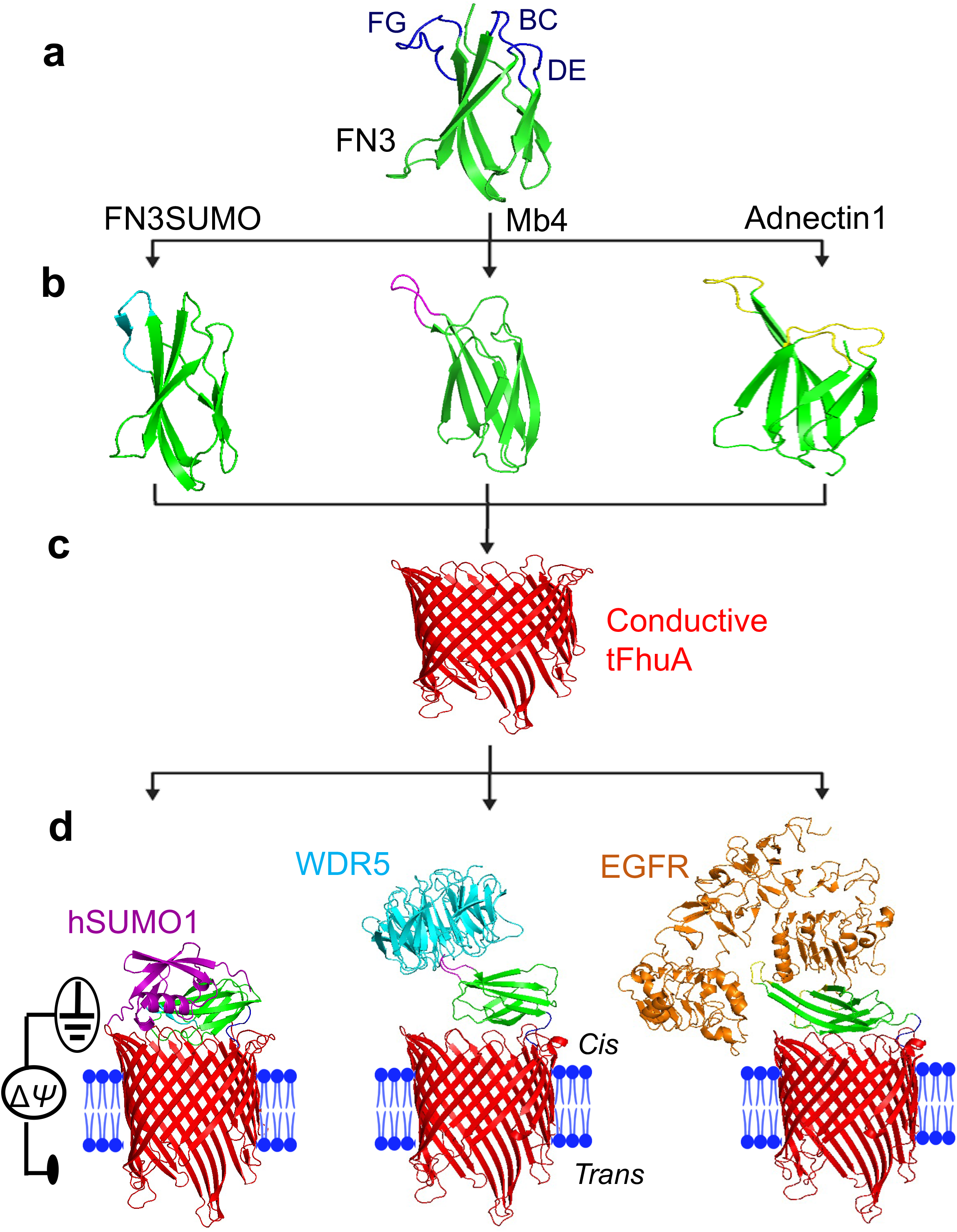
Rational protein design of a generalizable nanopore sensor for protein detection. **(a)** A tenth fibronectin type-III domain (FN3)^29, 59^ or monobody with the FG, BC, and DE loops highlighted in blue. **(b)** The FN3 variants, where cyan, magenta, and yellow were used to represent the binding loops in FN3SUMO,^41^ Mb4,^42^ and Adnectin1,^43^ respectively. **(c)** tFhuA,^22^ a monomeric β barrel with average internal diameters of ∼2.6 nm and ∼3.9 nm, measured from side chain to side chain. **(d)** Functional reconstitution of nanopore sensors into a lipid bilayer. The hSUMO1-binding monobody (FN3SUMO) is a single-polypeptide unit that comprises tFhuA, a (GGS)_2_tether, and FN3SUMO (*left*). WDR5-binding monobody (Mb4; *center*) and EGFR-binding monobody (Adnectin1; *right*) were also fused to tFhuA in the same way as FN3SUMO. The monobody-analyte complexes are shown as well. The structures of all sensors were predicted by AlphaFold2.^34, 35^

## Results and Discussion

### Development of monobody-based nanopore sensors

In this study, we developed monobody-based nanopore sensors for three targeted analytes: (i) human small ubiquitin-related modifier 1 (hSUMO1), a model protein with implications in various cellular processes, such as DNA damage repair, chromosome dynamics, and cell cycle;^30-33^ (ii) WD40 repeat protein 5 (WDR5),^34, 35^ a chromatin-associated protein hub involved in the epigenetic regulation of histone 3 lysine 4 (H3K4) methylation; (iii) epidermal growth factor receptor (EGFR),^36^ a prognosis protein biomarker in lung, colorectal, and breast cancers^37-40^ (**Fig. 1**; **Supplementary Table S1**). Therefore, we created three sensors using FN3SUMO,^41^ Mb4,^42^ and Adnectin1^43^ monobodies as binders against hSUMO1, WDR5, and the ectodomain of EGFR, respectively. These monobody-based sensors are denoted by FN3SUMO-tFhuA, Mb4-tFhuA, and Adnectin1-tFhuA, respectively (**Supplementary Fig. S1**).

Here, we employed AlphaFold2, an artificial intelligence approach to predict the overall three-dimensional conformation of a folded protein using its amino acid sequence.^44, 45^ The most suited structural model for FN3-tFhuA was reached when the predicted Local Distance Difference Test (pLDDT), a confidence score per residue, was between 80 and 100 for most residues (**Supplementary Fig. S2ab**). This model illustrates that FN3 orients almost perpendicularly on the central axis of tFhuA (**Supplementary Fig. S2c**). This finding is likely due to long-range electrostatic interactions between clusters of negative charges on tFhuA β turns and positive charges on FN3 loops (**Supplementary Fig. S3**). Similar results were obtained with FN3SUMO-tFhuA, Mb4-tFhuA, and Adnectin1-tFhuA (**Supplementary Fig. S4**).

Therefore, FN3 monobodies in all sensors potentially block a substantial ionic flow through tFhuA. Inspecting all sensors at a transmembrane potential of +40 mV revealed a relatively quiet single-channel electrical current recorded with FN3SUMO-tFhuA and Mb4-tFhuA, and a slightly noisy signal acquired with Adnectin1-tFhuA (**Supplementary Figs. S5-S6**). The unitary conductance of FN3SUMO-tFhuA, Mb4-tFhuA, and Adnectin1-tFhuA were (mean ± s.d.) 0.81 ± 0.03 nS, 0.99 ± 0.04 nS, and 0.90 ± 0.02 nS (**Supplementary Table S2**), respectively. These are significant conductance reductions compared to the unmodified tFhuA (1.52 ± 0.10 nS) (**Supplementary Fig. S7**).^21^ This finding is in accord with the predictions made by AlphaFold2.

### Real-time and label-free detection of hSUMO1 using FN3SUMO-tFhuA

FN3SUMO-tFhuA was functionally reconstituted into a lipid membrane at an applied transmembrane potential of +40 mV. The presence of hSUMO1 in the *cis* compartment at nanomolar concentrations produced frequent current blockades (**Fig. 2a**; **Supplementary Fig. S8**) between O_on_ open substate and O_off_ closed substate. Their normalized current amplitude, *A*/*I*_0_, was (91.5 ± 0.7)%. Here, *I*_0_ and *A* denote the single-channel current of the hSUMO1-released substate and the current amplitude of hSUMO1-produced current blockades, respectively (**Fig. 2bc**). In addition, infrequent and brief current spikes were observed when hSUMO1 was added to the *cis* side of an unmodified tFhuA-containing bilayer (**Supplementary Fig. S9**). Taken together, these negative-control measurements indicate that hSUMO1 did not produce any significant current blockades due to nonspecific interactions with the *cis* opening of the nanopore.

**Figure 2.**
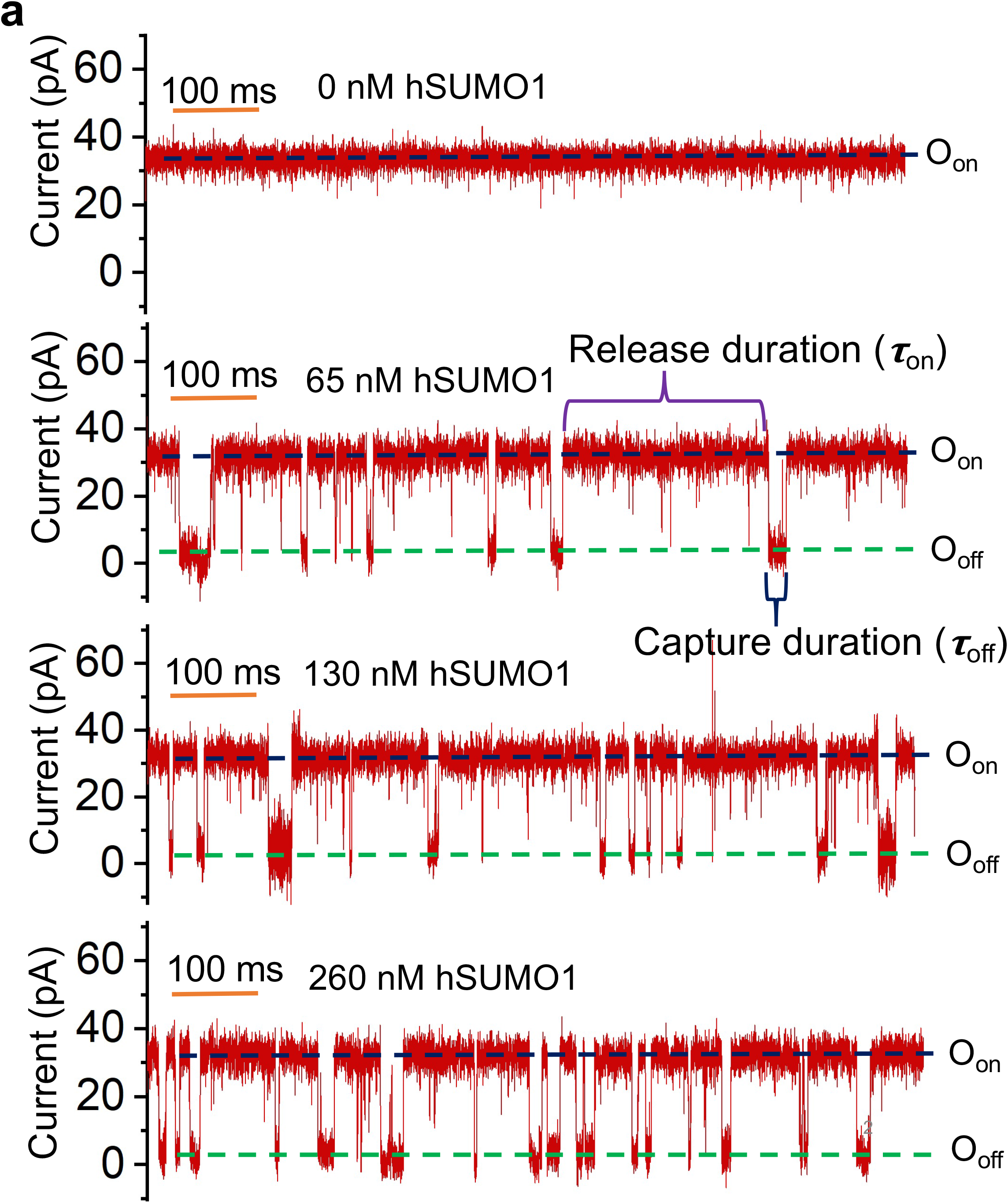
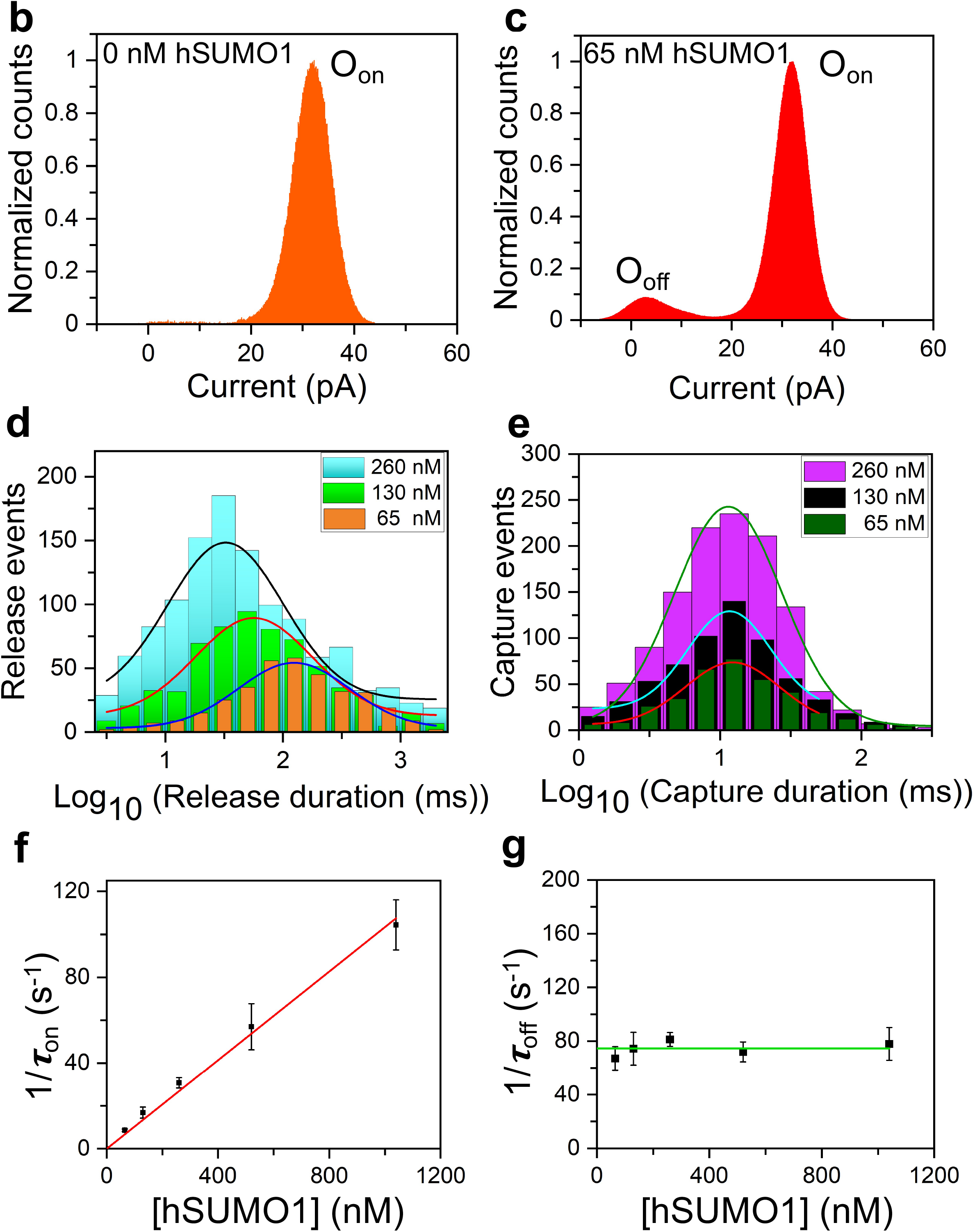
Real-time and label-free detection of hSUMO1. **(a)** Representative single-channel electrical traces of FN3SUMO-tFhuA in the presence of 0, 65, 130, and 260 nM hSUMO1. O_on_ and O_off_ are the hSUMO1-released and hSUMO1-captured substates, respectively. hSUMO1 was added to the *cis* compartment, which was grounded. These single-channel electrical signatures were replicated in *n* = 3 independent experiments. The applied transmembrane potential was +40 mV. Single-channel electrical traces were further low-pass filtered at 3 kHz using an 8-pole Bessel filter. **(b)** A representative all-point current histogram of the O_on_ substate of FN3SUMO-tFhuA. The current amplitude (mean ± s.e.m.) of the O_on_ substate was 32.1 ± 0.1 pA. **(c)** A representative all-point current histogram of the O^on^ and O^off^ substates of FN3SUMO-tFhuA at 65 nM hSUMO1. The current amplitude (mean ± s.e.m.) of the O_**b**off_ substate was 2.9 ± 0.1 pA. **(d)** Semilogarithmic histograms of the hSUMO1-released durations (*τ*_on_) at various hSUMO1 concentrations, [hSUMO1]. *τ*_on_ (mean ± s.e.m.) were 125 ± 4 ms (number of events: *N* = 349), 69 ± 5 ms (*N* = 623), and 35 ± 1 ms (*N* = 1168) at [hSUMO1] values of 65 nM, 130 nM, and 260 nM, respectively. **(e)** Semilogarithmic histograms of the hSUMO1-captured durations (*τ*_off_) at various [hSUMO1] values. *τ*_off_ (mean ± s.e.m.) were 15 ± 1 ms (*N* = 354 events), 16 ± 1 ms (*N* = 633), and 14 ± 1 ms (*N* = 1180) at [hSUMO1] values of 65 nM, 130 nM, and 260 nM, respectively. **(f)** Dependence of the event frequency in the form of 1/*τ*_on_ on [hSUMO1]. The slope of the linear fit of 1/*τ*^on^ versus [hSUMO1] is the association rate constant, *k*^on^, of hSUMO1-FN3SUMO interactions because *k*_on_ = 1/(*τ*_on_ [hSUMO1]). **(g)** Dependence of 1/*τ*_off_ on [hSUMO1]. The horizontal line is an average fit of the (1/*τ*_off_) data points recorded for various [hSUMO1] values. Data points in panels (f) and (g) represent mean ± s.d. obtained from *n* = 3 different experiments.

Moreover, hSUMO1-captured events were noted concentration-dependent (**Fig. 2a**). hSUMO1-released and hSUMO1-captured events recorded with FN3SUMO-tFhuA corresponded to the open-substate, O_on_, and closed-substate, O_off_, respectively. However, hSUMO1-captured events were not detectable when hSUMO1 was added to the *trans* compartment (**Supplementary Fig. S10**), confirming that tFhuA and its derivatives insert into the membrane with a single orientation.^46^

Next, we pursued detailed statistical analyses of the hSUMO1-released and hSUMO1-captured durations, whose mean values were denoted by *τ*_on_ and *τ*_off_, respectively. The maximum likelihood method^47^ and logarithm likelihood ratio (LLR) tests^48, 49^ were employed to determine the distribution model of these time constants. Durations of hSUMO1-released and hSUMO1-captured events showed a single-exponential distribution in the form of a single peak in a semilogarithmic plot (**Fig. 2de**). Although the bin size was identical in these histograms, we represented them differently for clarity. Increasing the hSUMO1 concentration, [hSUMO1], decreased the *τ*_on_ but did not alter *τ*_off_ (**Supplementary Table S3**). The association rate constants, *k*_on_, were consistent for all [hSUMO1] values (**Supplementary Table S4**). Here, *k*_on_ = 1/([hSUMO1] *τ*_on_). In addition, the frequency of hSUMO1-captured events, *f*, where *f* = 1/*τ*_on_, was proportional to [hSUMO1] in a ratio 1:1 (**Fig. 2f**), indicating a bimolecular association process of the hSUMO1-FN3SUMO complex. Using the linear fit of *f*([hSUMO1]), we obtain a *k*_on_ value (mean ± s.e.m.) of (1.12 ± 0.02) × 10^8^ M^-1^s^-1^. Dissociation rate constant *k*_off_ was determined as reciprocal of the mean hSUMO1-captured durations (1/*τ*_off_). This value was independent of [hSUMO1] (**Fig. 2g**; **Supplementary Table S4**), suggesting a unimolecular dissociation mechanism of the hSUMO1-FN3SUMO complex. A linear fit of *k*_off_([hSUMO1]) versus [hSUMO1] resulted in its mean ± s.e.m. of 74.5 ± 2.4 s^-1^, to yield an equilibrium dissociation constant (*K*_D_) of 665 ± 24 nM (**Supplementary Table S5**).

### Detection of a chromatin-associated protein hub using Mb4-tFhuA

We employed the same approach and experimental conditions to detect WDR5 using a functionally reconstituted Mb4-tFhuA sensor into a lipid bilayer. When added to the *cis* compartment at nanomolar concentrations, WDR5 produced frequent current blockades (**Fig. 3a**; **Supplementary Fig. S11**) between O_on_ open substate and O_off_ partly closed substate with a normalized current amplitude (14 ± 1)% (**Fig. 3bc**). Again, this kind of current blockades was not noted when an unmodified tFhuA was exposed to WDR5 added to the *cis* side (**Supplementary Fig. S12**) or when Mb4-tFhuA was subjected to WDR5 added to the *trans* side (**Supplementary Fig. S13**). These findings suggest that specific WDR5-Mb4 interactions bring about WDR5-induced current blockades. WDR5-released (O_on_) and WDR5-captured (O_off_) events also followed a single-exponential distribution (**Fig. 3de**). In addition, the frequency of WDR5-captured events was proportional to its concentration, [WDR5] (**Fig. 3f**), whereas their duration was independent of [WDR5] (**Fig. 3e, Fig. 3g**; **Supplementary Tables S6-S7**). Using linear fits of the functions *f*([WDR5]) and *k*_off_([WDR5]), we obtained a *k*_on_ value (mean ± s.e.m.) of (0.83 ± 0.01) × 10^8^ M^-1^s^-1^and a *k*_off_ value (mean ± s.e.m.) of 72.4 ± 3.7 s^-1^, resulting a *K*_D_ of 872 ± 45 nM (**Supplementary Table S8**). It should be noted the kinetics of WDR5-Mb4 interactions undergo fast association and dissociation rates, which were also noted with hSUMO1-FN3SUMO interactions.

**Figure 3.**
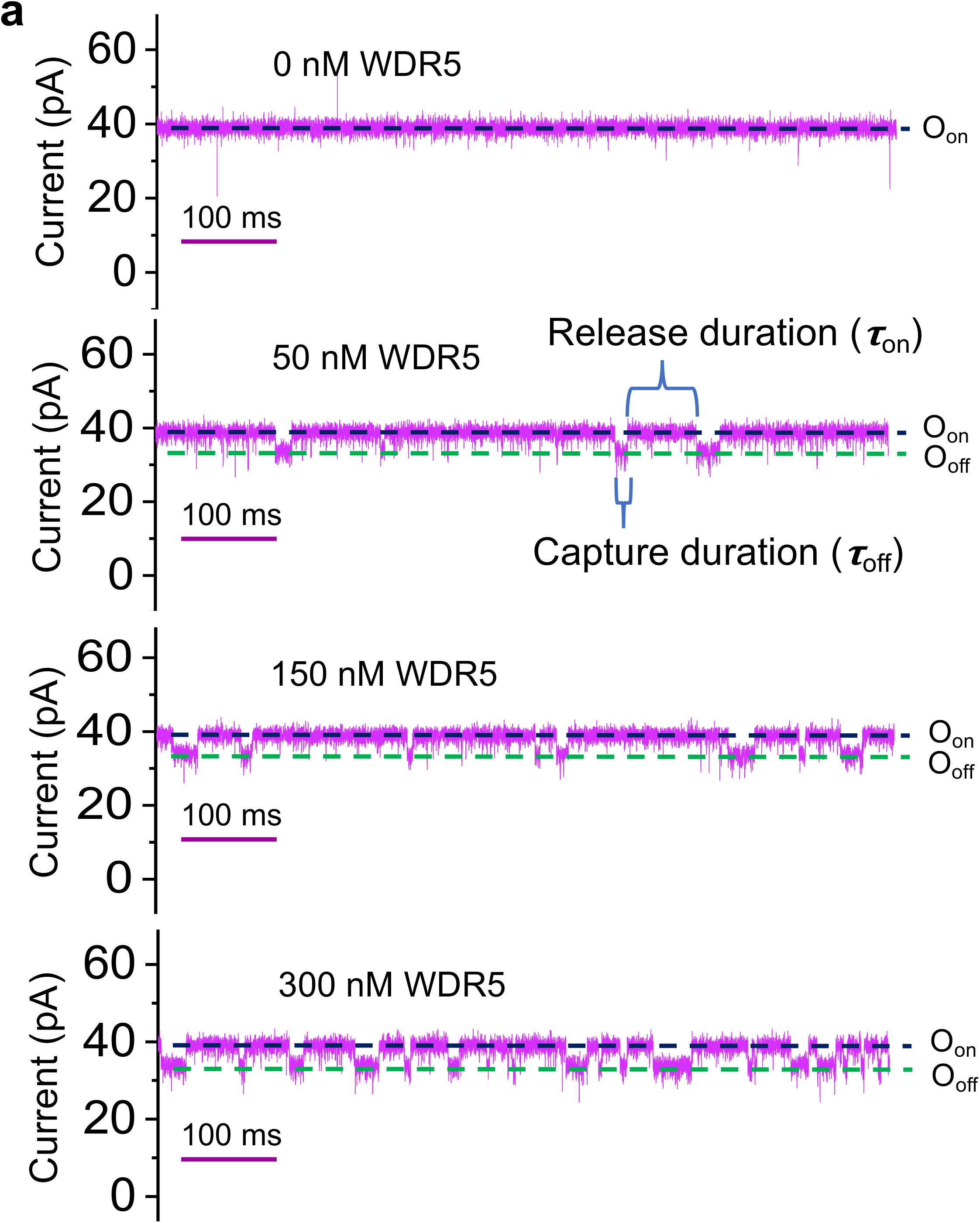
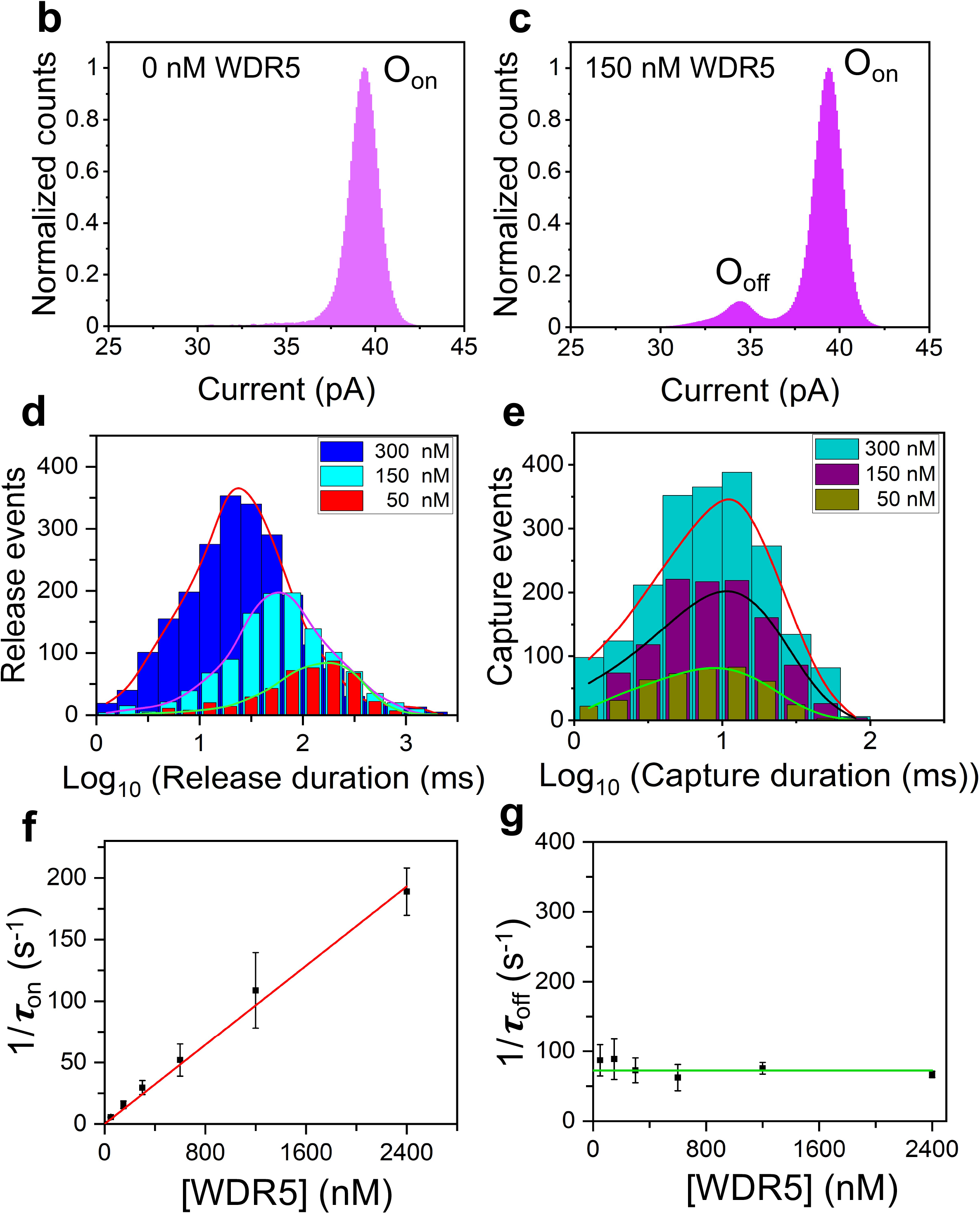
Single-molecule sensing of WDR5. **(a)** Representative single-channel electrical traces of Mb4-tFhuA in the presence of 0, 50, 150, and 300 nM WDR5. O_on_ and O_off_ are the WDR5-released and WDR5-captured substates, respectively. These single-channel electrical signatures were replicated in n=3 independent experiments. The other conditions were the same as those in **Fig. 2. (b)** An all-point current histogram of the O_on_ substate of Mb4-tFhuA. The current (mean ± s.e.m.) corresponding to the O_on_ substate was 39.5 ± 0.1 pA. **(c)** An all-point current histogram of the O_on_ and O_off_ substates of Mb4-tFhuA at 50 nM WDR5. The current (mean ± s.e.m.) corresponding to the O_off_ substate was 34.5 ± 0.1 pA. **(d)** Semilogarithmic histograms of the WDR5-released durations (*τ*_on_) at various WDR5 concentrations, [WDR5]. *τ*_on_ (mean ± s.e.m.) were 178 ± 6 ms (number of events: *N* = 466), 65 ± 7 ms (*N* = 1175), and 34 ± 4 ms (*N* = 2235) at [WDR5] values of 50 nM, 150 nM and 300 nM, respectively. **(e)** Semilogarithmic histograms of the WDR5-captured durations (*τ*_off_) at various [WDR5] values. *τ*_off_ (mean ± s.e.m.) were 12 ± 2 ms (*N* = 441 events), 10 ± 3 ms (*N* = 1127), and 14 ± 2 ms (*N* = 2034) at [WDR5] values of 50 nM, 150 nM and 300 nM, respectively. **(f)** Plot illustrating the dependence of the event frequency in the form of 1/*τ*_on_on [WDR5]. **(g)** Plot illustrating the dependence of 1/*τ*_off_on [WDR5]. The horizontal line is an average fit of the (1/*τ*_off_) data points. Data points in panels (f) and (g) represent mean ± s.d. obtained from *n* = 3 different experiments.

### An orthogonal method proves the rapid association and dissociation kinetics of WDR5-Mb4 interactions

To validate the fast kinetics recorded with the Mb4-tFhuA sensor, we performed additional measurements using biolayer interferometry (BLI).^50^ Mb4-tFhuA-containing micelles were immobilized onto the BLI sensor surface via a cysteine sulfhydryl engineered on the external L4 loop of tFhuA for biotin-streptavidin chemistry (**Methods**; **Supplementary Fig. 14a**). Hence, this experimental design mimics in some respect that of a sensing measurement with an Mb4-tFhuA sensor reconstituted into a lipid bilayer. WDR5 was added to different wells at increased concentrations. The association phases were recorded in real-time by placing the BLI sensors in WDR5-containing wells (**Supplementary Fig. 14b**). The dissociation phases were then recorded by putting the same BLI sensors in WDR5-free wells.

However, the rates of these kinetics are beyond the time resolution of BLI. Nevertheless, BLI sensorgrams acquired at various WDR5 concentrations qualitatively confirm the rapid kinetics of association and dissociation of WDR5-Mb4 interactions noted with the Mb4-tFhuA sensor (**Supplementary Table S8**).

### Adnectin1-tFhuA sensor reveals bimodal protein recognition of EGFR

The ectodomain of EGFR is proteolytically released into the bloodstream, allowing this biomarker to be used for screening, diagnosis, and disease progression.^36^ Hence, we employed Adnectin-1 against the ectodomain of EGFR.^43^ Adnectin1-tFhuA exhibited some current noise at +40 mV (**Supplementary Fig. S4c**). However, its traces showed a relatively quiet signature at a lower transmembrane potential of +20 mV (**Fig. 4a**; **Supplementary Fig. S15**). Interestingly, when EGFR was added to the *cis* side of the bilayer containing the Adnectin1-tFhuA sensor, reversible current blockades were observed in a broad temporal range and with various current amplitudes. In contrast, we noted only low-amplitude and brief current spikes when EGFR was added to the *cis* side of the bilayer containing tFhuA alone (**Supplementary Fig. S16**). A two-peak distribution was found for the current amplitudes of individual EGFR-captured events (**Fig. 4b**).

**Figure 4.**
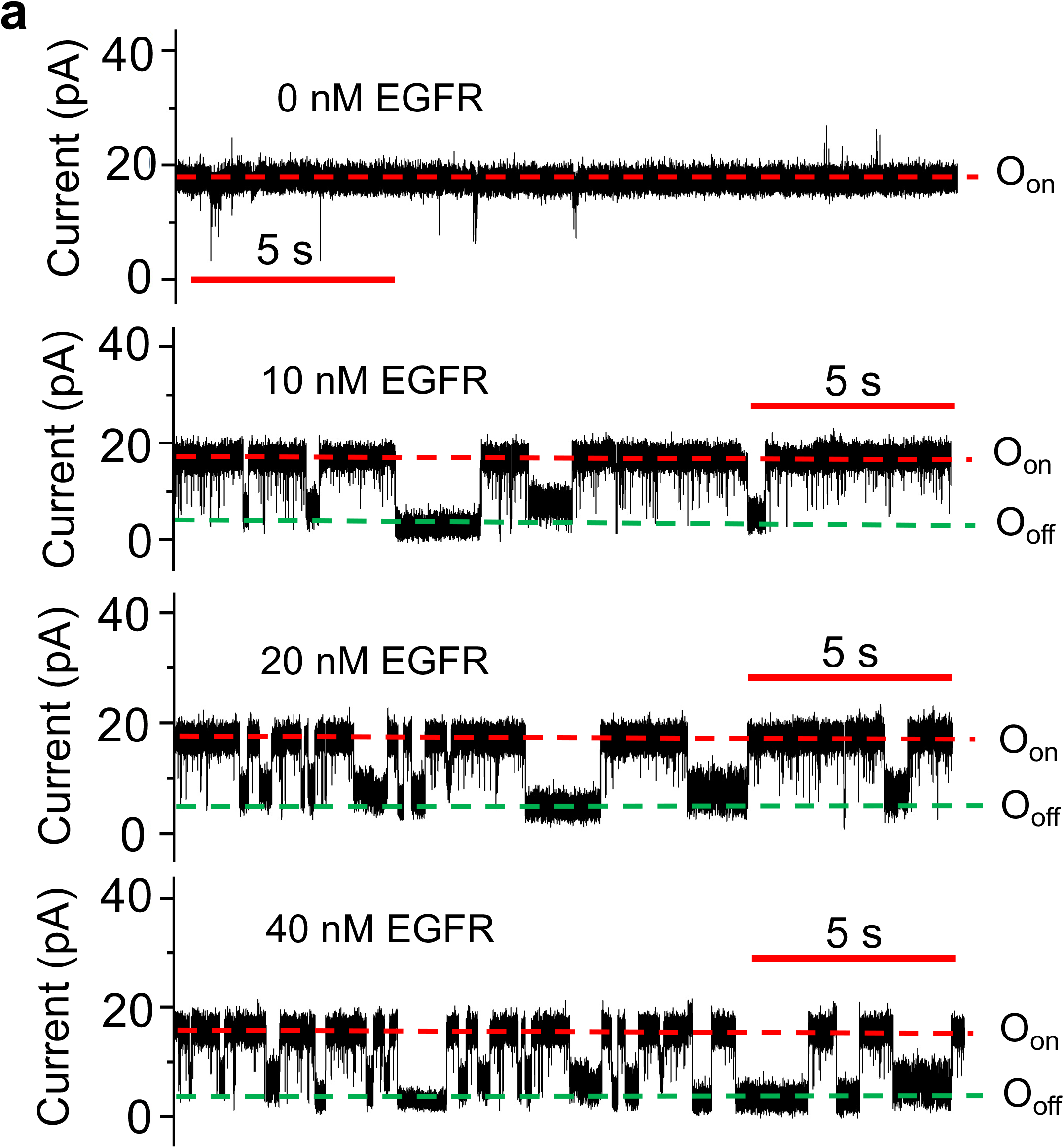
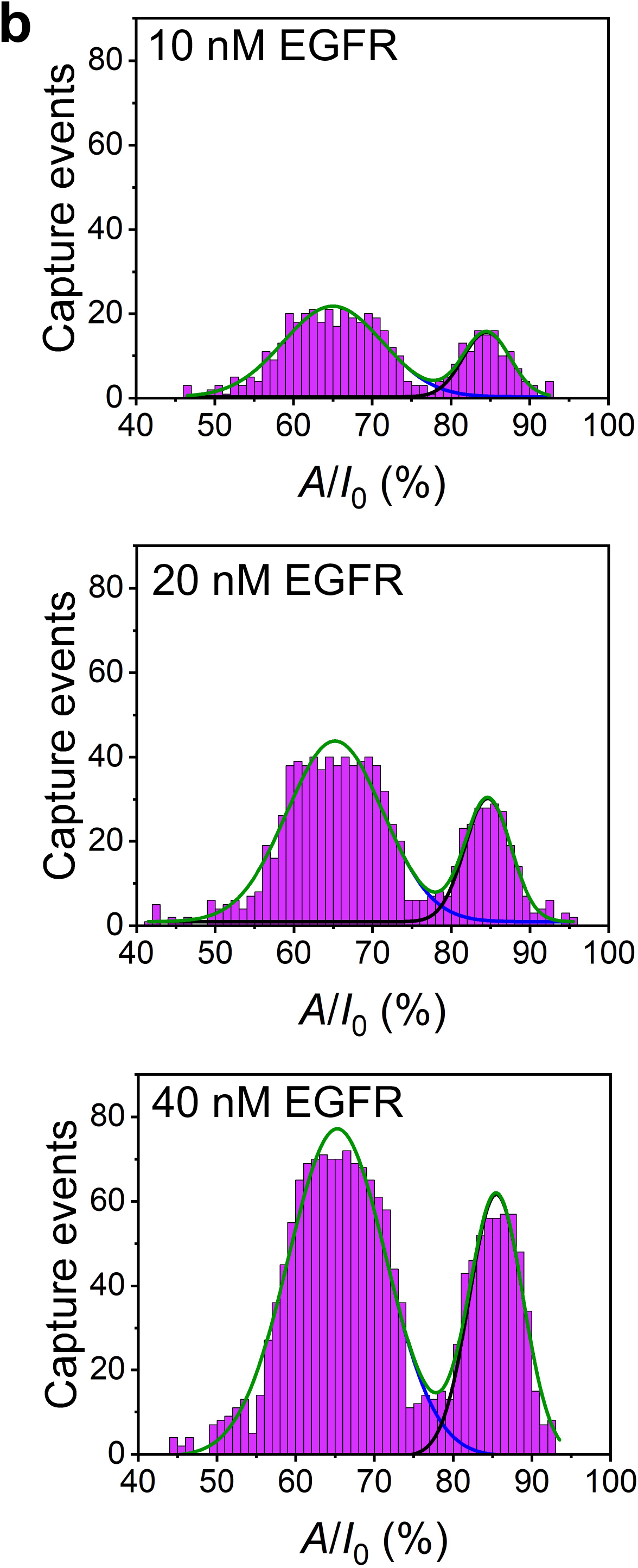
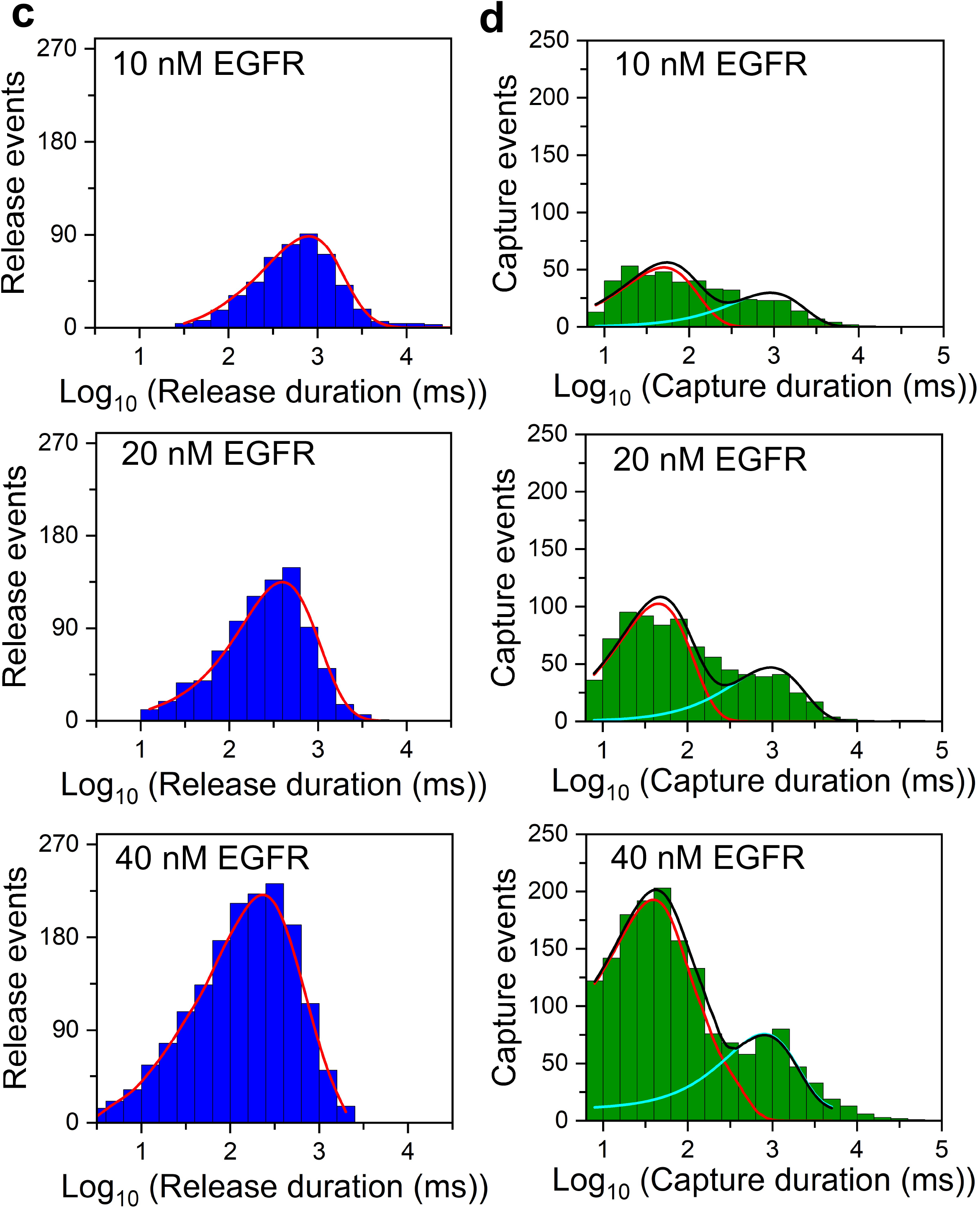
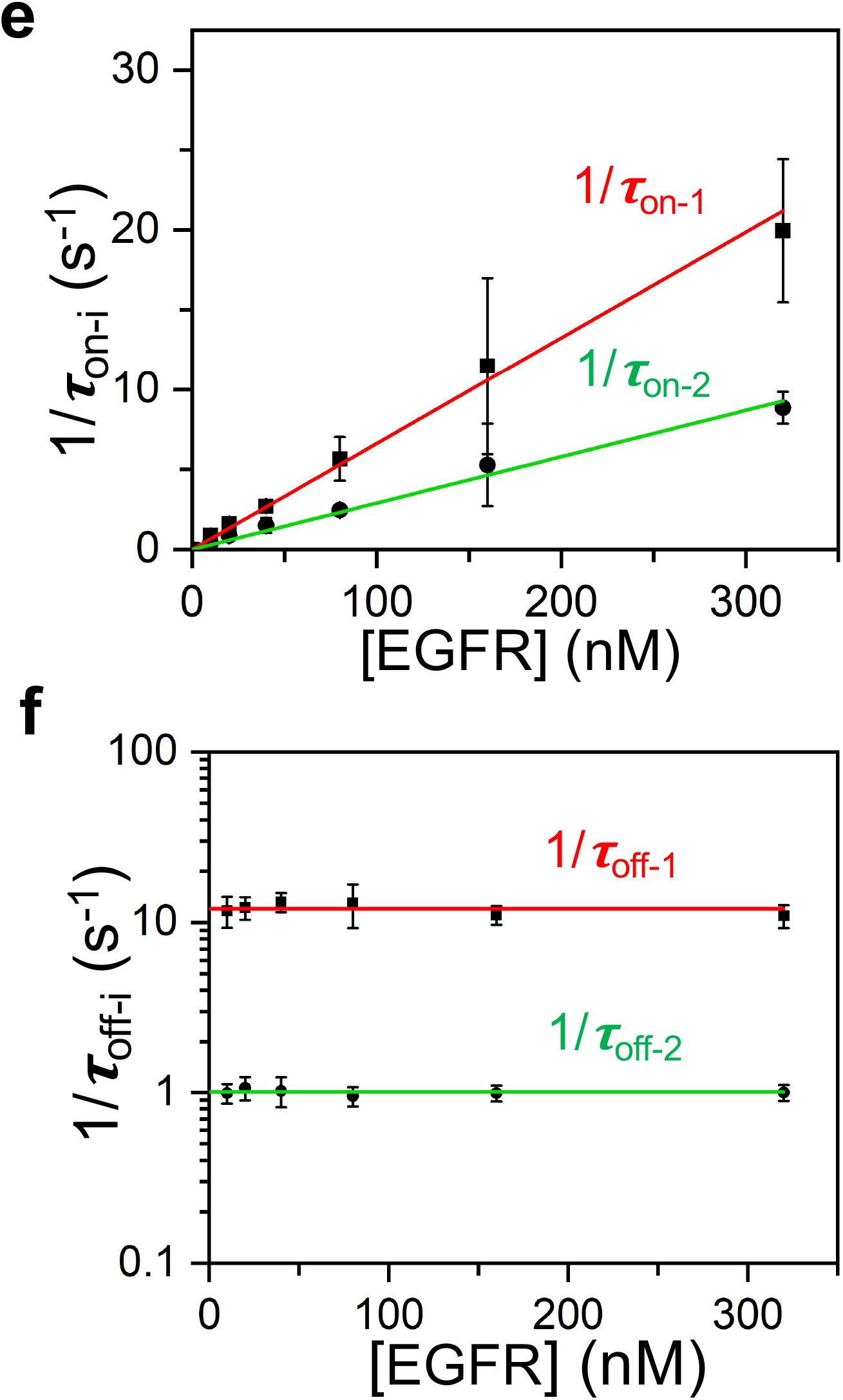
EGFR exhibits a bimodal protein recognition with Adnectin1. **(a)** Representative single-channel electrical traces of Adnectin1-tFhuA in the presence of 0, 10, 20, and 40 nM EGFR. O_on_ and O_off_ are the EGFR-released and EGFR-captured substates, respectively. These single-channel electrical signatures were replicated in n = 3 independent experiments. The applied transmembrane potential was +20 mV. Single-channel electrical traces were further low-pass filtered at 2 kHz using an 8-pole Bessel filter. **(b)** Event histograms of the normalized current blockades *A*/*I*_0_, where *A* and *I*_0_ are the current amplitude of individual blockades and the current amplitude of the O^on^ substate, respectively. The cumulative fits are marked in green. The blue and black curves indicate fits of low- and large-amplitude current blockades, respectively. For 10 nM EGFR, these values (mean ± s.e.m.) were (65.0 ± 0.3)% and (84.5 ± 0.3)%, respectively (number of events, *N* = 467). For 20 nM EGFR, they were (65.2 ± 0.2)% and (85.1 ± 0.2)%, respectively (*N* = 924). For 40 nM EGFR, they were (65.3 ± 0.2)% and (85.5 ± 0.2)%, respectively (*N* = 1711). **(c)** Semilogarithmic histograms of the EGFR-released durations (*τ*_on_) at various EGFR concentrations, [EGFR]. *τ*_on_ (mean ± s.e.m.) were 0.78 ± 0.04 s (number of events: *N* = 491), 0.42 ± 0.03 s (*N* = 843), and 0.25 ± 0.02 s (*N* = 1641) at [EGFR] values of 10 nM, 20 nM and 40 nM, respectively. **(d)** Semilogarithmic EGFR-captured durations (*τ*_off_) at various [EGFR] concentrations. The cumulative fits are marked in black. The red and cyan curves indicate fits for short- and long-lived EGFR captures, respectively. For 10 nM EGFR, they (mean ± s.e.m.) were 0.072 ± 0.011 s and 1.2 ± 0.1 s, respectively (number of events: *N* = 441). For 20 nM EGFR, they were 0.069 ± 0.007 s and 0.96 ± 0.09 s, respectively (*N* = 806). For 40 nM EGFR, they were 0.066 ± 0.006 s and 0.81 ± 0.11 s, respectively (*N* = 1598). **(e)** Plot illustrating the dependence of the event frequencies in the form of 1/*τ*_on-i_on [EGFR]. *τ*_on-1_and *τ*_on-2_ are the EGFR-released durations between the short- and long-lived EGFR captures, respectively. The slopes of the linear fits of 1/*τ*_on-i_ versus [EGFR] are the association rate constants, *k*_on-i_, of Adnectin1-EGFR interactions because *k*_on-i_ = 1/(*τ*_on-i_[EGFR]). **(f)** Plot illustrating the dependence of 1/*τ*_off-i_ on [EGFR]. The red and green horizontal lines are average fits of the (1/*τ*_off-1_) and (1/*τ*_off-2_) data points, respectively. Here, i=1 and i=2 are subscripts corresponding to the short- and long-lived EGFR captures, respectively. Data points in panels (e) and (f) represent mean ± s.d. obtained from *n* = 3 different experiments.

For example, at 40 nM EGFR, the normalized current blockades of the two peaks were (65.0 ±2.1)% and (86.1 ± 1.6)% with the probabilities of 0.72 ± 0.02 and 0.28 ± 0.02, respectively. Furthermore, the relative position and probability of these peaks were independent of EGFR concentration, [EGFR] (**Supplementary Table S9**).

EGFR-released (*τ*_on_) and EGFR-captured (*τ*_off_) durations followed single-peak and double-peak event distributions (**Fig. 4cd; Supplementary Tables S10-S13**), respectively, as judged by the maximum likelihood method^47, 51^ and logarithm likelihood ratio (LLR) tests.^48, 49^ Hence, our statistical analyses revealed two subpopulations of binding events, the short-lived and long-lived EGFR-captured events, whose durations were *τ* _off-1_ = ∼80 ms and *τ* _off-2_ = ∼1 s, respectively.

Interestingly, the probabilities of short-lived EGFR capture durations, *P*_1_, were close to those of low-amplitude current blockades (**Supplementary Tables S9-S10**). This outcome suggests two distinct mechanisms of binding of EGFR to Adnectin1, which correlate with the extent of the normalized current amplitude of EGFR-captured events and their duration. The event frequencies of short-lived and long-lived EGFR-captured events, in the form of 1/*τ*_on-1_ and 1/*τ*_on-2_, respectively, were linearly dependent on the EGFR concentration, [EGFR] (**Fig. 4e**). Here, *τ*_on-1_ and *τ*_on-2_ are the release (e.g., interevent) durations corresponding to the short-lived and long-lived current blockades, respectively (**Supplementary Table S12**). Again, the dissociation constants of the short-lived (*k*_off-1_) and long-lived (*k*_off-2_) current blockades were independent of [EGFR] (**Fig. 4f**; **Supplementary Table S14**). We interpret that these blockades are produced by specific bindings of EGFR to Adnectin1. We obtained the association rate constants, *k*_on-1_ and *k*_on-2_ (mean ± s.e.m.), of (6.62 ± 0.21) × 10^7^ M^-1^s^-1^ and (2.89 ± 0.10) × 10^7^ M^-1^s^-1^, respectively. The dissociation rate constants, *k*_off-1_ and *k*_off-1_ (mean ± s.e.m.), were 12.0 ± 0.4 s^-1^ and 1.01 ± 0.01 s^-1^, respectively (**Supplementary Table S15**). These values yield the equilibrium dissociation constants of the short-lived and long-lived current blockades, *K*_D-1_ and *K*_D-2_ (mean ± s.e.m.), of 181 ± 8 nM and 34 ± 2 nM, respectively.

The EGFR structure in the EGFR/EGF complex (1NQL.pdb)^52^ is similar to that of EGFR in the EGFR-Adnectin1 complex (3QWQ.pdb).^43^ It is believed to be an inactive form of the receptor (**Supplementary Fig. S17ab**).^43, 52^ Adnectin1 and EGF bind to the EGFR domain D-I with a highly overlapping binding surface (**Supplementary Fig. S17cd**). It is well established that EGFR is a remarkably adaptable molecule where the domains D-I and D-III are relatively rigid, whereas the domains D-II and D-IV can adopt multiple conformations that place domain D-III differently in relation to domain D-I.^52, 53^ We speculate that such distinct conformers of a flexible EGFR may likely be responsible for the bimodal protein recognition of EGFR by Adnectin1. The extended time bandwidth of our measurements facilitated the detection and quantification of conformational binding substates of the EGFR-Adnectin1 complex that are hidden in ensemble or low-resolution single-molecule measurements.^54^ Earlier studies using the resistive-pulse technique have also reported multimodal conformational transitions in the case of the dihydrofolate reductase (DHFR) enzyme.^55^

### Are there interconversion transitions between the capture substates?

Next, we asked whether these reversible current transitions may also involve transitions between the two EGFR-captured substates. Hence, a related question is whether a kinetic model including interconversion transitions between these EGFR-captured substates would more accurately reflect experimentally determined rate constants. An interconversion-dependent kinetic model was developed, encompassing two supplementary rate constants between EGFR-captured substates, *k*_12_ and *k*_21_ (**Supplementary Table S16, Fig. S18**). At a confidence level of *C* > 0.95, we found that fits to an interconversion-dependent kinetic model were not statistically superior over those corresponding to an interconversion-independent kinetic model, as indicated by the LLR test. Finally, to test the reactivity crosscheck of our sensors, we recorded electrical traces of Adnectin1-tFhuA in the presence of either hSUMO1 (**Supplementary Fig. S19**) or WDR5 (**Supplementary Fig. S20**). In both cases, very short-lived and low-amplitude current blockades were noted. These blockades resemble those typically found in the case of nonspecific interactions of folded proteins with the *cis* opening of tFhuA (**Supplementary Figs. S9, S12, S16**). This finding proves that the Adnectin1-tFhuA sensor is highly specific to EGFR.

### Single-molecule detection of a protein biomarker in a biofluid

We challenged this sensor in the presence of 5% (v/v) fetal bovine serum (FBS) to examine the stability of this system in a harsh environment and the ability to distinguish analyte-captured events from other nonspecific transitions of the solution constituents. The serum threshold for the soluble EGFR ectodomain level is 45 ng/ml (∼112 nM). The tumor state can be evaluated at EGFR levels significantly exceeding this threshold.^56, 57^ **Fig. 5a** and **Fig. 5b** show a representative signature of Adnectin1-tFhuA without and with 20 nM EGFR, respectively. However, the addition of 5% (v/v) FBS decorated the standard signature of EGFR-captured events with brief current spikes in the low-millisecond range (**Fig. 5c**). An analysis of the power spectral density (PSD) of current fluctuations revealed a transition from white noise in the absence of FBS to 1/*f* flicker noise in the presence of FBS (**Fig. 5d**). This outcome suggests low-frequency equilibrium fluctuations in the local mobility and density of charges at the nanopore tip in the presence of FBS.^58^ The brief FBS-induced current fluctuations had a lower current amplitude around the open O_on_ substate (**Fig. 5e-g**), indicating that these may result from trafficking moieties of serum constituents at the *cis* opening of Adnectin1-tFhuA (**Fig. 1**). An extensive statistical analysis of the current blockades corresponding to the O_off-1_ and O_off-2_ levels confirmed the presence of two EGFR-captured event types in the presence of FBS (**Fig. 5h-k; Supplementary Tables S17-S18**). No statistically significant impact of FBS was noted on the *k*_off-1_ and *k*_off-2_, but small changes, within the same order of magnitude, on the *k*_on-1_ and *k*_on-2_ (**Supplementary Table S19**). These changes may result from the interference of serum constituents with the binding interfaces of EGFR and Adnectin1. The mean duration of long-lived EGFR-induced current blockades was *τ*_off-2_ = 0.93 ±0.14 s, much longer than the brief millisecond-timescale FBS-induced closures. Under these conditions, we determined a corresponding *τ*_on-2_of 1.65 ± 0.50 s. Using a *k*_on-2_ of (2.89 ± 0.17) × 10^7^ M^-1^s^-1^ in the absence of FBS, we can evaluate the EGFR concentration in the serum sample, [EGFR], using the equation [EGFR] = 1/(*τ*_on-2_*k*_on-2_). Employing these values, we determined an [EGFR] of 22.2 ± 5.9 nM in the FBS-containing sample, near the actual concentration of 20 nM.

**Figure 5.**
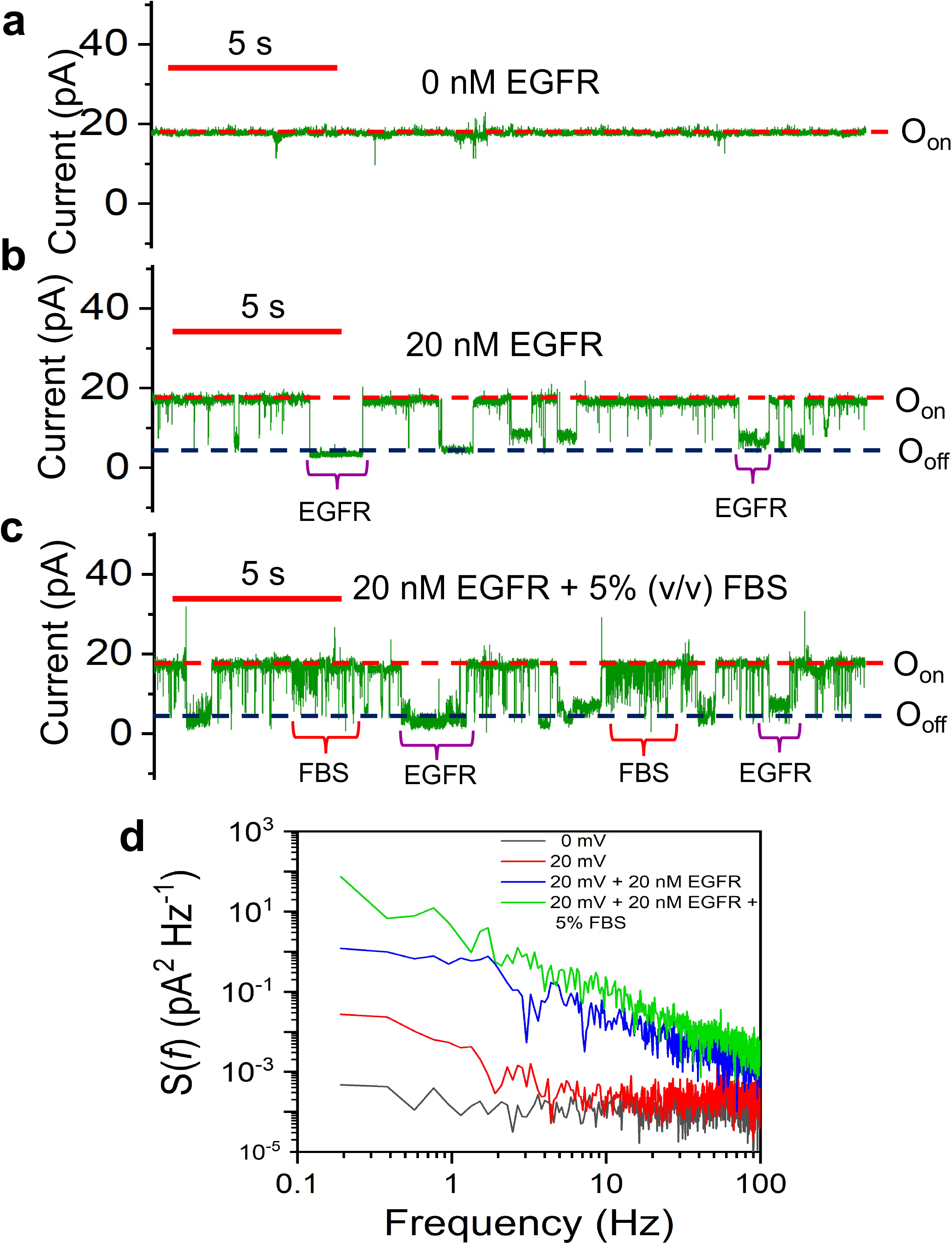
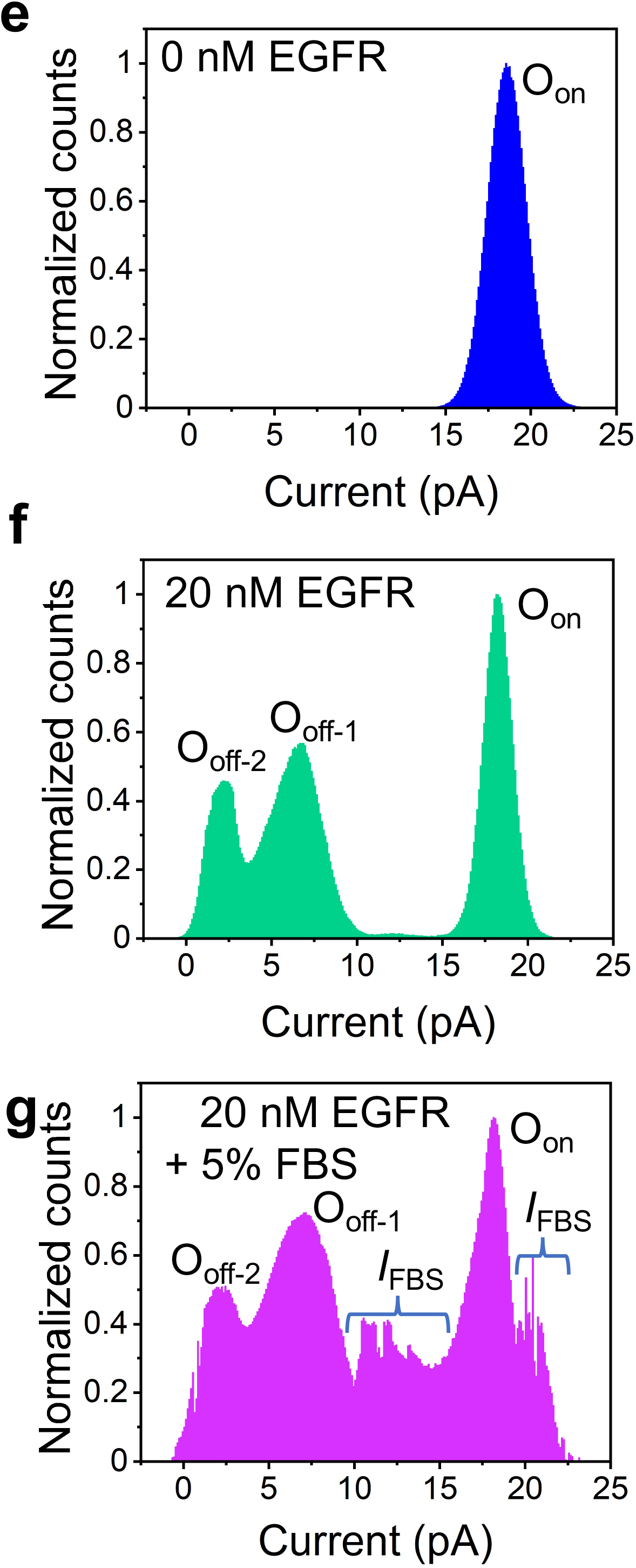
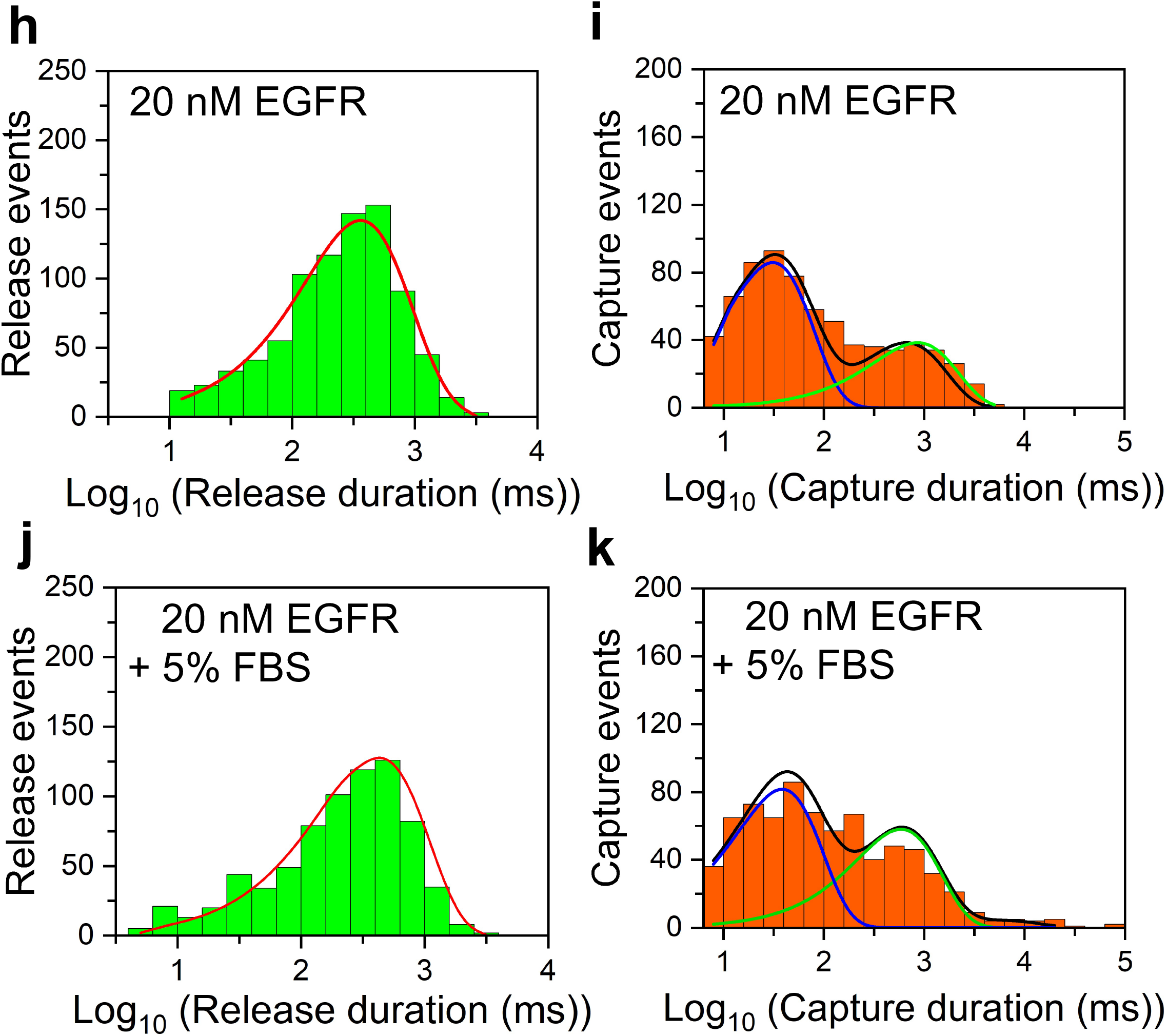
Single-molecule detection and quantification of EGFR in a heterogeneous solution. **(a)** A representative single-channel electrical trace of Adnectin1-tFhuA. **(b)** The trace in (a) in the presence of 20 nM EGFR. **(c)** The trace in (b) in the presence of 5% (v/v) FBS. The transmembrane potential was +20 mV. This subset of single-channel electrical signatures was replicated in *n* = 3 independent experiments. Single-channel electrical traces were further low-pass filtered at 500 Hz using an 8-pole Bessel filter. **(d)** Power spectral density (PSD) of current noise of traces illustrated in panels (a) - (c). Each spectrum represents an average of three independent traces. **(e)** A representative all-point current histogram of the O_on_ substate of Adnectin1-tFhuA. The current amplitude (mean ± s.e.m.) of the O_on_ substate was 18.2 ± 0.1 pA. **(f)** An all-point current histogram of the O_on_ and O^off^ substates of Adnectin1-tFhuA at 20 nM EGFR. The currents (mean ± s.e.m.) of the O_off-1_ and O_off-2_ substates were 6.4 ± 0.1 pA and 2.3 ±0.1 pA, respectively. **(g)** An all-point current histogram of the O_on_ and O_off_ substates of Adnectin1-tFhuA at 20 nM EGFR and in the presence of 5% fetal bovine serum (FBS). This plot reveals the residual signal produced by the FBS constituents (*I*_FBS_). **(h)** A semilogarithmic histogram of the EGFR-released durations (*τ*_on_) at 20 nM EGFR. *τ*_on_ (mean ± s.e.m.) was 0.40 ± 0.03 s (number of events: *N* = 844). **(i)** A semilogarithmic histogram of the EGFR-captured durations (*τ*_off_) at 20 nM EGFR. *τ*_off-1_ and *τ*_off-2_(mean ± s.e.m.) were 0.044 ± 0.015 s and 0.982 ± 0.049 s, respectively (number of events: *N* =734). **(j)** A semilogarithmic histogram of the EGFR-released durations (*τ*_on_) at 20 nM EGFR and in the presence of 5% FBS. *τ*_on_ (mean ± s.e.m.) was 0.607 ± 0.051 s (*N* = 738). **(k)** A semilogarithmic histogram of the EGFR-captured durations (*τ*_off_) at 20 nM EGFR and in the presence of 5% FBS. *τ*_off-1_and *τ*_off-2_(mean ± s.e.m.) were 0.036 ± 0.013 s and 0.806 ± 0.078 s, respectively (*N* = 694). The cumulative fits are marked in black in panels (i) and (k). The blue and green curves indicate fits for the short- and long-lived EGFR captures, respectively.

### Distinct outcomes with monobody-based sensors

In this study, we provide a detailed signature analysis of single-molecule protein detection of three analytes using three nanopore sensors that share a modular architecture but differ by their binding surface (**Supplementary Figs. S21-S23**). Fortuitously, all monobodies significantly block the ionic flow through tFhuA, allowing direct electrical detection of analyte bindings without needing any peptide tag.^21, 23^ Because the protein analytes and their complexes with the specific monobodies drastically vary in size, charge, and structural complexity, distinct current blockades are noted in each case (**Supplementary Fig. S24**). For example, WDR5 interacts with a distal FG loop of Mb4 and away from the tFhuA pore opening (**Supplementary Fig. S22**), suggesting a modest current blockade made by the WDR5-Mb4 complex. In accord with this expectation, we note low-amplitude current blockades produced by WDR5-captured events (**Supplementary Table S20**). In contrast, the conformation complexity and structural properties of the hSUMO1-FN3SUMO and EGFR-Adnectin1 complexes at the tip of tFhuA indicated a potentially large ionic flow block, as also found by electrical recordings. In addition, we probed distinct single-molecule kinetic signatures of each analyte without the steric restrictions of the nanopore confinement (**Supplementary Fig. S25**). These unique characteristics of protein detections using externally engineered complex binding interfaces culminated with the discovery of a bimodal protein recognition of EGFR.

### Validation of the monobody-based sensors

Next, we examined the binding affinity of detergent-refolded sensors with their cognate analytes using steady-state fluorescence polarization (FP) anisotropy. If the labeled protein analyte interacts with the corresponding monobody-containing sensor, its tumbling rate (e.g., the coefficient of rotational diffusion) decreases, increasing the FP anisotropy. In accord with our expectation, the FP anisotropy substantially increased at elevated sensor concentrations (**Supplementary Fig. S26**). On the contrary, the presence of tFhuA at increased concentrations did not alter the FP anisotropy, confirming no interaction between labeled proteins and sensor-containing detergent micelles. The calculated *K*_D_ values of hSUMO1 and WDR5 with their respective nanopore sensors were 186 ± 16 nM and 455 ± 59 nM, respectively, which agree with the outcomes of single-channel electrical recordings of these sensors (**Supplementary Tables S5, S8**). EGFR is unsuitable for this assay because of its large molecular weight, so its tumbling rate is longer than the fluorescence lifetime of most fluorophores. However, the *K*_D_determined for the long-lived EGFR-captured events using Adnectin1-tFhuA sensor is in accord with a previously reported study (**Supplementary Table S15**).^43^ It should be mentioned that restraining one binding partner to a surface can decrease the affinity to one order of magnitude.^50^ Hence, this explains a slightly weaker binding interaction with the immobilized nanopore sensor on a lipid bilayer than that value measured in solution by steady-state FP spectroscopy.

### Advantages of these nanopore sensors and their implications in nanobiotechnology

In this study, we engineered a new class of nanopore sensors made of a single-polypeptide unit that features a selective protein binder adaptable with atomic precision. The monomeric nature of these sensors circumvents the necessity of tedious purification steps of the assembly reaction, otherwise required for multimeric nanopores. The overall architecture of the sensors can be maintained while changing the interaction interface of the antibody-mimetic binder. This way, such an approach substantially extends the applications of these sensing elements for numerous protein biomarkers. This critical benefit is facilitated by the genetically encoded nature of these sensors so that they can create combinatorial libraries of tethered binders. For instance, the loops of monobodies are analogous to the complementarity determining regions (CDRs) of antibodies. One significant advantage of monobodies is their ability to interact with challenging binding surfaces that are not generally exposed to the CDRs of antibodies.^59^ In addition, there is no fundamental limitation in replacing the monobody with another small protein binder, such as a nanobody or an affibody.^60-63^ Furthermore, the main benefits of using antibody-mimetic proteins include strong binding affinities with different epitopes, straightforward expression and purification procedures, lack of disulfide bonds, and high thermodynamic stability.^59, 64^

Notably, our method has the potential to detect and characterize functionally distinct subpopulations of specific binding events in a challenging biofluid. We probe the complexity and heterogeneity of protein recognition events without requiring any additional exogenous tag or peptide tail. In addition, this sensor formulation includes a system that precludes the occurrence of nonspecific binding events or protein inactivation at the liquid-surface interface, as in the case of surface immobilization-based sensors. Our proposed approach shows prospects for discovering rare and short-lived binding events, which are unlikely to be detectable by prevailing technologies. In extreme conditions of unusually high *k*_on_, such as those in the range of 10^7^ - 10^9^ M^-1^s^-1^,^19^ we show that our method can be utilized to measure such values (e.g., for hSUMO1 and WDR5). In nanopore-based sensing, the *k*_off_ can be recorded up to a value of ∼10^5^ s^-1^.^11^ Hence, these sensors can operate at clinically relevant concentration ranges of protein biomarkers and with an extended time bandwidth. In this process, the analyte-induced events are unambiguously distinguished from other nonspecific current blockades of biofluid constituents. With further developments, these sensors can be integrated with high-throughput technologies for biomarker profiling in biomedical diagnostics.

## Methods

### Computational grafting of monobodies onto tFhuA

For the structural prediction of nanopore sensors, the amino acid sequence of each monobody (FN3, FN3SUMO, Mb4 and Adnectin1) was inserted at the N-terminus of tFhuA via a (GGS)_2_ peptide tether. 3D structural models of the nanopore sensors were generated *in silico* using AlphaFold2.^44, 45^ All parameters were kept the same for all nanopore sensors. The predicted structures of sensors were confirmed by comparisons with individual structures of FhuA and monobodies.

### Synthetic gene construction

Three derivatives of wild-type fibronectin type-III (FN3) were used to develop these sensors. The cDNA sequences of these *fn3* genes, namely *fn3sumo, mb4*, and *adnectin1*, were fused to the 5’ end of the *tfhua* gene via a (GGS)_2_-encoding linker by a restriction-free cloning method.^65^ The cDNA sequences of Mb4 and Adnectin1 were synthesized by Eurofins Genomics (Louisville, KY) and Integrated DNA Technologies (IDT, Coralville, Iowa), respectively. The construction of the *fn3sumo* gene was made based on ySMB9.^41^ The cDNA sequence of all three fibronectin derivatives was first amplified using Q5 high-fidelity DNA polymerase (New England BioLabs, Ipswich, MA) from their respective template DNA. PCR products were separated on 1% agarose gel and purified using a Gel extraction kit (Promega, CA). Sequences of forward and reverse primers are listed in **Supplementary Table S21**. Amplified products of *fn3sumo* and *mb4* genes were then fused to the 5’ end of *tfhua* cloned in pPR-IBA1 plasmid (IBA, Goettingen, Germany). *adnectin1* was joined at the 5’ end of *tfhua* in pET28a (EMD Millipore, Burlington, MA). The pET28-tFhuA plasmid was constructed by inserting the gene between *Bam*HI and *Xho*I restriction sites after amplification with forward and reverse primers of tFhuA **(Supplementary Table S21)**. All the gene sequences were verified by sequencing (MCLab, San Francisco, CA). The pET11a-hSUMO1 was kindly provided by Fauke Mechior (Addgene plasmid #53138).

### Protein expression and purification

For the expression of FN3SUMO-tFhuA, Mb4-tFhuA, and Adnectin1-tFhuA, the plasmids mentioned above were transformed into *E. coli* BL21(DE3) cells. These monobody-containing protein nanopores were purified as previously described.^21, 22^ The protein purity was validated by SDS-PAGE analysis (**Supplementary Fig. S1)**. In the case of hSUMO1, BL21(DE3) cells were transformed with pET11a-hSUMO1 and grown in Luria– Bertani (LB) medium at 37°C until OD_600_ reached a value of ∼0.5. Then, the temperature was changed to 20°C. Expression was initiated by inducing the cells with 250 µM IPTG. After induction, the cells were cultured for ∼18 h at 20°C. Cells were then centrifuged at 3,700*g* for 30 min at 4°C, followed by their resuspension in 50 mM Tris-HCl, 50 mM NaCl, and pH 8.0. The lysozyme was added to the suspended cells and incubated on ice for 15 min, and cell lysis was accomplished using sonication (30 s on, 60 s off × 4 times). The cell lysate was centrifuged at 108,500*g* for 30 min at 4°C to separate the insoluble pellet and supernatant. The supernatant was collected and filtered using a 0.22 µm filter. The supernatant was loaded onto a Q-Sepharose column (Cytiva, Marlborough, MA), which was washed with 50 mM Tris-HCl, 50 mM NaCl, pH 8.0, and eluted with 50 mM Tris-HCl, 1 M NaCl, pH 8.0 in a gradient manner. The desired fractions were collected, dialyzed, and concentrated. Furthermore, the protein sample was loaded on an S75 gel-filtration column (GE Healthcare, Chicago, IL). Pure fractions were collected and dialyzed against 20 mM Tris-HCl, 150 mM NaCl, pH 8.0, and 0.5 mM TCEP overnight at 4°C. Purification of WDR5 was done as described previously.^21^

For the purification of the ectodomain of epidermal growth factor receptor (EGFR), Expi293F cells (Thermo Fischer Scientific) were seeded at 10^6^ cells/ml density in 1 L of Dynamis growth medium (Gibco) 24 h before the transfection and supplemented with Tryptone/Glucose. For the sake of simplicity, we name this EGFR throughout this article. The culture was transfected with 2 µg/mL of the pCMV_EGFR plasmid containing the signal peptide with 3.75 × polyethylenimine (PEI). Transfected cells were cultured for five days, and the protein was allowed to excrete from the cells. Five days post-transfection, the culture was pelleted, and the supernatant was filtered. The sample was loaded onto an immobilized metal-affinity column (1 mL, HIStrap HP column, GE Healthcare), which was washed with 50 mM sodium phosphate (NaPi) (pH 8.0), 300 mM NaCl, 20 mM imidazole. The protein was eluted using 50 mM NaPi (pH 8.0), 300 mM NaCl, and 500 mM imidazole. Peak fractions were collected and confirmed by SDS-PAGE (**Supplementary Fig. S27)**. Finally, the protein sample was concentrated and exchanged with phosphate buffer saline (PBS, pH 7.5) using a PD10 column (GE Healthcare) and stored at -80°C. The purity of all protein analytes was tested by SDS–PAGE analysis.

### Protein refolding

The purified FN3SUMO-tFhuA, Mb4-tFhuA, and Adnectin1-tFhuA were adjusted to a final concentration of ∼10 µM. Next, n-dodecyl-β-d-maltopyranoside (DDM) was added to denatured samples to a final concentration of 1% (w/v). The protein samples were immediately dialyzed against the buffer containing 200 mM KCl, 20 mM Tris-HCl, pH 8, at 4°C for 96 h. The dialysis solution was replaced at 24-h intervals. These refolded protein samples were centrifuged to eliminate any protein precipitations, and the supernatant was used as the running sample for single-channel electrical recordings. Protein concentrations were determined by their molar absorptivity at a wavelength of 280 nm.

### Single-channel electrical recordings

Electrical detection of protein ligands at single-molecule precision was conducted using planar lipid bilayers.^66^ The two halves of the chamber were divided by a 25 µm-thick Teflon septum (Goodfellow Corporation, Malvern, PA). A planar lipid bilayer was made of 1,2-diphytanoyl-*sn*-glycero-phosphatidylcholine (Avanti Polar Lipids, Alabaster, AL) across an ∼100 μm-diameter aperture of the Teflon septum. For all experiments, the buffer solution contained 300 mM KCl, 10 mM Tris-HCl, and pH 8.0. In addition, this buffer included 0, 0.5, and 1 mM TCEP in experiments with EGFR, hSUMO1, and WDR5, respectively. The nanopore protein samples (final concentration, 0.5-1.5 ng/µl) and analytes were added to the *cis* compartment, which was grounded. Single-channel electrical currents were acquired using an Axopatch 200B patch-clamp amplifier (Axon Instruments, Foster City, CA).

The applied transmembrane potential was +40 mV, unless otherwise stated. The electrical signal was sampled at 50 kHz using a low-noise acquisition system (Model Digidata 1440 A; Axon Instruments). A low-pass Bessel filter (Model 900; Frequency Devices, Ottawa, IL) was further employed for signal filtering at 10 kHz. For the data processing and analysis, the electrical traces were digitally filtered with a low-pass 8-pole Bessel filter at 3 kHz, unless otherwise stated. All single-channel electrical recordings were acquired at a temperature of 24 ± 1°C.

### EGFR detection in a heterogeneous solution

For the detection and quantification of EGFR in heterogeneous solutions, fetal bovine serum (FBS, Gibco, Thermo Fisher Scientific, Pittsburgh, PA) was used. FBS was sterilized through a syringe filter before being stored at −80°C. For single-channel recording, an aliquot was defrosted on ice and kept at room temperature before adding to the chamber. Single-channel electrical traces were recorded in the presence of FBS at a final concentration of 5% (v/v). These traces were filtered with a low-pass 8-pole Bessel filter at 500 Hz.

### Biolayer interferometry (BLI) assay using immobilized proteomicelles

These experiments were conducted using an Octet Red384 instrument (FortéBio, Fremont, CA) at 24°C. For BLI experiments, a site-specific insertion of cysteine at position 287 was achieved in the long L4 loop of Mb4-tFhuA by site-directed mutagenesis (Q5 mutagenesis kit, New England Biolabs). This cysteine-containing Mb4-tFhuA was expressed and purified as described above, except for the presence of a reducing agent. Cys287 was biotinylated using maleimide chemistry. A flexible (PEG)_11_ linker was used between the biotin and maleimide groups. This way, there was a satisfactory distance between Mb4 and the surface of the BLI sensor. The BLI running buffer contained 300 mM KCl, 20 mM Tris-HCl, 1 mM TCEP, 1% DDM, 1 mg/ml bovine serum albumin (BSA), pH 8.0. It was used to soak streptavidin (SA) sensors for 30 min. The 50 nM Mb4-tFhuA_Cys287-(PEG)_11_-Biotinyl was loaded onto the sensors for 2.5 min via biotin-streptavidin chemistry. By dipping the sensors in a protein-free solution for 6 minutes, the unattached Mb4-tFhuA_Cys287 was washed away. The association process was examined using various concentrations of WDR5, ranging from 1.5 µM to 6 µM. The BLI sensors were dipped in a WDR5-free running buffer to inspect the dissociation phase. For all WDR5 concentrations, the Mb4-tFhuA_Cys287-free BLI sensors were run in parallel as controls. The baseline and drift in the sensorgrams were subtracted using these controls. The FortéBio Octet data analysis software (FortéBio) was used for the sensorgram analysis.

### Steady-state fluorescence polarization (FP) measurements

hSUMO1 and WDR5 were labeled with fluorescein and rhodamine, respectively, at pH9.0 by primary amine chemistry. These labeled proteins were added to the well at a final concentration of 50 nM. Steady-state fluorescence polarization (FP) anisotropy assays were conducted in triplicate with an 18-point serial dilution of FN3SUMO-tFhuA, Mb4-tFhuA, or unmodified tFhuA, against a fixed concentration of labeled proteins on black 96-well plates. All steady-state FP measurements were recorded using a SpectraMax i3x plate reader (Molecular Devices, San Jose, CA) at 0 min and after a one-hour incubation at room temperature in the dark. The resulting dose-response data were averaged and fitted using logistic regression to obtain each interaction’s dissociation constant (*K*_D_).

### Statistical analysis

pClamp 10.7 (Axon Instruments) was used for the data acquisition and analysis. Capture and release events were collected using the single-channel event search in ClampFit 10.7 (Axon Instruments), and figures were prepared by Origin 9.7 (OriginLab, Northampton, MA). The probability distribution function (PDF) was generated using a kinetic rate matrix, and the kinetic rate constants were determined by fitting the data using the maximum likelihood method.^51^ To evaluate the results of multiple models and select the number of statistically significant peaks that are best matched to the data, a logarithm likelihood ratio (LLR) test was performed.^48, 49^ At a confidence number of C = 0.95, a single-exponential fit was the best model for the release and capture durations of hSUMO1 and WDR5. For EGFR, a two-exponential fit was the best model for the capture durations.

### Molecular graphics

All cartoons showing molecular graphics were prepared using PyMOL 2 (Version 2.4.0; Schrödinger, LLC) and Chimera X (Version 1.4; The University of California at San Francisco).

### Data availability

Data from this study is presented in the main text and **Supplementary Information** file. Other data that support the findings of this study are available from the corresponding author upon reasonable request.

### Code availability

pdb codes used in this article are listed in the **Supplementary Information** file, **Table S22**. All custom codes and mathematical algorithms used in this study are available from the corresponding author upon reasonable request.

## Acknowledgments

We thank our colleagues in the Movileanu and Loh laboratories and at Ichor Life Sciences laboratories for their comments on the manuscript and for their technical assistance during the very early stage of this project. We are grateful to Tom Duncan for his instrumental help in the initial phase of BLI experiments and Ali Imran for his gracious assistance with the Q-matrix analysis. This work was supported by the National Institute of General Medical Sciences of the U.S. National Institutes of Health grants GM088403 (to L.M.) and GM115762 (to S.N.L.).

## Author contributions

M.A., J.H.H., L.A.M., M.F.P., S.N.L., and L.M. designed research. M.A., J.H.H., M.F.P. performed research. M.A. and L.A.M. analyzed data. A.J.W., K.J.M., S.N.L., and L.M. supervised research and provided research materials and funding. M.A., J.H.H, and L.M. wrote the paper.

## Additional information

### Supplementary information

The online version contains supplementary materials available at http://doi.org/XXXXXXXX

