## Supplementary File Online for "A Generalizable Nanopore Sensor for Highly Specific Protein Detection at Single-Molecule Precision"

**Table of Contents of the Supplementary Information file.**


1. Structural and physical features of the three protein analytes inspected in this study (**Supplementary Table S1**).

2. SDS-PAGE gel analysis of the three monobody-containing sensors (**Supplementary Figure S1**).

3. Three-dimensional structure predictions of monobody-containing nanopore sensors (**Supplementary Figures S2-S4**).

4. Single-channel electrical signatures and unitary conductance values of the three monobody-containing sensors (**Supplementary Figures S5-S7**, **Table S2**).

5. hSUMO1-induced current blockades noted with the FN3SUMO-tFhuA sensor (**Supplementary Figure S8**).

6. Negative- and positive-control experiments with the FN3SUMO-tFhuA sensor (**Supplementary Figures S9-S10**).

7. Time, rate, and equilibrium constants of hSUMO1-FN3SUMO interactions (**Supplementary Tables S3-S5**).

8. WDR5-induced current blockades noted with the Mb4-tFhuA sensor (**Supplementary Figure S11**).

9. Negative- and positive-control experiments with the Mb4-tFhuA sensor (**Supplementary Figures S12-S13**).

10. Time, rate, and equilibrium constants of WDR5-Mb4 interactions (**Supplementary Tables S6-S8**).

11. Biolayer interferometry (BLI) measurements of WDR5−Mb4-tFhuA interactions (**Supplementary Figure S14**).

12. EGFR-induced current blockades noted with the Adnectin1-tFhuA sensor (**Supplementary Figure S15**).

13. Negative-control experiments with the Adnectin1-tFhuA sensor (**Supplementary Figure S16**).

14. Normalized current amplitudes of EGFR-produced blockades (**Supplementary Table S9**).

15. Time, rate, and equilibrium constants of EGFR-Adnectin1 interactions (**Supplementary Tables S10-S15**).

16. Structures of Adnectin1 and EGF in complex with EGFR (**Supplementary Figure S17**).

17. Interconversion-dependent and interconversion-independent kinetic models of EGFR-Adnectin1 interactions (**Supplementary Table S16, Figure S18**).

18. Positive-control experiments crosschecking the reactivity of the Adnectin1-tFhuA sensor for other protein analytes (**Supplementary Figures S19-S20**).

19. Time, rate, and equilibrium constants of EGFR−Adnectin1 interactions in the presence of mammalian serum (**Supplementary Tables S17-S19**).

20. Side and top views of the three monobody-based sensors in complexes with their cognate protein analytes (**Supplementary Figures S21-S23**).

21. The three monobody-containing sensors exhibit different electrical and kinetic signatures (**Supplementary Table S20 Figures S24-S25**).

22. Steady-state FP anisotropy curves of hSUMO1−FN3SUMO-tFhuA and WDR5−Mb4-tFhuA interactions in bulk phase (**Supplementary Figure S26**).

23. List of primers used in this study (**Supplementary Table S21**).

24. SDS-PAGE analysis of purified EGFR (**Supplementary Figure S27**).

25. PDB entries used for visualization and molecular graphics of all protein structures employed in this study (**Supplementary Table S22**).

26. Supplementary references.

***1. Structural and physical features of the three protein analytes inspected in this study.***

**<u>Supplementary Table S1.</u> Comparison of the size, charge, and structural complexity of three protein analytes.**

| Protein analytes* | Size (kDa) | Charge** | Structural organization |
|---|---|---|---|
| **hSUMO1** | 11.1 | -5.8 | 3 α helices and 4 β strands |
| **WDR5** | 36.5 | 1.9 | 7-bladded beta-propeller fold with a total of 28 β strands |
| **EGFR*** ** | 69.2 | -16.3 | Four distinct structural domains |

*hSUMO1, WDR5, and EGFR stand for human small ubiquitin-related modifier 1,[1-4] WD40 repeat protein 5,[5, 6] and epidermal growth factor receptor,[7] respectively.
**Charges of these protein analytes were calculated using protein calculator v3.4 by amino acid sequence at pH 8.0.
***This is the ectodomain (the extracellular domain) of EGFR.

***2. SDS-PAGE gel analysis of the three monobody-containing sensors.***

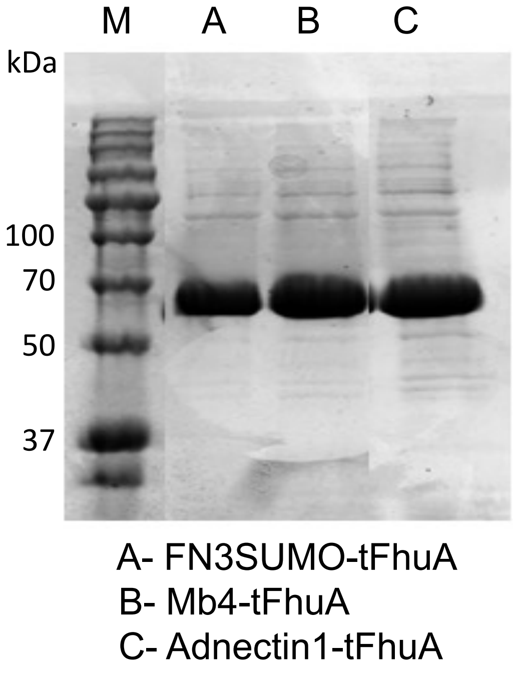


**<u>Supplementary Figure S1.</u> SDS-PAGE gel analysis of FN3SUMO-tFhuA, Mb4-tFhuA, and Adnectin1-tFhuA sensors.** The purity and size of all three proteins were checked by a 12% SDS-PAGE gel analysis. The expected molecular weights (MW) were observed on gel.

### *3. Three-dimensional structure predictions of monobody-containing nanopore sensors.*

**a**

VSDVPRDLEVVAATPTSLLISWDAPAVTVRYYRITYGETGGNSPVQEFTVPGSKSTATISGLKPGV
DYTITVYAVTGRGDSPASSKPISINYRTGGSGGSLKEVQFKAGTDSLFQTGFDFSDSLDDDGVYS
YRLTGLARSANAQQKGSEEQRYAIAPAFTWRPDDKTNFTFLSYFQNEPETGNSEGSTYSRNEK
MVGYSFDHEFNDTFTVRQNLRFAENKTSQNSVYGNSEGSRKYVVDDEKLQNFSVDTQLQSKF
ATGDIDHTLLTGVDFMRMRNDINAWFGYNSEGSSGPYRILNKQKQTGVYVQDQAQWDKVLV
TLGGRYDWADQESLNRVAGTTDKRDDKQFTWRGGVNYLFDNGVTPYFSYSESFEPSSQVGK
DGNIFAPSKGKQYEVGVKYVPEDRPIVVTGAVYNLTKTNNLMADPEGSFFSVEGGEIRARGVEI
EAKAALSASVNVVGSYTYTDAEYTTDTTYKGNTPAQVPKHMASLWADYTFFDGPLSGLTLGTG
GRYTNSEGSYTVVDALVRYDLARVGMAGSNVALHVNSEGSQVVATATFRF

**b**

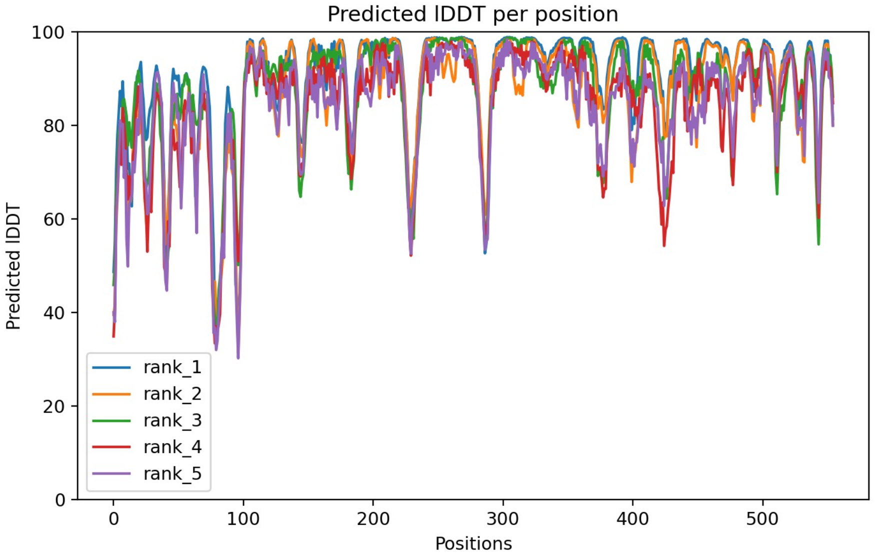


**c**

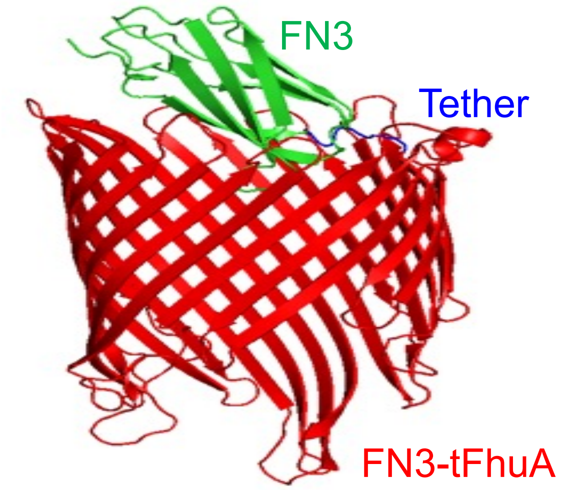


**Supplementary Figure S2. Structure prediction of FN3-tFhuA using AlphaFold2.** **(a)** Amino acid sequence of FN3-tFhuA. **(b)** The predicted Local Distance Difference Test (pLDDT) per residue was between 80 and 100 for most residues. pLDDT per residue was extracted using AlphaFold2.[8, 9] **(c)** The structure of FN3-tFhuA predicted by AlphaFold2, demonstrating the orientation of the parent FN3 domain (green) with respect to the central axis of tFhuA (red).

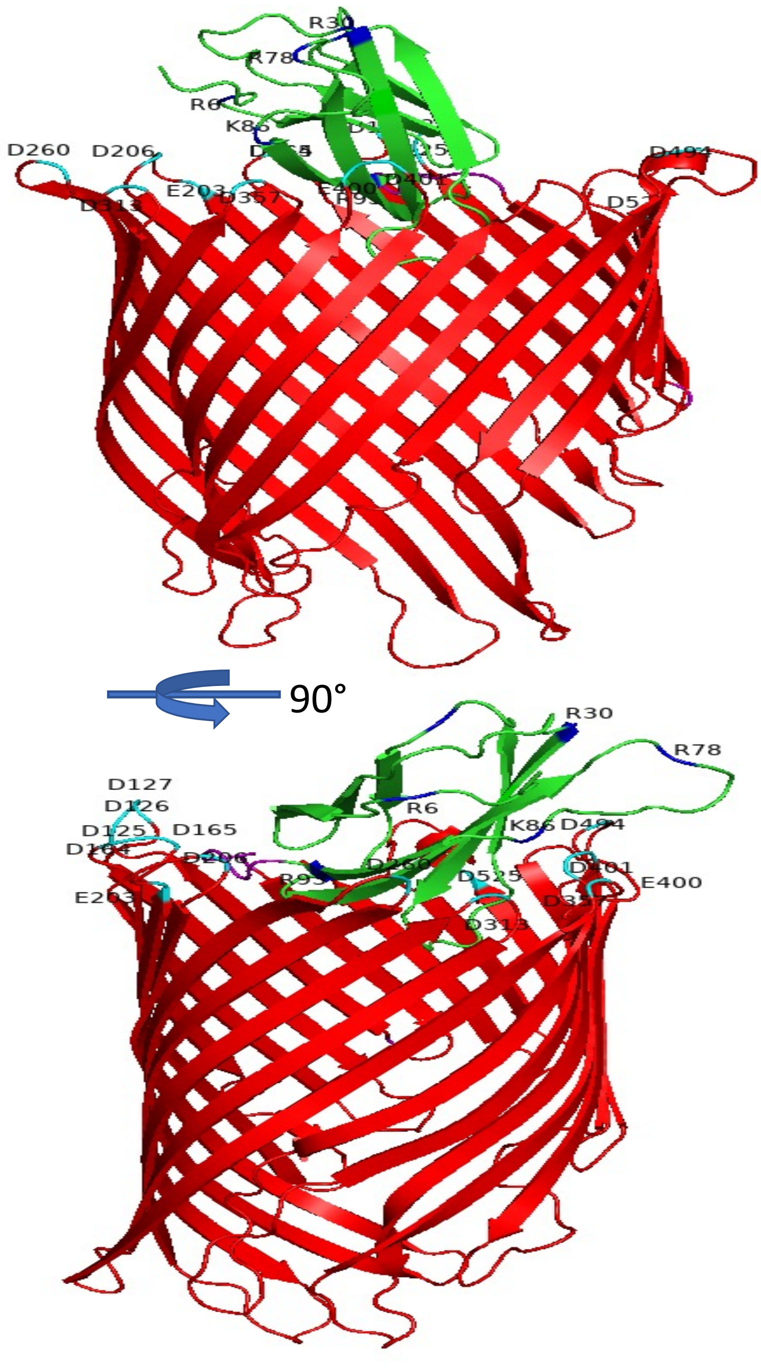


**<u>Supplementary Figure S3.</u> Negative charge distribution on the β turns of tFhuA and positive charge distribution on the loops of FN3.** FN3 domain was covalently attached to the tFhuA at the N terminus via a $(GGS)_2$ tether. The negatively and positively charged residues are shown in cyan and blue, respectively. The negatively charged residues on the β turns of tFhuA are exposed to most positively charged residues on the loops of the FN3 domain.

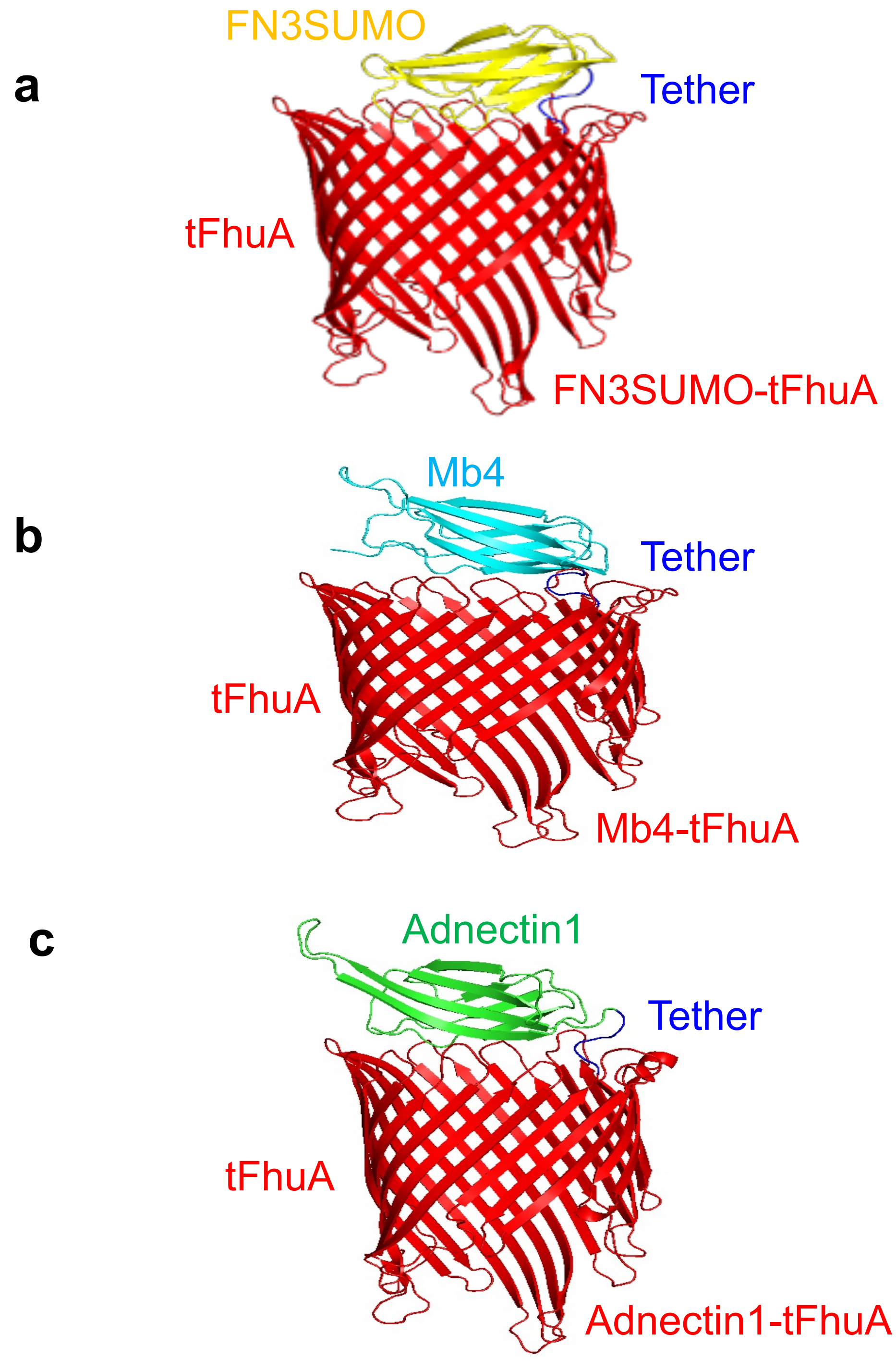


**<u>Supplementary Figure S4.</u> Structural predictions of FN3SUMO-tFhuA, Mb4-tFhuA, and Adnectin1-tFhuA using AlphaFold2.**[8, 9] **(a)** FN3SUMO-tFhuA. **(b)** Mb4-tFhuA. **(c)** Adnectin1-tFhuA. These structures were replicated in at least two independent runs using Alphafold2 on Google Colab server (https://colab.research.google.com/github/sokrypton/ColabFold/blob/main/AlphaFold2.ipynb). Identical 3D structures were observed in each run.

***4. Single-channel electrical signatures and unitary conductance values of the three monobody-containing sensors.***

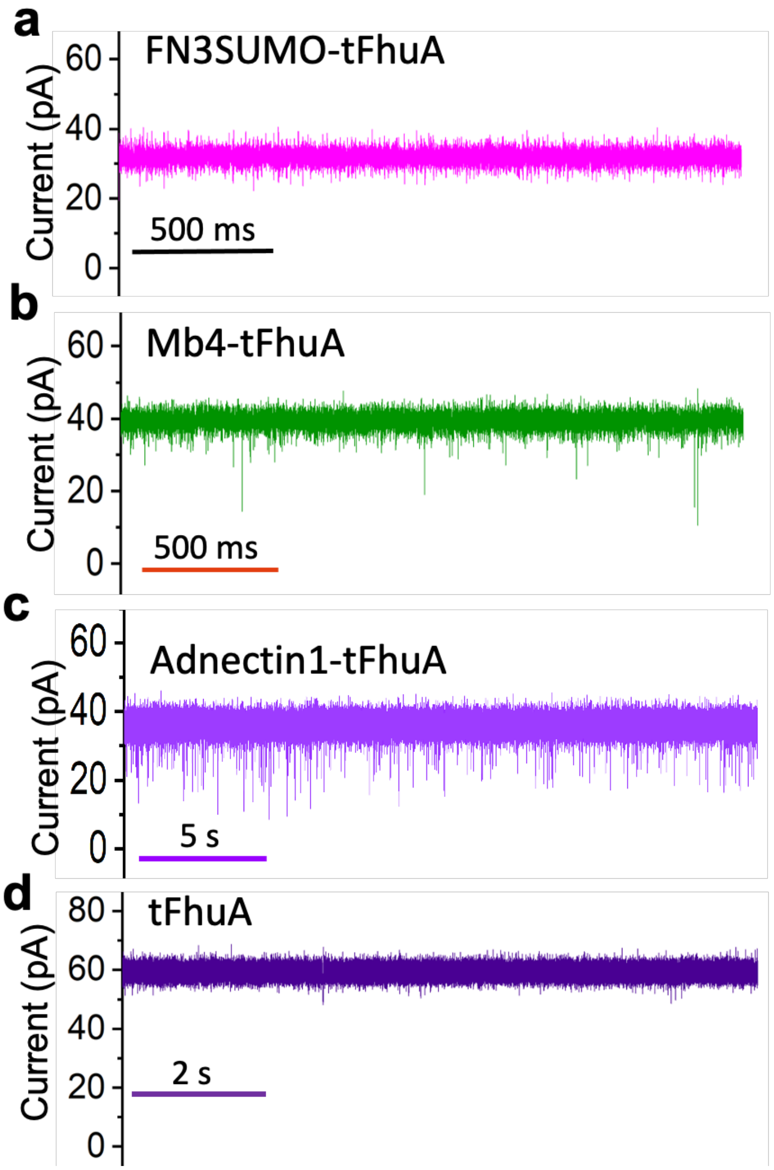


<u>Supplementary Figure S5.</u> **Single-channel electrical traces of modified and unmodified tFhuA. (a)** A representative single-channel electrical trace of FN3SUMO-tFhuA, indicating a unitary conductance of ~0.8 nS. This single-channel trace was low-pass Bessel filtered at a frequency of 3 kHz. **(b)** A representative single-channel electrical trace of Mb4-tFhuA, showing a unitary conductance of ~1 nS. This single-channel electrical trace was low-pass Bessel filtered at a frequency of 3 kHz. **(c)** A representative single-channel electrical trace of Adnectin1-tFhuA, indicating a unitary conductance of ~0.9 nS. This single-channel electrical trace was low-pass Bessel filtered at a frequency of 4 kHz. **(d)** A representative single-channel trace of the unmodified tFhuA, showing a unitary conductance of ~1.5 nS. This single-channel trace was low-pass Bessel filtered at a frequency of 3 kHz. All traces were recorded at a transmembrane potential of +40 mV. All recordings were performed in 300 mM KCl, 10 mM Tris-HCl, pH 8.0. These recordings were replicated in n = 3 independent experiments.

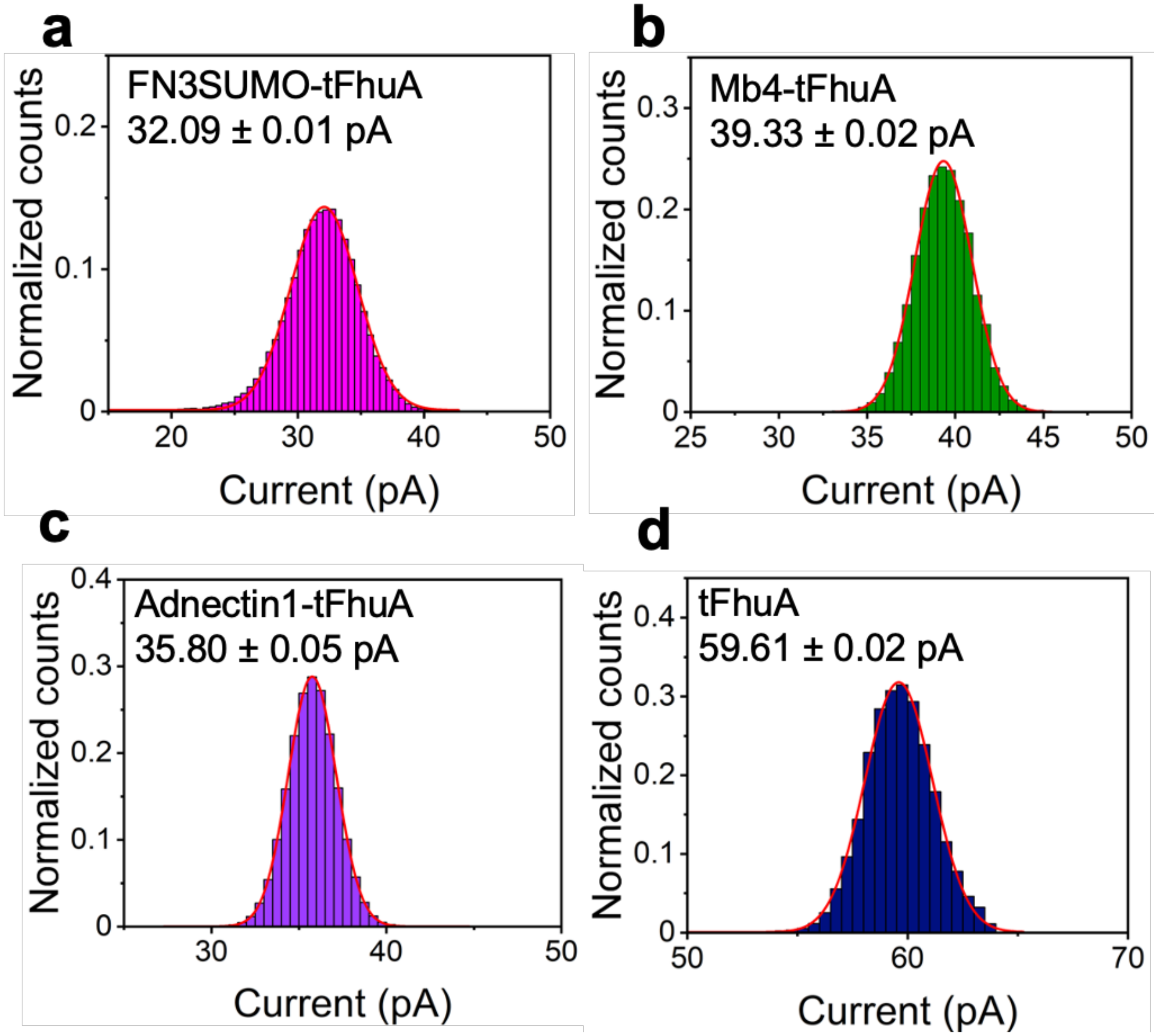


**<u>Supplementary Figure S6.</u> Normalized all-point histograms of the open-substate currents. (a)** FN3SUMO-tFhuA. **(b)** Mb4-tFhuA. **(c)** Adnectin1-tFhuA. **(d)** tFhuA. All recordings were performed at an applied potential of +40 mV and in 300 mM KCl, 10 mM Tris-HCl, pH 8.0. These recordings were replicated in n = 3 independent experiments.

**Supplementary Table S2. Unitary conductance values of the engineered monobody-based sensors, and their comparisons with the unmodified tFhuA nanopore.**

| Nanopore sensor | Applied potential (mV) | Conductance (nS) |
|---|---|---|
| **FN3SUMO-tFhuA** | +40 | 0.81 ± 0.03 |
| **Mb4-tFhuA** | +40 | 0.99 ± 0.04 |
| **Adnectin1-tFhuA** | +40 | 0.90 ± 0.02 |
| **tFhuA (control)** | +40 | 1.52 ± 0.10 |

Values represent mean ± s.d. from n = 3 distinct experiments. The experimental conditions were the same as those stated in **Methods**.

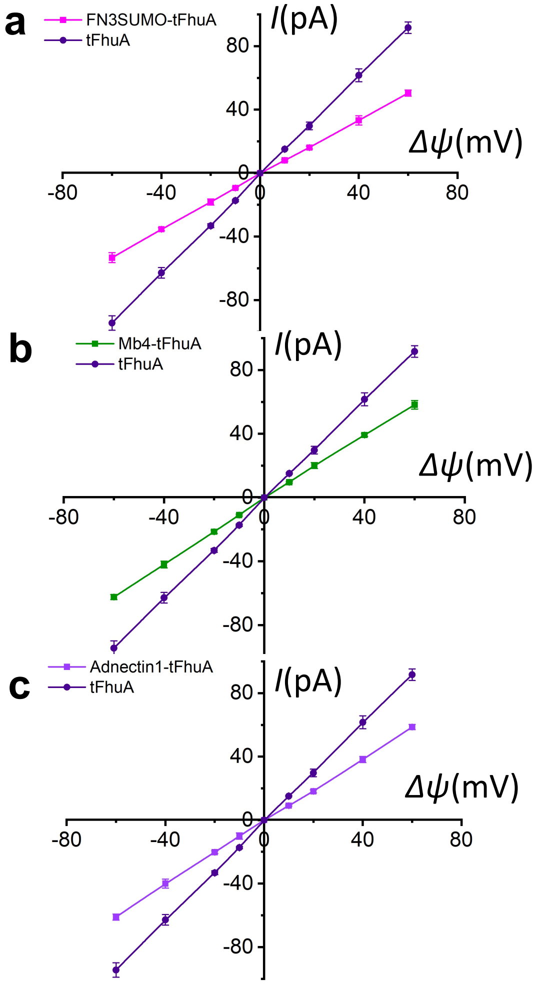


**Supplementary Figure S7. The current-voltage curves of the three sensors and their comparisons with the unmodified tFhuA nanopore. (a)** FN3SUMO-tFhuA. **(b)** Mb4-tFhuA. **(c)** Adnectin1-tFhuA. Here, $I$ and $\Delta\Psi$ indicate the unitary current and applied transmembrane potential, respectively. Data are mean ± s.d. from n = 3 independent experiments. The other experimental conditions were the same as those stated in **Methods**.

*5. hSUMO1-induced current blockades noted with the FN3SUMO-tFhuA sensor.*

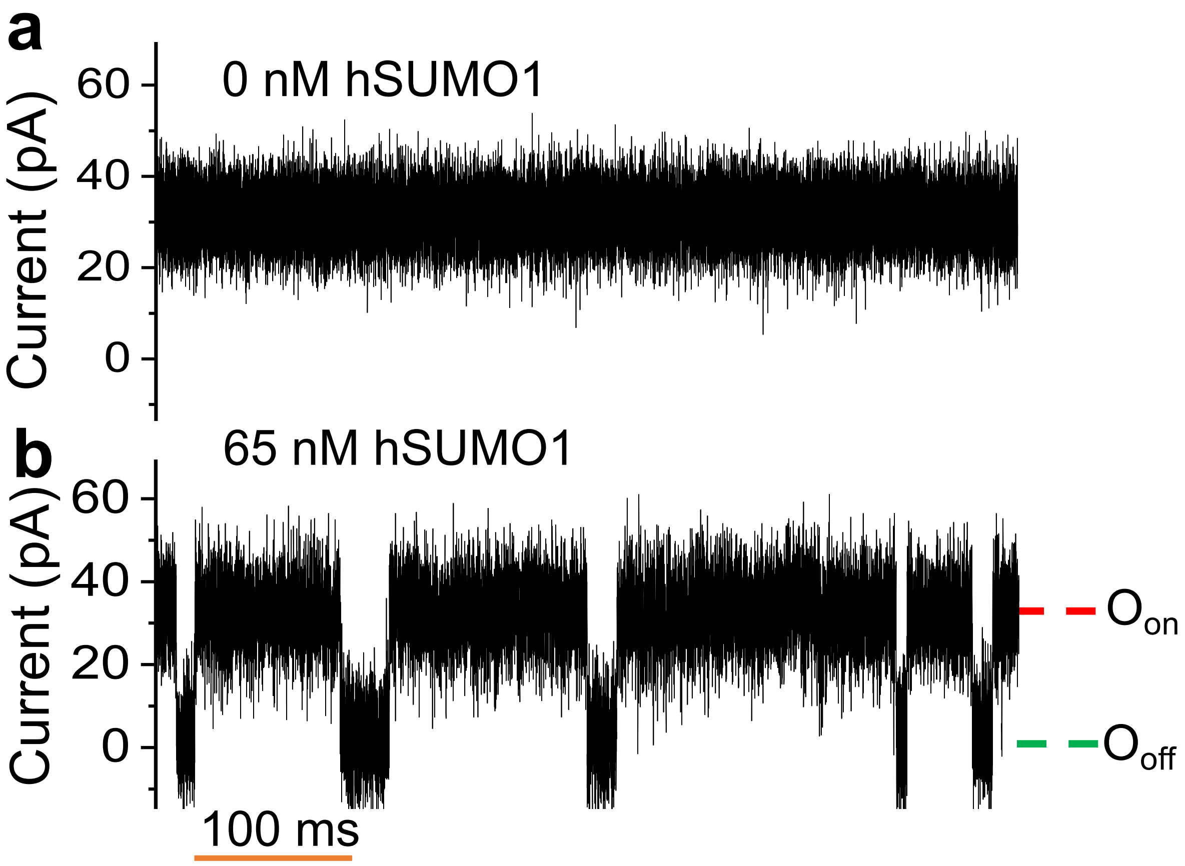


<u>Supplementary Figure S8.</u> **Effect of hSUMO1 on the raw signature of the FN3SUMO-tFhuA sensor. (a)** Single-channel electrical trace was recorded in the absence of hSUMO1. **(b)** A representative raw trace of FN3SUMO-tFhuA when 65 nM hSUMO1 was added to the *cis* side of the chamber. These traces were low-pass filtered using an 8-pole Bessel filter at a frequency of 7 kHz. The other experimental conditions are stated in **Methods**. This single-channel electrical signature was replicated in n = 3 independent experiments.

*6. Negative- and positive-control experiments with the FN3SUMO-tFhuA sensor.*

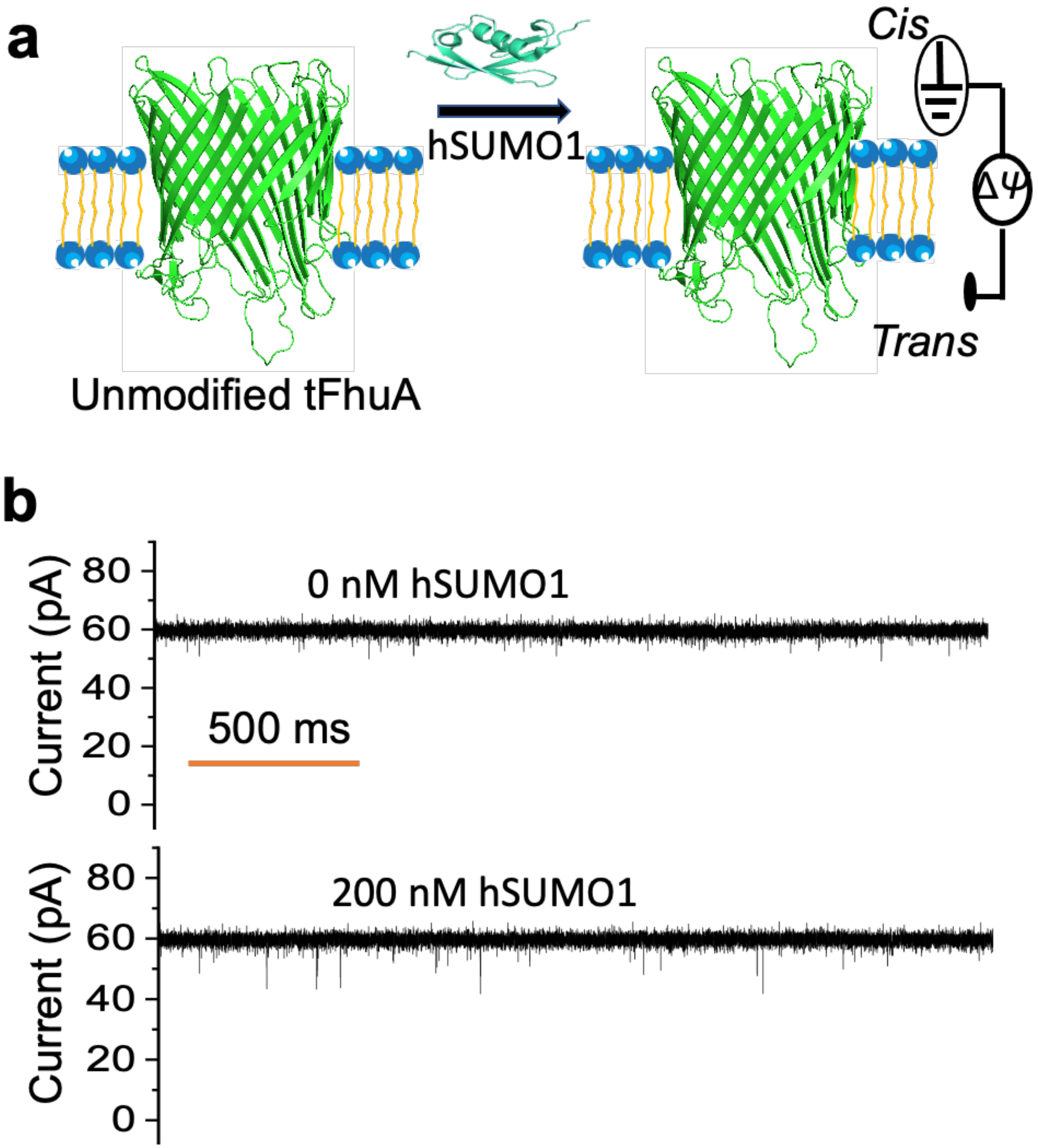


Supplementary Figure S9. **A negative-control experiment for testing nonspecific binding of hSUMO1 with the "*cis*" pore opening. (a)** The unmodified tFhuA was functionally reconstituted into a lipid bilayer. **(b)** Upper trace is showing the signature when no hSUMO1 was added. tFhuA shows a conductance of ~1.5 nS. In the lower trace, 200 nM hSUMO1 was added to the *cis* compartment. Very rare and brief current blockades were noted. The applied transmembrane potential was +40 mV. The buffer solution contained 300 mM KCl, 10 mM Tris-HCl, pH 8.0. Single-channel traces were low-pass filtered using an 8-pole Bessel at a frequency of 2 kHz. These recordings were replicated in n = 3 independent experiments.

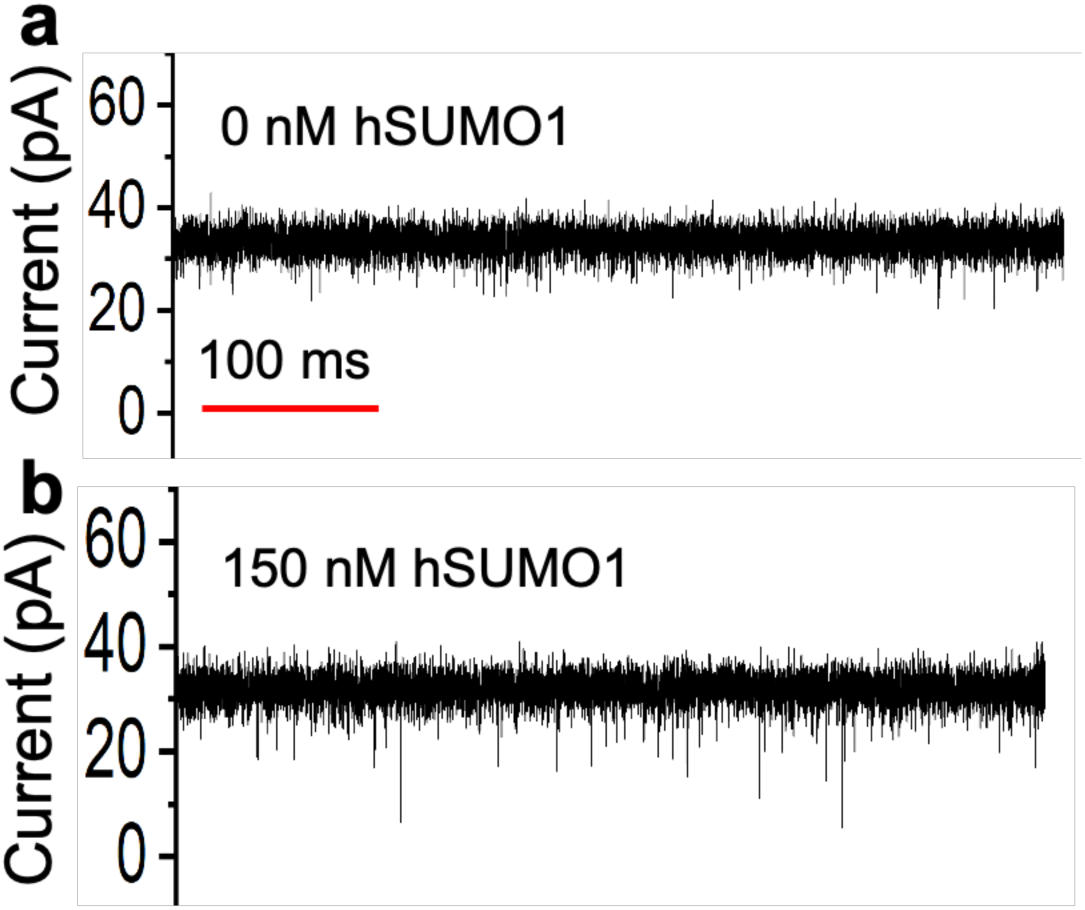


**Supplementary Figure S10. Examining the orientation of the FN3SUMO-tFhuA sensor.** **(a)** No hSUMO1 was added to the *trans* side. **(b)** The addition of 150 nM hSUMO1 to the *trans* side did produce rare and brief current transitions when an FN3SUMO-tFhuA sensor was functionally reconstituted into a lipid bilayer. This result confirmed the insertion of the FN3SUMO-tFhuA nanopore into a lipid bilayer with a preferred orientation.[10] Single-channel electrical traces were low-pass Bessel filtered at a frequency of 4 kHz. These single-channel electrical traces were acquired in two replicates. All recordings were performed under an applied potential of 40 mV and in 300 mM KCl, 0.5 mM TCEP, 10 mM Tris-HCl, pH 8.0.

*7. Time, rate, and equilibrium constants of* **hSUMO1–FN3SUMO interactions.**

**Supplementary Table S3. Mean values of the release ($\tau_{on}$) and capture durations ($\tau_{off}$) of hSUMO1-FN3SUMO interactions using an FN3SUMO-tFhuA sensor.**

| [hSUMO1] (nM) | $\tau_{on}$ (ms) | $\tau_{off}$ (ms) |
|---|---|---|
| 65 | 116 ± 8 | 15 ± 2 |
| 130 | 60 ± 9 | 13 ± 3 |
| 260 | 32 ± 3 | 12 ± 1 |
| 520 | 18 ± 4 | 14 ± 3 |
| 1040 | 10 ± 1 | 13 ± 2 |

Values are mean ± s.d. from $n$ = 3 independent experiments. The applied transmembrane potential was +40 mV. The buffer solution contained 300 mM KCl, 10 mM Tris-HCl, 0.5 mM TCEP, pH 8.0.

**Supplementary Table S4. Mean values of the association ($k_{on}$) and dissociation ($k_{off}$) rate constants of hSUMO1-FN3SUMO interactions.** Here, $k_{on} = 1/([\text{hSUMO1}]\tau_{on})$ and $k_{off} = 1/\tau_{off}$.

| [hSUMO1] (nM) | $k_{on}$ ($M^{-1}s^{-1}$) × $10^{-8}$ | $k_{off}$ ($s^{-1}$) |
|---|---|---|
| 65 | 1.3 ± 0.1 | 67 ± 9 |
| 130 | 1.3 ± 0.2 | 74 ± 12 |
| 260 | 1.2 ± 0.1 | 81 ± 5 |
| 520 | 1.1 ± 0.2 | 72 ± 7 |
| 1040 | 1.0 ± 0.1 | 78 ± 12 |

Values are mean ± s.d. from $n$ = 3 independent experiments. The applied transmembrane potential was +40 mV. The buffer solution contained 300 mM KCl, 10 mM Tris-HCl, 0.5 mM TCEP, pH 8.0.

**Supplementary Table S5. The association ($k_{on}$) and dissociation ($k_{off}$) rate constants and equilibrium dissociation constant ($K_D$) of hSUMO1-FN3SUMO interactions.** $k_{on}$ value is the slope of the linear fit in **Fig. 2f.** $k_{off}$ value is the axis intercept of the horizontal line fit in **Fig. 2g.** Here, $K_D$ was calculated using the equation $K_D = k_{off}/k_{on}$.

| $k_{on}$ ($M^{-1}s^{-1}$) × $10^{-8}$ | $k_{off}$ ($s^{-1}$) | $K_D$ (nM) |
|---|---|---|
| 1.12 ± 0.02 | 74.5 ± 2.4 | 665 ± 24 |

Values are provided as mean ± s.e.m. The other experimental conditions were the same as those stated in **Methods**.

***8. WDR5-induced current blockades noted with the Mb4-tFhuA sensor.***

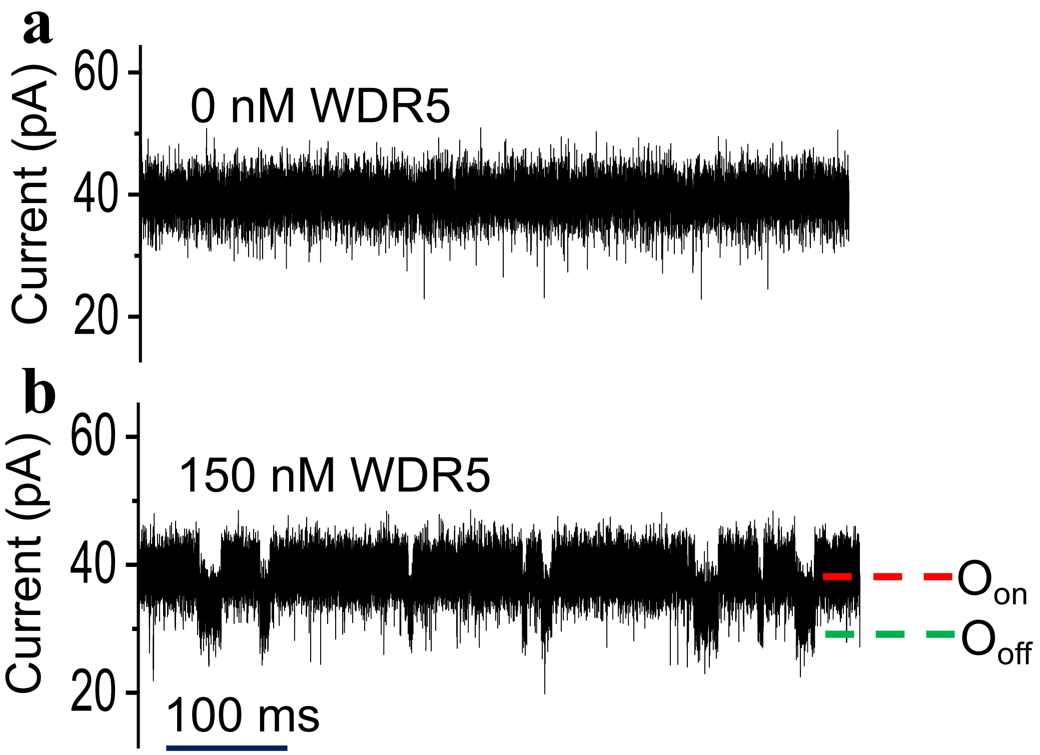


<u>**Supplementary Figure S11.**</u> **A representative raw trace of the Mb4-tFhuA sensor when 150 nM WDR5 was added to the *cis* side of the chamber. (a)** The single-channel trace was recorded in the absence of WDR5. **(b)** The single-channel trace was recorded in the presence of 150 nM WDR5 added to the *cis* side of the chamber. These traces were low-pass filtered using an 8-pole Bessel filter at a frequency of 7 kHz. The other experimental conditions are stated in **Methods**. This single-channel electrical signature was replicated in n = 3 independent experiments.

***9. Negative- and positive-control experiments with the Mb4-tFhuA sensor.***

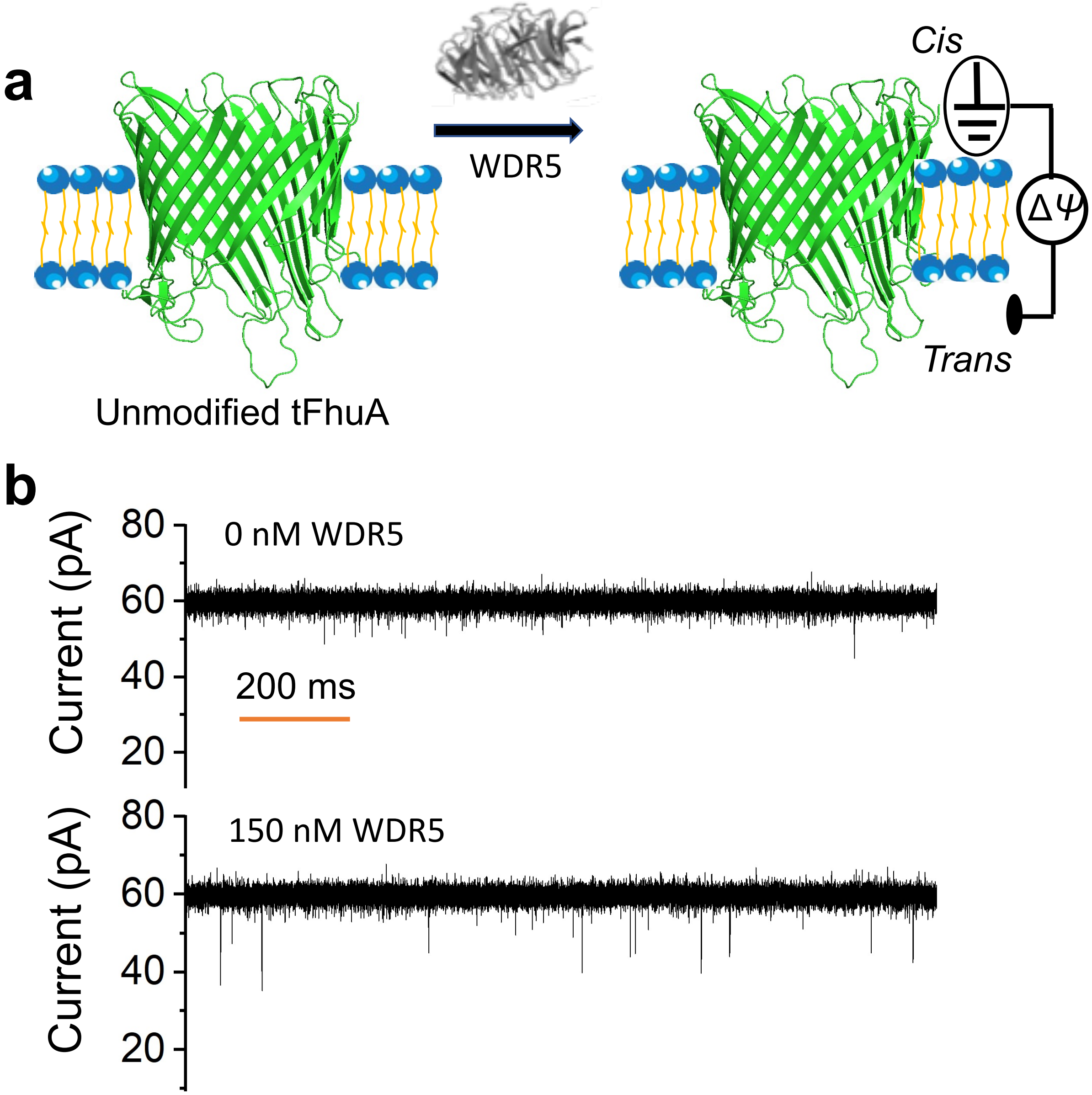


__Supplementary Figure S12.__ **Testing nonspecific bindings of WDR5 to unmodified tFhuA (the negative-control experiment). (a)** The unmodified tFhuA was reconstituted into lipid bilayer. **(b)** Fully open tFhuA shows a higher unitary conductance (~1.5 nS). The upper trace shows the signature when no WDR5 was added to the chamber. In the lower panel, WDR5 was added to the *cis* compartment. Some nonspecific binding events of WDR5 with tFhuA were noted in the form of low-amplitude and brief current spikes, likely due to collisions of WDR5 with the opening of tFhuA. The applied transmembrane potential was +40 mV. The buffer solution contained 300 mM KCl, 10 mM Tris-HCl, pH 8.0. These traces were replicated in n = 3 independent experiments. Single-channel electrical traces were low-pass filtered using an 8-pole Bessel filter at a frequency of 2 kHz.

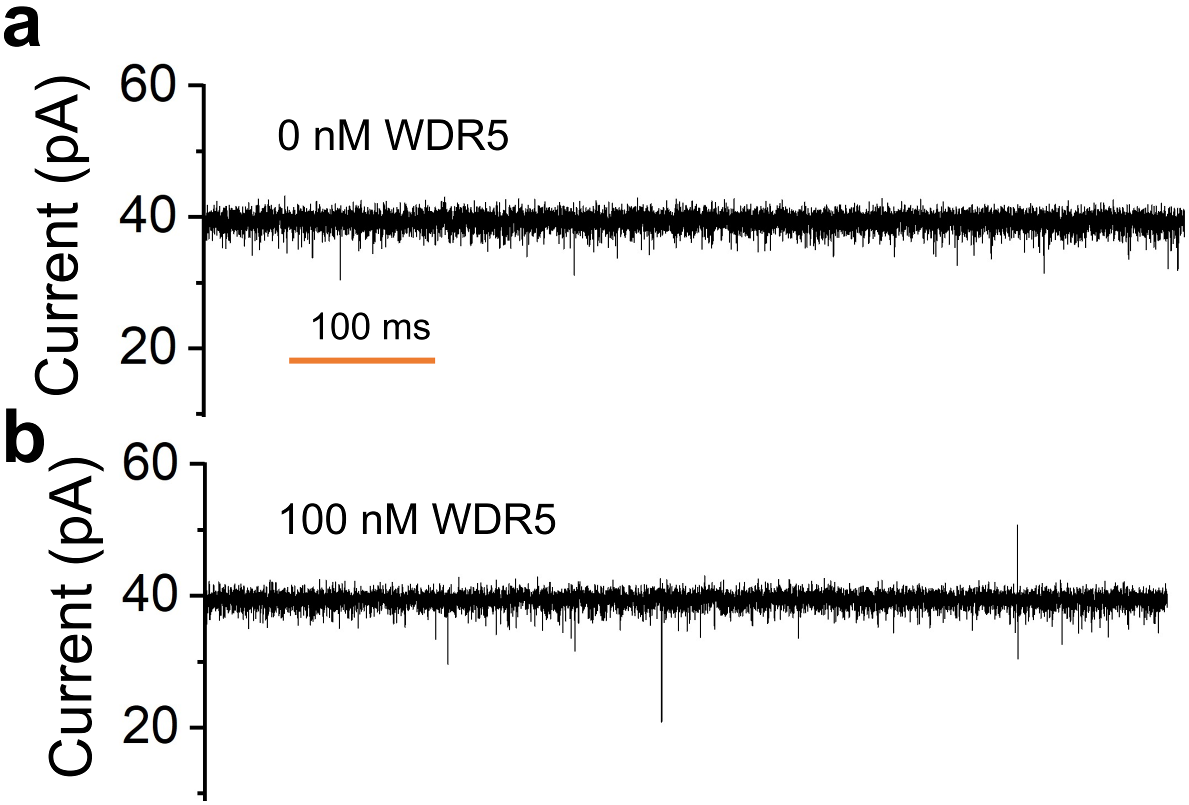


<u>Supplementary Figure S13.</u> **The addition of WDR5 to the *trans* side of a Mb4-tFhuA-containing lipid bilayer did not produce long-lived current transitions (the positive-control experiment). (a)** No WDR5 was added to the *trans* side. (**b**) 100 nM WDR5 was added to the *trans* side. This single-channel electrical signature was replicated in n = 3 distinct experiments. This finding validated the successful insertion of the Mb4-tFhuA nanopore into the lipid bilayer in a favored orientation, as predicted by a previous study.[10] Single-channel electrical traces were low-pass filtered using an 8-pole Bessel filter at a frequency of 2 kHz. All recordings were performed under an applied potential of +40 mV, and in 300 mM KCl, 1 mM TCEP, 10 mM Tris-HCl, pH 8.0.

***10. Time, rate, and equilibrium constants of WDR5–Mb4 interactions.***

**Supplementary Table S6. Mean values of the release ($\tau_{on}$) and capture durations ($\tau_{off}$) of WDR5-Mb4 interactions using an Mb4-tFhuA sensor.**

| [WDR5] (nM) | $\tau_{on}$ (ms) | $\tau_{off}$ (ms) |
|---|---|---|
| 50 | 178 ± 11 | 12 ± 3 |
| 150 | 65 ± 12 | 12 ± 5 |
| 300 | 34 ± 6 | 14 ± 3 |
| 600 | 20 ± 5 | 17 ± 4 |
| 1200 | 10 ± 3 | 13 ± 2 |
| 2400 | 5 ± 1 | 15 ± 1 |

Values are mean ± s.d. from n = 3 independent experiments. The applied transmembrane potential was +40 mV. The buffer solution contained 300 mM KCl, 10 mM Tris-HCl, 1 mM TCEP, pH 8.0.

**Supplementary Table S7. Mean values of the association ($k_{on}$) and dissociation ($k_{off}$) rate constants of WDR5-Mb4 interactions using an Mb4-tFhuA sensor.**
Here, $k_{on} = 1/([\text{WDR5}]\,\tau_{on})$ and $k_{off} = 1/\tau_{off}$.

| [WDR5] (nM) | $k_{on}$ ($M^{-1}s^{-1}$) × $10^{-8}$ | $k_{off}$ ($s^{-1}$) |
|---|---|---|
| 50 | 1.1 ± 0.1 | 87 ± 22 |
| 150 | 1.0 ± 0.2 | 88 ± 29 |
| 300 | 1.0 ± 0.2 | 73 ± 18 |
| 600 | 0.9 ± 0.2 | 62 ± 19 |
| 1200 | 0.9 ± 0.3 | 76 ± 8 |
| 2400 | 0.8 ± 0.1 | 67 ± 5 |

Values are mean ± s.d. from n =3 independent experiments. The applied transmembrane potential was +40 mV. The buffer solution contained 300 mM KCl, 10 mM Tris-HCl, 1 mM TCEP, pH 8.0.

**Supplementary Table S8. The association ($k_{on}$) and dissociation ($k_{off}$) rate constants and equilibrium dissociation constant ($K_D$) of WDR5-Mb4 interactions.** $k_{on}$ value is the slope of the linear fit in **Fig. 3f.** $k_{off}$ value is the axis intercept of the horizontal line fit in **Fig. 3g.** Here, $K_D$ was calculated using the equation $K_D = k_{off}/k_{on}$.

| $k_{on}$ ($M^{-1}s^{-1}$) × $10^{-8}$ | $k_{off}$ ($s^{-1}$) | $K_D$ (nM) |
|---|---|---|
| 0.83 ± 0.01 | 72.4 ± 3.7 | 872 ± 45 |

Values are mean ± s.e.m. The other experimental conditions were the same as those stated in **Methods**.

*11. Biolayer interferometry (BLI) measurements of WDR5–Mb4-tFhuA interactions.*

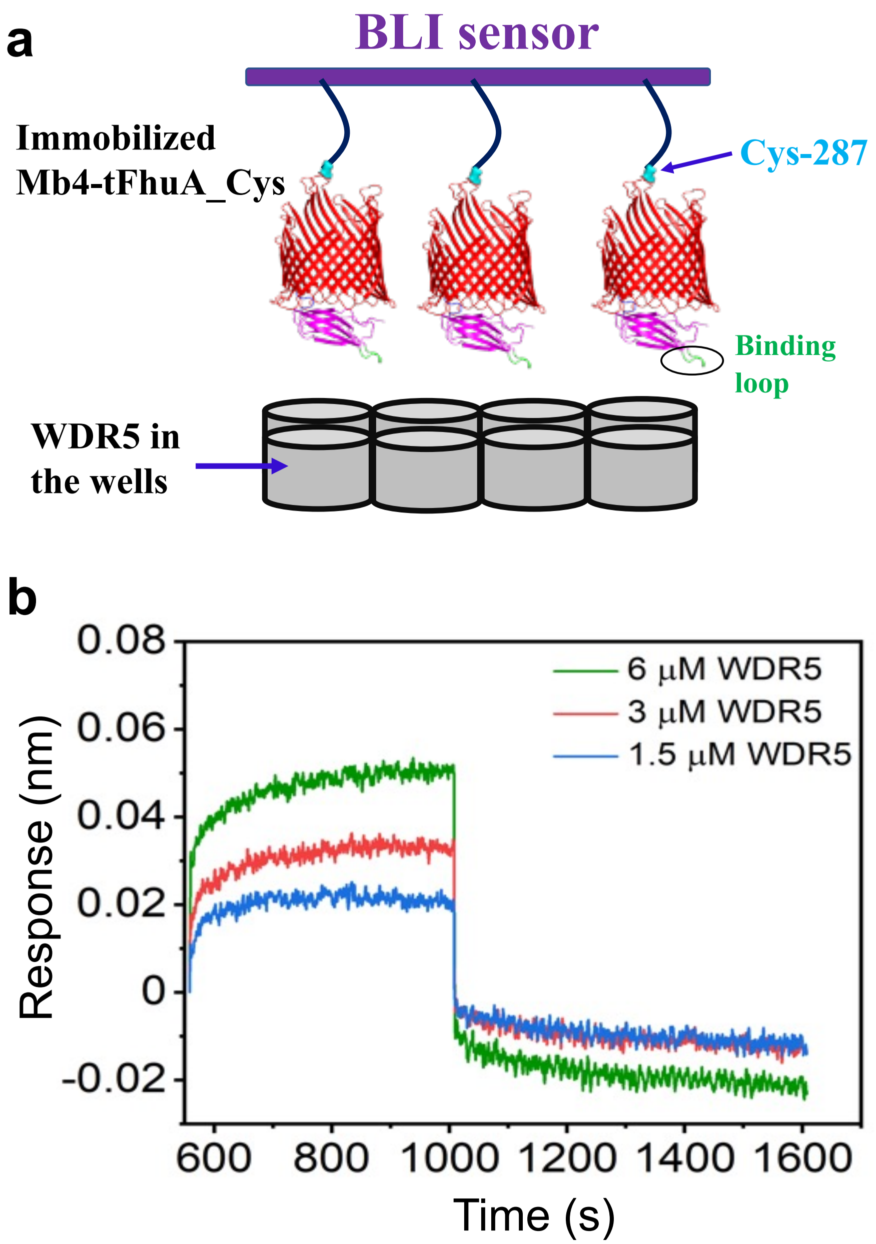


**<u>Supplementary Figure S14.</u> Biolayer interferometry (BLI) measurements of WDR5–Mb4-tFhuA interactions. (a)** In this case, Mb4-tFhuA_Cys287 was biotinylated at -$SH_2$ group of an engineered cysteine (C287) and immobilized onto streptavidin (SA) sensors' surface. WDR5 was added to the wells. **(b)** Real-time optical measurements of WDR5-Mb4 interactions. 50 nM biotin-tagged Mb4-tFhuA_Cys287 was loaded onto the SA sensor surface. Titration series of WDR5 (1.5, 3.0 and 6.0 μM) were employed as protein analytes. Corresponding association and dissociation curves are shown for these WDR5 concentrations. These BLI sensorgrams were acquired in n = 6 replicates.

*12. EGFR-induced current blockades noted with the Adnectin1-tFhuA sensor.*

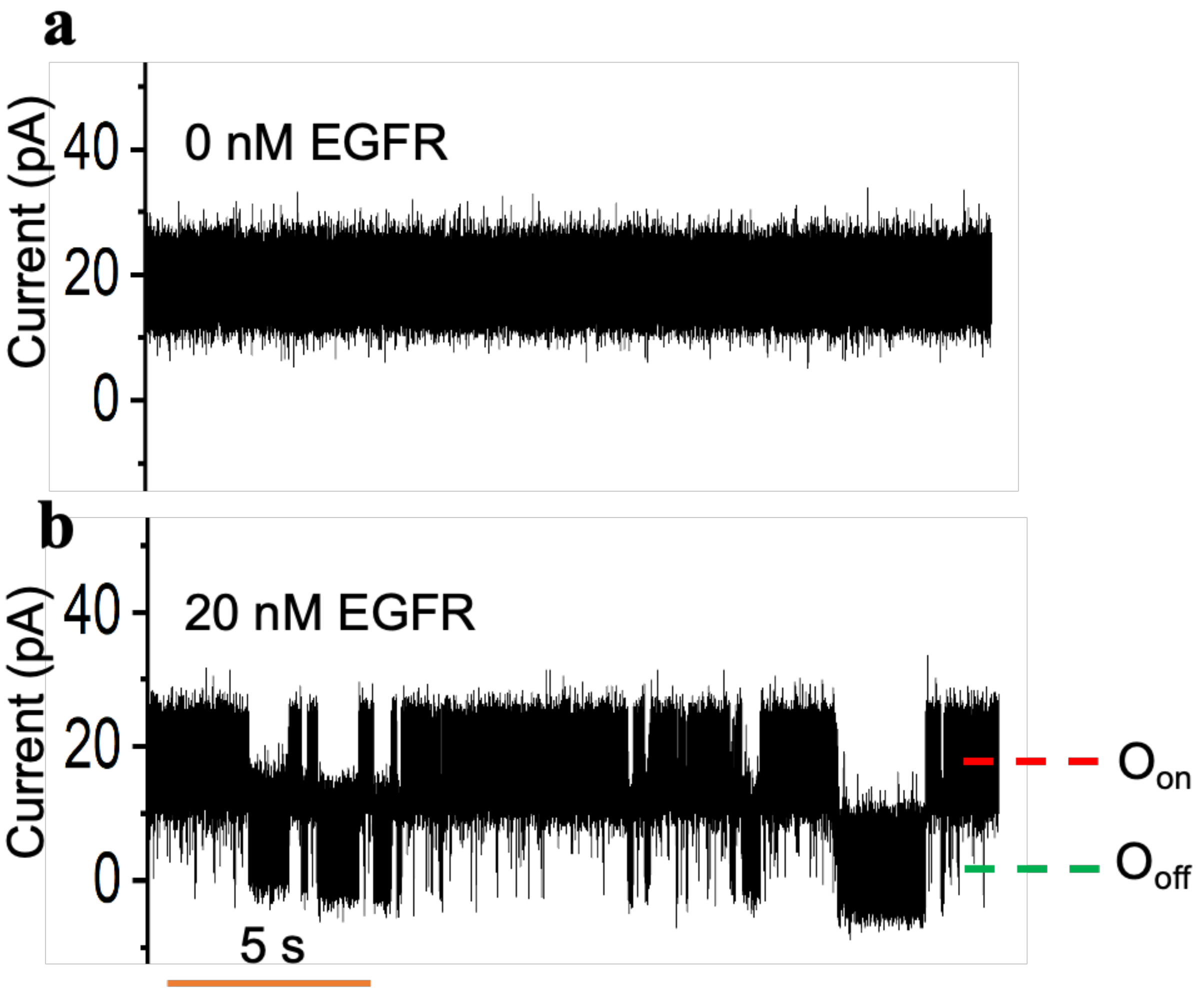


__Supplementary Figure S15.__ **A representative raw trace of the Adnectin1-tFhuA sensor when 20 nM EGFR was added to the *cis* side of the chamber. (a)** The single-channel trace was recorded in the absence of EGFR. **(b)** The single-channel trace was recorded in the presence of 20 nM EGFR added to the *cis* side of the chamber. These traces were low-pass filtered using an 8-pole Bessel filter at a frequency of 7 kHz. The other experimental conditions are stated in **Methods**. This single-channel electrical signature was replicated in n = 3 independent experiments.

*13. Negative-control experiments with the Adnectin1-tFhuA sensor.*

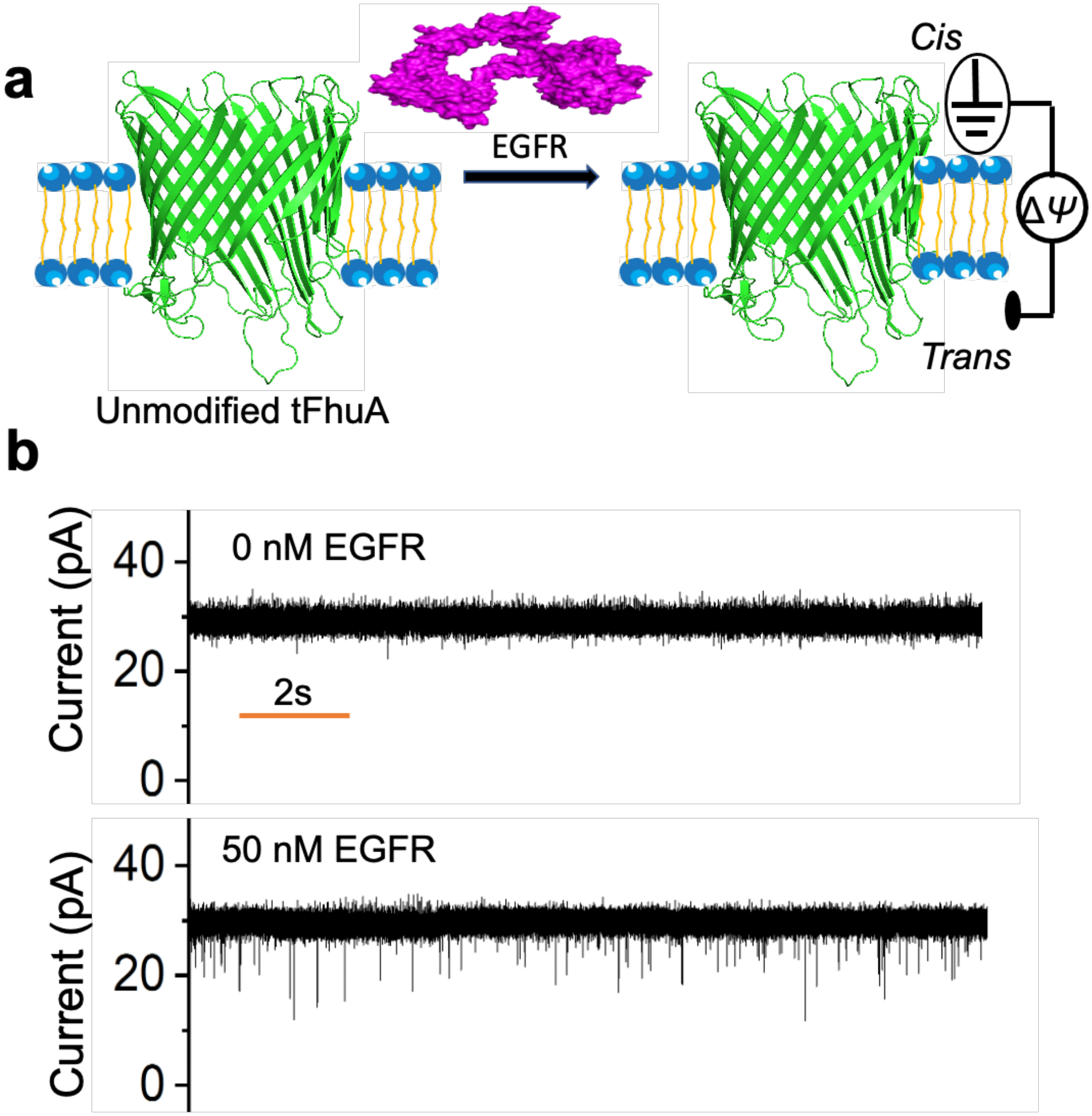


**<u>Supplementary Figure S16.</u> An experiment for testing nonspecific bindings of EGFR with the pore opening of tFhuA (the negative-control experiment). (a)** The unmodified tFhuA was functionally reconstituted into a lipid bilayer. EGFR (magenta) and tFhuA (green) are not represented on the same scale. **(b)** tFhuA shows a unitary conductance of ~1.5 nS. In the upper trace, no EGFR was added. In the lower trace, 50 nM EGFR was added to the *cis* compartment. Some low-amplitude brief current spikes were noted, presumably due to nonspecific binding events of EGFR to tFhuA. These events are perhaps due to high negative charge of EGFR at pH 8.0 (**Supplementary Table S1**). The applied transmembrane potential was +20 mV. The buffer solution contained 300 mM KCl, 10 mM Tris-HCl, pH 8.0. These traces were replicated in n = 3 independent experiments. Single-channel electrical traces were low-pass filtered using an 8-pole Bessel filter at a frequency of 2 kHz

*14. Normalized current amplitudes of EGFR-produced blockades.*

**Supplementary Table S9. Values of the mean normalized amplitude of EGFR-produced current blockades, $A/I_0$, for the two binding events at various [EGFR] values.** Here, $I_0$ and $A$ denote the single-channel current of the EGFR-released substate and the current amplitude of EGFR-produced current blockades, respectively. $A/I_0$ were then converted into percentages. $P_1$ and $P_2$ represent the probabilities of two distinct current blockades. A representative event histogram of normalized current amplitudes is shown in **Fig. 4b**.

| EGFR [nM] | $P_1$ | $A_1/I_0$ (%) | $P_2$ | $A_2/I_0$ (%) |
|---|---|---|---|---|
| 10 | 0.72 ± 0.01 | 63.8 ± 1.5 | 0.27 ± 0.01 | 85.1 ± 2.3 |
| 20 | 0.73 ± 0.02 | 64.9 ± 1.1 | 0.27 ± 0.02 | 84.4 ± 3.1 |
| 40 | 0.72 ± 0.02 | 65.0 ± 2.1 | 0.28 ± 0.02 | 86.1 ± 1.6 |
| 80 | 0.75 ± 0.03 | 67.3 ± 6.9 | 0.25 ± 0.03 | 85.2 ± 1.5 |
| 160 | 0.74 ± 0.04 | 66.6 ± 5.2 | 0.26 ± 0.04 | 86.8 ± 4.0 |
| 320 | 0.76 ± 0.04 | 66.0 ± 3.6 | 0.24 ± 0.04 | 87.3 ± 5.8 |

Values of the normalized current blockades are mean ± s.d. from n = 3 independent experiments. The other experimental conditions are indicated in **Methods**.

*15. Time, rate, and equilibrium constants of EGFR–Adnectin1 interactions.*

**Supplementary Table S10. The probability distribution of the two binding events of EGFR-Adnectin1 interactions.** These events were differentiated by the EGFR-captured duration. Individual experimental values were derived using event histograms in ClampFit (Axon) and fits in a semilogarithmic representation. The maximum likelihood method[11, 12] and logarithm likelihood ratio (LLR) tests[13-15] were used for all fits to determine the best multi-exponential distribution model (**Methods**).

| EGFR [nM] | $P_1$ | $P_2$ |
|---|---|---|
| 10 | 0.65 ± 0.05 | 0.34 ± 0.05 |
| 20 | 0.65 ± 0.02 | 0.34 ± 0.02 |
| 40 | 0.64 ± 0.05 | 0.35 ± 0.05 |
| 80 | 0.69 ± 0.03 | 0.30 ± 0.03 |
| 160 | 0.68 ± 0.02 | 0.31 ± 0.02 |
| 320 | 0.69 ± 0.05 | 0.31 ± 0.05 |

Values are mean ± s.d. from n = 3 independent experiments. $P_1$ and $P_2$ are the probabilities of the short- and long-lived EGFR-captured events, respectively. The applied transmembrane potential was +20 mV. The buffer solution contained 300 mM KCl, 10 mM Tris-HCl, pH 8.0.

**Supplementary Table S11. Mean values of durations of the short-lived ($\tau_{off\text{-}1}$) and long-lived ($\tau_{off\text{-}2}$) EGFR-captured events.** All histogram fittings were conducted using a semilogarithmic representation. The maximum likelihood method[11, 12] and logarithm likelihood ratio (LLR) tests[13-15] were used for all fits to determine the best multi-exponential distribution model (**Methods**).

| EGFR [nM] | $\tau_{off\text{-}1}$ (ms) | $\tau_{off\text{-}2}$ (ms) |
|---|---|---|
| 10 | 88 ± 19 | 1016 ± 135 |
| 20 | 83 ± 12 | 950 ± 150 |
| 40 | 76 ± 10 | 1000 ± 199 |
| 80 | 81 ± 23 | 1060 ± 140 |
| 160 | 91 ± 11 | 1013 ± 111 |
| 320 | 92 ± 13 | 1005 ± 109 |

Values are mean ± s.d. from n = 3 independent experiments. The applied transmembrane potential was +20 mV. The buffer solution contained 300 mM KCl, 10 mM Tris-HCl, pH 8.0.

**Supplementary Table S12. Mean values of the durations $\tau_{on}$, $\tau_{on\text{-}1}$ and $\tau_{on\text{-}2}$ of EGFR-released events.** Subscripts "1" and "2" stand for the short- and long-lived EGFR-captured events, respectively. $\tau_{on}$ are mean values of the single-exponential distributions of EGFR-released duration histograms. $\tau_{on\text{-}1} = \tau_{on}/P_1$, where $P_1$ is the event probability of the short-lived events for each experiment. $\tau_{on\text{-}2} = \tau_{on}/P_2$, where $P_2$ is the event probability of the long-lived events for each experiment. The mean values of those probabilities are listed in **Supplementary Table S10**. All histogram fits were conducted using a semilogarithmic representation. The maximum likelihood method[11, 12] and logarithm likelihood ratio (LLR)[14, 16] tests were used for all fits to determine the best multi-exponential distribution model (**Methods**).

| EGFR [nM] | $\tau_{on}$ (S) | $\tau_{on\text{-}1}$ (S) | $\tau_{on\text{-}2}$ (S) |
|---|---|---|---|
| 10 | 0.79 ± 0.07 | 1.2 ± 0.1 | 2.5 ± 0.2 |
| 20 | 0.41 ± 0.06 | 0.62 ± 0.07 | 1.2 ± 0.2 |
| 40 | 0.24 ± 0.04 | 0.37 ± 0.05 | 0.70 ± 0.18 |
| 80 | 0.12 ± 0.02 | 0.18 ± 0.04 | 0.41 ± 0.05 |
| 160 | 0.07 ± 0.03 | 0.10 ± 0.04 | 0.21 ± 0.08 |
| 320 | 0.035 ± 0.005 | 0.05 ± 0.01 | 0.11 ± 0.01 |

Values are mean ± s.d. from n = 3 independent experiments. The applied transmembrane potential was +20 mV. The buffer solution contained 300 mM KCl, 10 mM Tris-HCl, pH 8.0.

**<u>Supplementary Table S13.</u> Mean values of the association rate constants corresponding to the short-lived ($k_{on-1}$) and long-lived ($k_{on-2}$) current blockades produced by EGFR-Adnectin1 interactions.** Here, $k_{on-i} = 1/([EGFR]\tau_{on-i})$ (i = 1, 2).

| EGFR [nM] | $k_{on-1}$ ($M^{-1}s^{-1}$) × $10^{-7}$ | $k_{on-2}$ ($M^{-1}s^{-1}$) × $10^{-7}$ |
|---|---|---|
| 10 | 8.8 ± 1.0 | 4.1 ± 0.3 |
| 20 | 8.1 ± 1.1 | 4.4 ± 0.7 |
| 40 | 6.8 ± 0.9 | 3.8 ± 1.2 |
| 80 | 7.1 ± 1.7 | 3.1 ± 0.4 |
| 160 | 7.2 ± 3.5 | 3.3 ± 1.6 |
| 320 | 6.2 ± 1.4 | 2.8 ± 0.3 |

Values are mean ± s.d. from n = 3 independent experiments. The applied transmembrane potential was +20 mV. The buffer solution contained 300 mM KCl, 10 mM Tris-HCl, pH 8.0.

**<u>Supplementary Table S14.</u> Mean values of the dissociation rate constants corresponding to the short-lived ($k_{off-1}$) and long-lived ($k_{off-2}$) EGFR-captured events of EGFR-Adnectin1 interactions.** Here, $k_{off-i} = 1/\tau_{off-i}$ (i = 1, 2).

| EGFR [nM] | $k_{off-1}$ ($s^{-1}$) | $k_{off-2}$ ($s^{-1}$) |
|---|---|---|
| 10 | 12 ± 2 | 1.0 ± 0.1 |
| 20 | 12 ± 2 | 1.1 ± 0.2 |
| 40 | 13 ± 2 | 1.0 ± 0.2 |
| 80 | 13 ± 4 | 1.0 ± 0.1 |
| 160 | 11 ± 1 | 1.0 ± 0.1 |
| 320 | 11 ± 2 | 1.0 ± 0.1 |

Values are mean ± s.d. from n = 3 independent experiments. The applied transmembrane potential was +20 mV. The buffer solution contained 300 mM KCl, 10 mM Tris-HCl, pH 8.0.

**<u>Supplementary Table S15.</u> The association and dissociation rate constants and the equilibrium dissociation constant, $K_D$, of EGFR-Adnectn1 interactions.** $k_{on\text{-}1}$ and $k_{on\text{-}2}$ values are the slopes of the linear fits in **Fig. 4e.** $k_{off\text{-}1}$ and $k_{off\text{-}2}$ values are the axis intercepts of the horizontal line fits in **Fig. 4f.** $K_{D\text{-}1}$ and $K_{D\text{-}2}$ are the equilibrium dissociation constants of the short-lived and long-lived current blockades. Here, $K_D$ was calculated using the equation $K_D = k_{off}/k_{on}$.

| $k_{on\text{-}1}$ ($M^{-1}s^{-1}$) × $10^{-7}$ | $k_{on\text{-}2}$ ($M^{-1}s^{-1}$) × $10^{-7}$ | $k_{off\text{-}1}$ ($s^{-1}$) | $k_{off\text{-}2}$ ($s^{-1}$) | $K_{D\text{-}1}$(nM) | $K_{D\text{-}2}$ (nM) |
|---|---|---|---|---|---|
| 6.62 ± 0.21 | 2.89 ± 0.10 | 12.0 ± 0.4 | 1.01 ± 0.01 | 181 ± 8 | 34 ± 2 |

Values are mean ± s.e.m. The other experimental conditions were the same as those stated in **Methods**.

*16. Structures of Adnectin1 and EGF in complex with EGFR.*

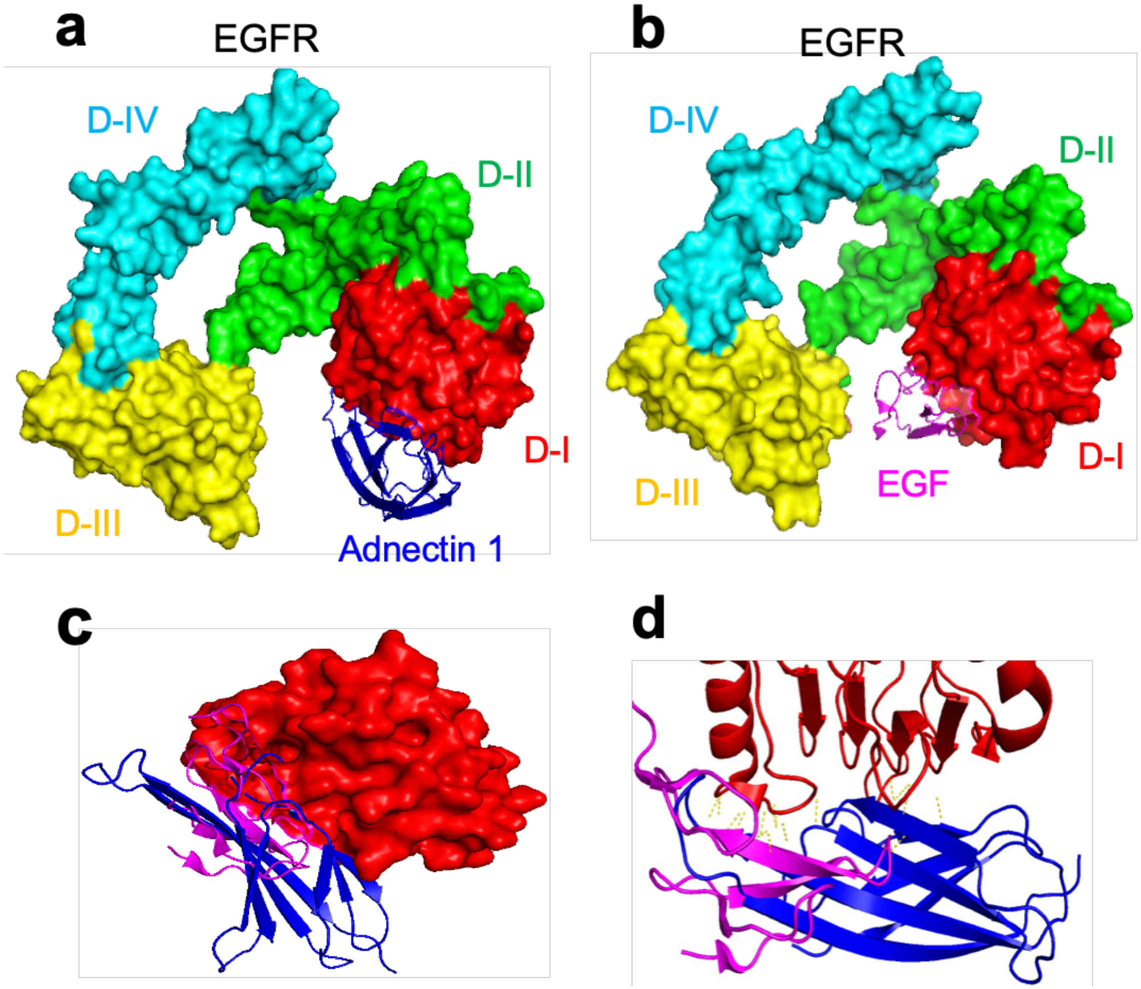


**<u>Supplementary Figure S17.</u> Structures of Adenctin1 and EGF in complex with EGFR.** **(a)** Specific binding between domain I (D-I) of EGFR and Adnectin1. Adnectin1 is illustrated in blue. EGFR surface is represented by various colors according to different domains (3QWQ.pdb).[17] **(b)** Interaction between EGF and D-I of EGFR. EGF is shown in magenta and EGFR as a surface by various colors (1NQL.pdb).[18] **(c)** Overlap of Adnectin1- and EGF-contacting surfaces on EGFR domain D-I is shown. Adnectin1 (blue) and EGF (magenta) are represented as cartoons but domain D-I is represented as a surface (red). **(d)** Adnectin1 (blue) and EGF (magenta) showing their hydrogen bonds (in yellow) with D-I of EGFR (red).

***17. Interconversion-dependent and interconversion-independent kinetic models of EGFR-Adnectin1 interactions.***

**Supplementary Table S16. Transition rate constants obtained for the interconversion-dependent kinetic model.** MATLAB (MathWorks, Natick, MA) was used to analyze raw single-channel data. The EGFR concentration was 20 nM. Values were provided as mean ± s.d. (n = 3), where *n* is the number of independently reconstituted nanopores. The association rate constants $k_{on-1}$ and $k_{on-2}$ obtained by this model were the same as those determined by the interconversion-independent kinetic model. The interconversion-independent and interconversion-dependent kinetic models are illustrated in **Supplementary Fig. S18**.

| [EGFR] (nM) | $k_{off-1}$ ($s^{-1}$) | $k_{12}$ ($s^{-1}$) | $k_{21}$ ($s^{-1}$) | $k_{off-2}$ ($s^{-1}$) |
|---|---|---|---|---|
| 20 | 15.6 ± 2.8 | 1.08 ± 0.20 | 0.46 ± 0.03 | 0.39 ± 0.09 |

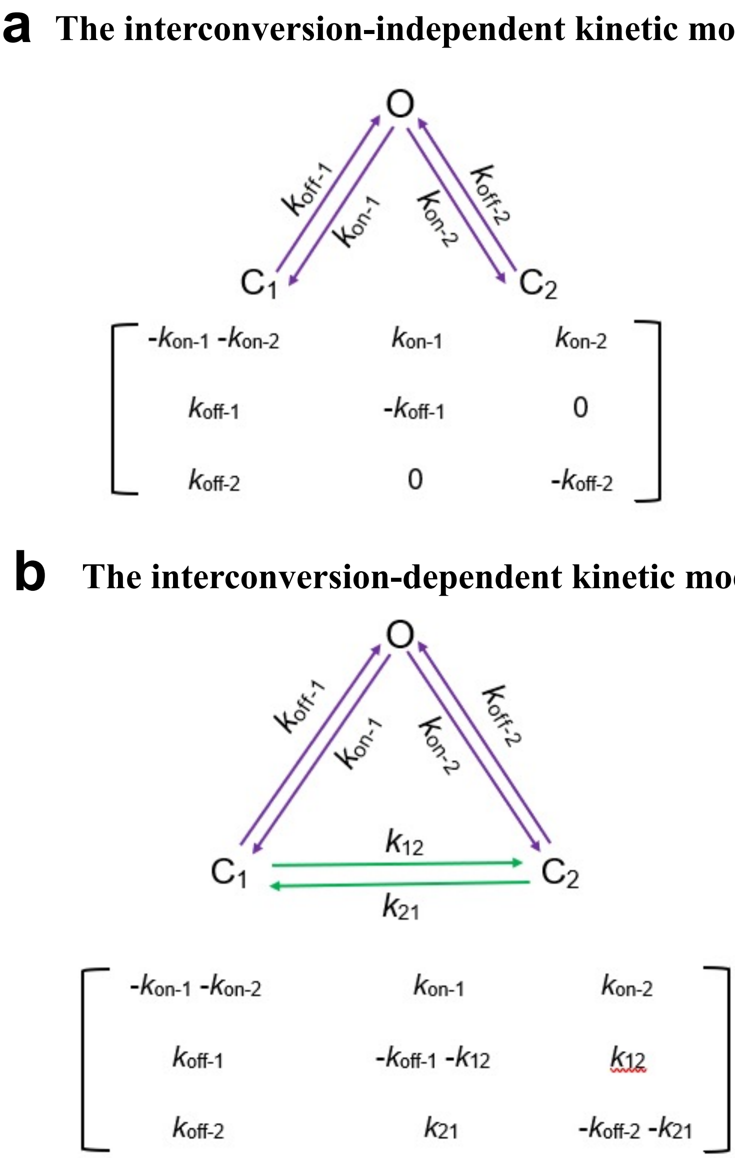


**<u>Supplementary Figure S18.</u> The interconversion-independent and interconversion-dependent kinetic model. (a)** The interconversion-independent kinetic model. This model assumes no transitions between different EGFR-captured substates, $C_1$ and $C_2$. The schematic of the model. O represents the EGFR-released substate. $C_1$ and $C_2$ indicate the short- and long-lived EGFR-captured substates, respectively. Q-matrix[11, 19-21] of the interconversion-independent three-substate kinetic model is also presented in the same panel. **(b)** The interconversion-dependent kinetic model. This model assumes transitions between the EGFR-captured substates $C_1$ and $C_2$. O represents the EGFR-released substate. $C_1$ and $C_2$ indicate the short- and long-lived EGFR-captured substates, respectively. The interconversion rate constants show transitions between different substates. Here, the first digit denotes the initial state, and the second digit indicates the final state. Q-matrix[11, 19-21] of the interconversion-dependent three-substate kinetic model is also presented in the same panel. Transitions from the EGFR-captured substates to EGFR-released substates and vice-versa are marked by magenta arrows. Transitions between the EGFR-captured substates $C_1$ and $C_2$ are marked in green.

*18. Positive-control experiments crosschecking the reactivity of the Adnectin1-tFhuA sensor for other protein analytes.*

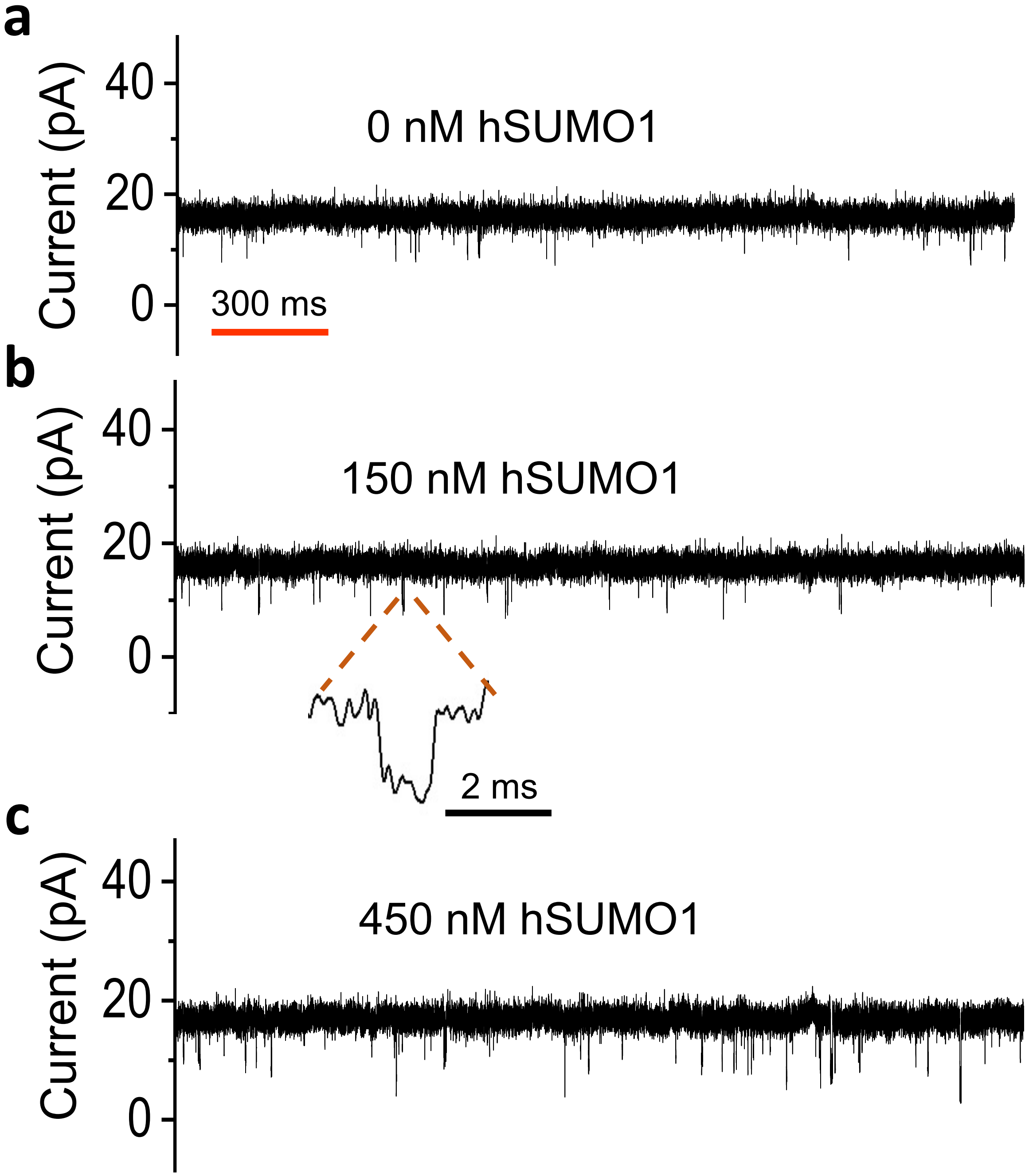


<u>Supplementary Figure S19.</u> **Positive-control experiment for testing the specificity of Adnectin1-tFhuA sensor in the presence of hSUMO1.** These are single-channel electrical recordings with Adnectin1-tFhuA when hSUMO1 was added to the *cis* chamber. The applied potential was +20 mV. **(a)** 0 nM hSUMO1. **(b)** 150 nM hSUMO1. **(c)** 450 nM hSUMO1. All recordings were performed in 300 mM KCl, 0.5 mM TCEP, 10 mM Tris-HCl, pH 8.0. The single-channel electrical traces were low-pass filtered at 2 kHz using an 8-pole Bessel filter. Specific hSUMO1-captured events were not observed with the Adnectin1-tFhuA sensor. These single-channel electrical signatures were replicated in n = 4 independent experiments.

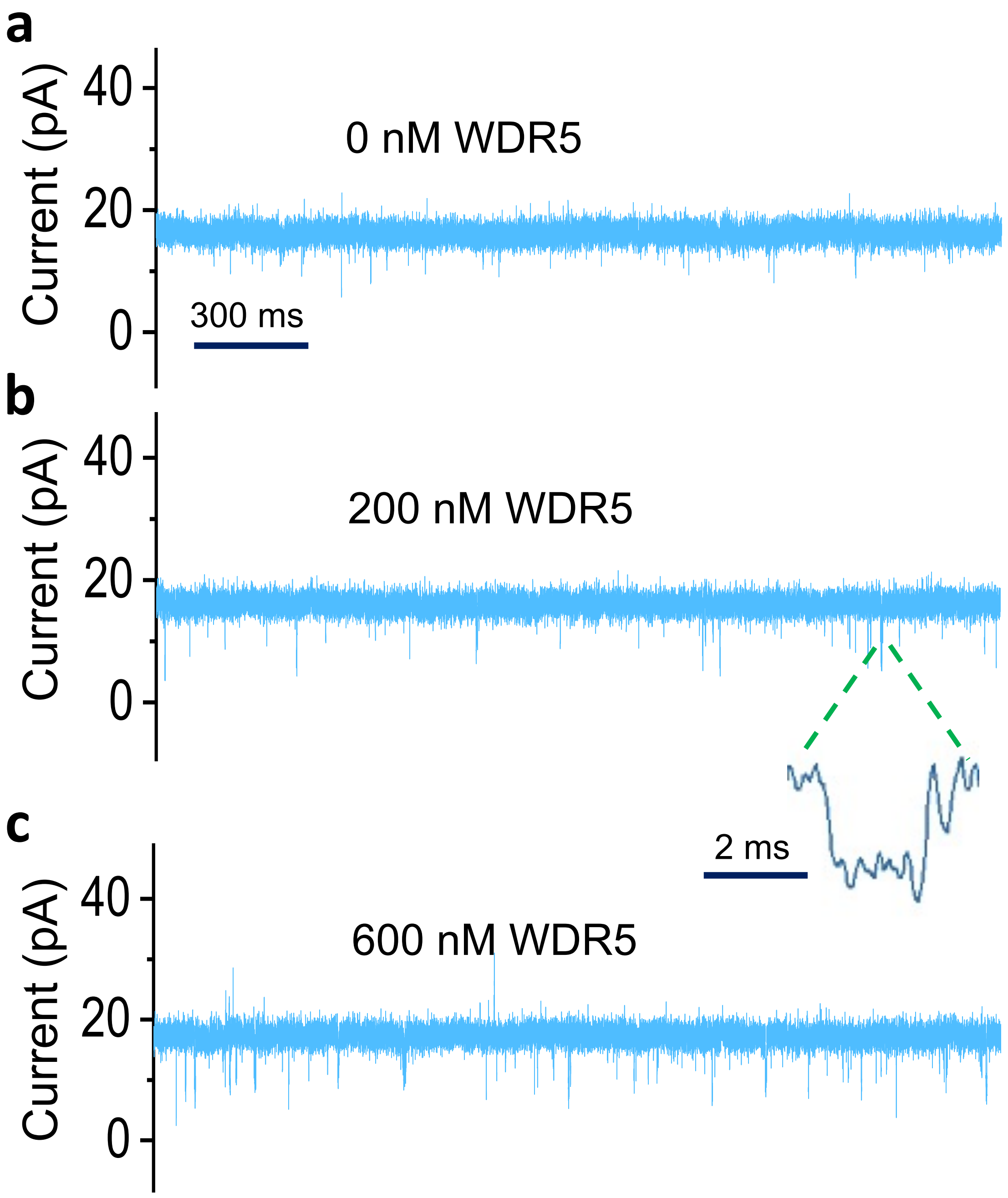


**<u>Supplementary Figure S20.</u> Positive-control experiment for testing the specificity of Adnectin1-tFhuA sensor in the presence of WDR5.** These are single-channel electrical recordings with Adnectin1-tFhuA when WDR5 was added to the *cis* chamber. The applied potential was +20 mV. **(a)** 0 nM WDR5. **(b)** 200 nM WDR5. **(c)** 600 nM hSUMO1. All recordings were performed in 300 mM KCl, 1 mM TCEP, 10 mM Tris-HCl, pH 8.0. The single-channel electrical traces were low-pass filtered at 2 kHz using an 8-pole Bessel filter. Specific WDR5-captured events were not observed with the Adnectin1-tFhuA sensor. These single-channel electrical signatures were replicated in n = 3 independent experiments.

***19. Time, rate, and equilibrium constants of EGFR–Adnectin1 interactions in the presence of mammalian serum.***

<u>**Supplementary Table S17.**</u> **The probability distribution of the two binding events noted with EGFR-Adnectin1 interactions in the absence and presence of 5% (v/v) fetal bovine serum (FBS).** These events were differentiated by the EGFR-captured durations. Individual experimental values were derived using event-list histograms in ClampFit (Axon) and their fits in a semilogarithmic representation. The maximum likelihood method[11, 12] and logarithm likelihood ratio (LLR)[13, 14, 16] tests were used for all fits to determine the best multi-exponential distribution model (**Methods**).

| 5% (v/v) FBS | $P_1$ | $P_2$ |
|---|---|---|
| - | 0.65 ± 0.04 | 0.35 ± 0.04 |
| + | 0.62 ± 0.06 | 0.38 ± 0.06 |

Values are mean ± s.d. from n = 3 independent experiments. $P_1$ and $P_2$ are the probabilities of the short- and long-lived EGFR-captured events, respectively. The other experimental conditions were the same as those stated in **Methods**.

<u>**Supplementary Table S18.**</u> **Mean values of the $\tau_{on}$ and $\tau_{off}$ constants of EGFR-Adnectin1 interactions in the absence and presence of FBS.**

| 5% (v/v) FBS | $\tau_{on}$ (s) | $\tau_{on-1}$ (s) | $\tau_{on-2}$ (s) | $\tau_{off-1}$ (s) | $\tau_{off-2}$ (s) |
|---|---|---|---|---|---|
| - | 0.40 ± 0.05 | 0.61 ± 0.043 | 1.2 ± 0.1 | 0.074 ± 0.027 | 0.97 ± 0.09 |
| + | 0.61 ± 0.09 | 0.98 ± 0.05 | 1.7 ± 0.5 | 0.062 ± 0.023 | 0.93 ± 0.14 |

Values in are mean ± s.d. ("-" stands for the absence of FBS from n = 3 independent experiments and "+" stands for the presence of 5% (v/v) FBS from n = 3 independent experiments). The applied transmembrane potential was +20 mV. The buffer solution contained 20 nM EGFR, 300 mM KCl, 10 mM Tris-HCl, pH 8.0.

<u>**Supplementary Table S19.**</u> **Mean values of the rate and equilibrium constants of EGFR-Adnectin1 interactions in the absence and presence of 5% (v/v) FBS.** Here, $K_D$ was calculated using the equation $K_D = k_{off}/k_{on}$.

| FBS | $k_{on-1}$ ($M^{-1}s^{-1}$) × $10^{-7}$ | $k_{on-2}$ ($M^{-1}s^{-1}$) × $10^{-7}$ | $k_{off-1}$ ($s^{-1}$) | $k_{off-2}$ ($s^{-1}$) | $K_{D-1}$ (nM) | $K_{D-2}$ (nM) |
|---|---|---|---|---|---|---|
| - | 8.8 ± 0.6 | 4.4 ± 0.7 | 15 ± 7 | 1.0 ± 0.1 | 191 ± 92 | 24 ± 2 |
| + | 5.1 ± 0.3 | 3.2 ± 0.9 | 18 ± 8 | 1.1 ± 0.2 | 347 ± 145 | 35 ± 5 |

Values are mean ± s.d. "-" stands for the absence of 5% (v/v) FBS from n = 3 independent experiments. "+" indicates the presence of 5% (v/v) FBS from n = 3 independent experiments. The applied transmembrane potential was +20 mV. The buffer solution contained 20 nM EGFR, 300 mM KCl, 10 mM Tris-HCl, pH 8.0.

***20. Side and top views of the three monobody-based sensors in complexes with their cognate protein analytes.***

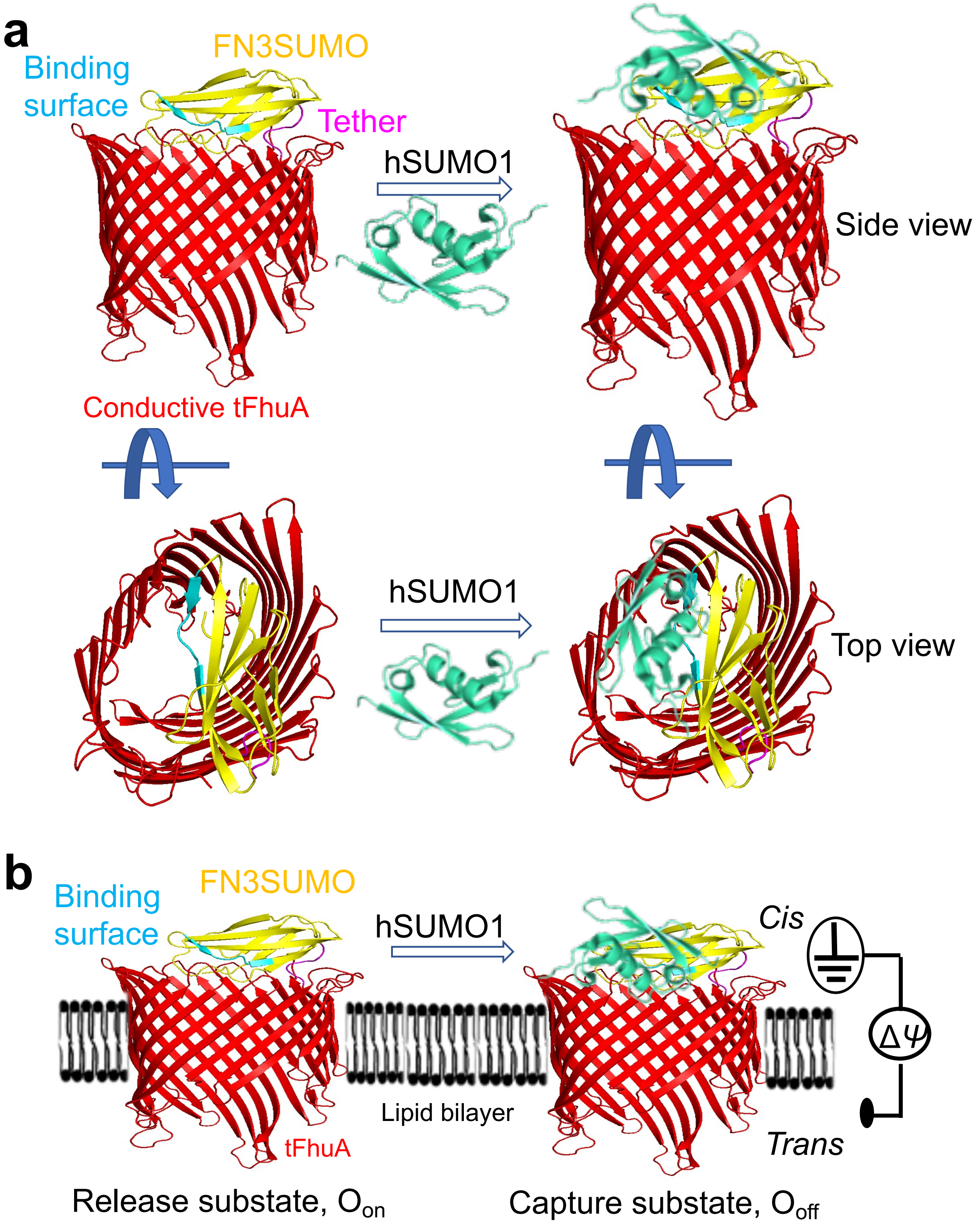


**<u>Supplementary Figure S21.</u> Side and top views of the modeled structure of the hSUMO1-FN3SUMO-tFhuA complex. (a)** FN3SUMO is oriented approximately 90° with respect to the central axis of tFhuA (*side view*), as judged by the AlphaFold2 approach.[8, 9] Binding of hSUMO1 to FN3SUMO almost fully blocks the pore opening (*top view*). **(b)** This panel shows the functionally reconstituted FN3SUMO-tFhuA into a lipid bilayer in the hSUMO1-released ($O_{on}$) and hSUMO1-captured ($O_{off}$) substates.

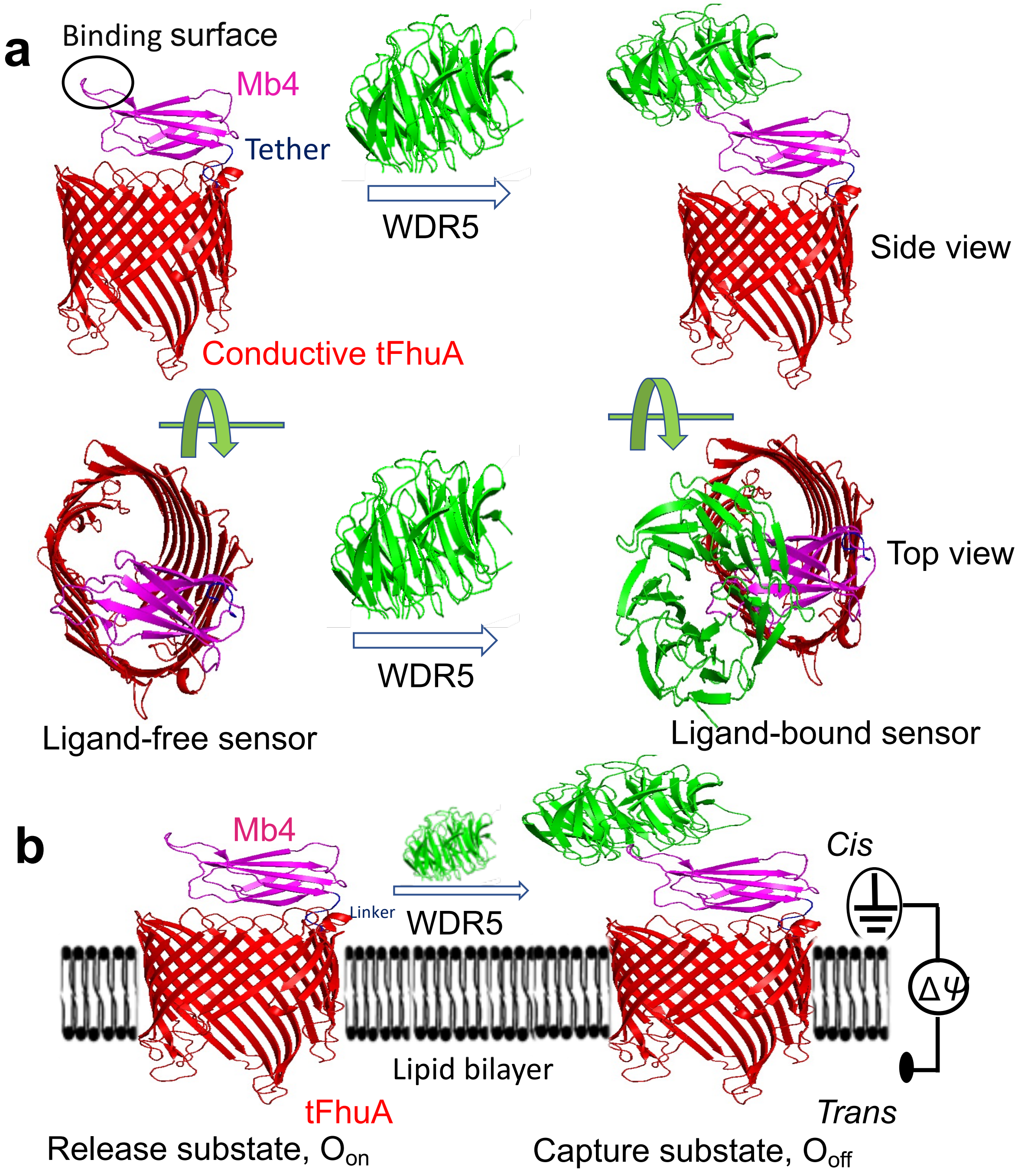


<u>**Supplementary Figure S22.**</u> **Side and top views of the modeled structure of the WDR5-Mb4-tFhuA complex. (a)** Mb4 is oriented approximately 90° with respect to the central axis of tFhuA (*side view*), as judged by the AlphaFold2 approach.[8, 9] Binding of WDR5 to Mb4 partly blocks the pore opening (*top view*). **(b)** This panel shows the functionally reconstituted Mb4-tFhuA into a lipid bilayer in the WDR5-released ($O_{on}$) and WDR5-captured ($O_{off}$) substates.

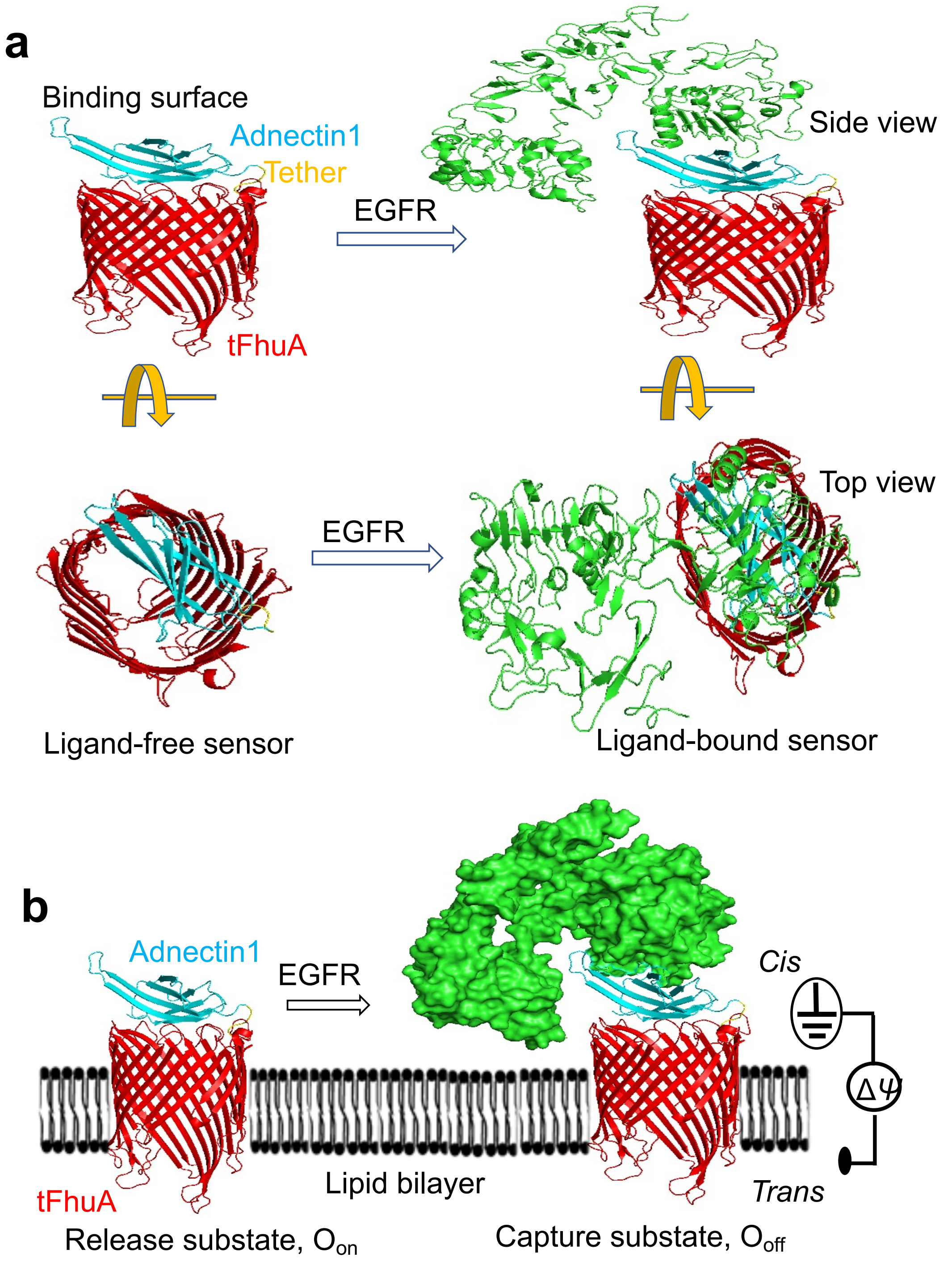


<u>**Supplementary Figure S23.**</u> **Side and top views of the modeled structure of the EGFR-Adnectin1-tFhuA complex. (a)** Adnectin1 is oriented approximately 90° with respect to the central axis of tFhuA (*side view*), as judged by the AlphaFold2 approach.[8, 9] Binding of EGFR to Adnectin1 almost fully blocks the pore opening (*top view*). **(b)** This panel shows the functionally reconstituted Adnectin1-tFhuA into a lipid bilayer in the EGFR-released ($O_{on}$) and EGFR-captured ($O_{off}$) substates.

*21. The three monobody-containing sensors exhibit different electrical and kinetic signatures.*

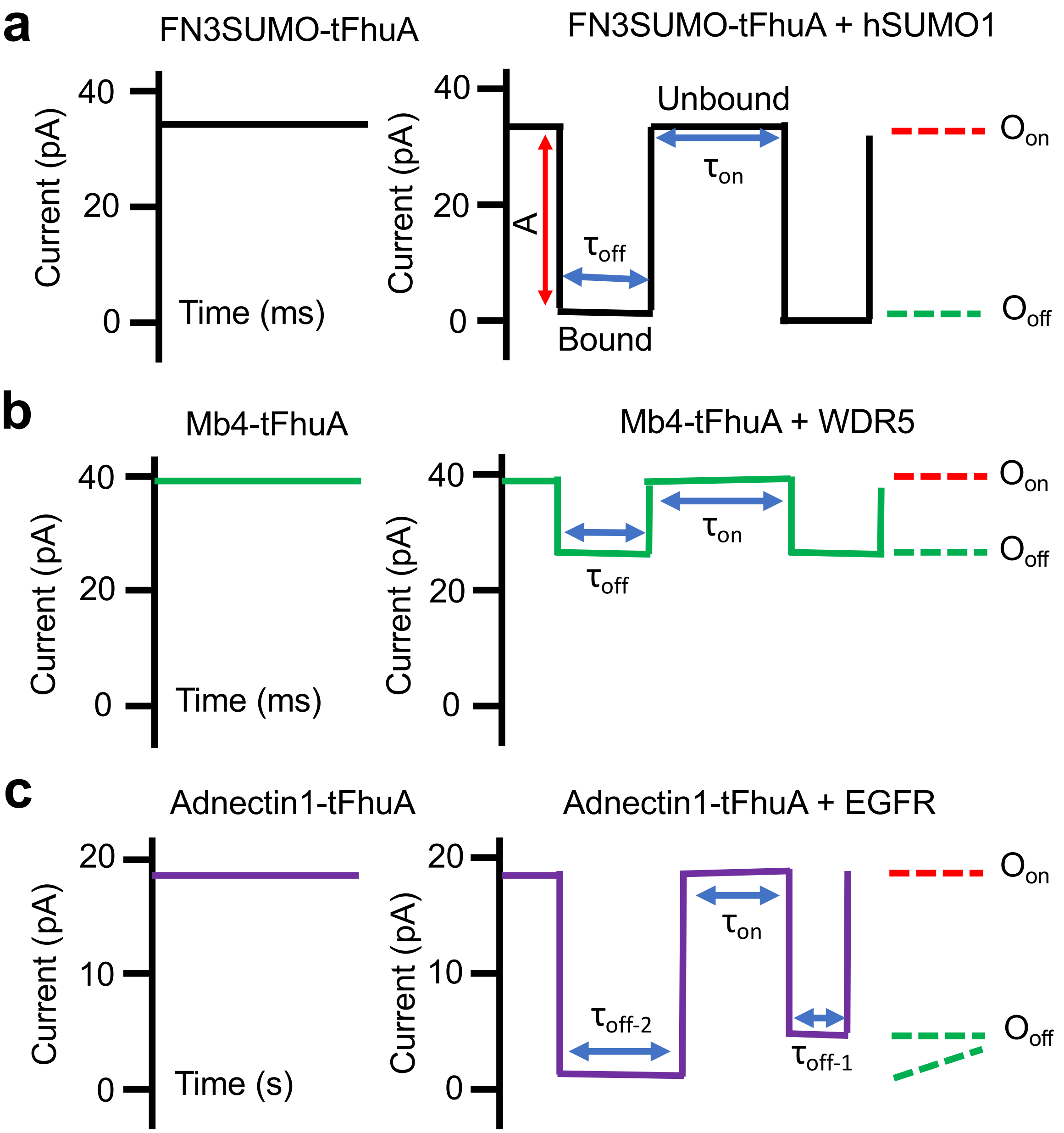


__Supplementary Figure S24.__ **Schematic of the relative current blockades acquired with FN3SUMO-tFhuA, Mb4-tFhuA, and Adnectin1-tFhuA. (a)** Graphic representation of stochastic sensing of hSUMO1 using an FN3SUMO-tFhuA sensor, which maintains a basal open-state current (*left panel*). When hSUMO1 is added to the *cis* side, the analyte produces large-amplitude, long-lived current transitions between two current substates (*right panel).* **(b)** Mb4-tFhuA sensor maintains a basal open-state current (*left panel*). When added to the *cis* side, WDR5 produces low-amplitude, long-lived current transitions (*right panel*). **(c)** Adnectin1-tFhuA protein maintains a basal open-state current (*left panel*). When added to the *cis* side, EGFR produces large-amplitude, long-lived current transitions (*right panel*).

**<u>Supplementary Table S20.</u> Comparison of capture durations and current blockades produced by different protein analytes.**

| Protein analyte | Capture duration (ms) | Normalized current blockades $A/I_0$ (%) |
|---|---|---|
| **hSUMO1** | 13 ± 1 | 92 ± 1 |
| **WDR5** | 14 ± 1 | 14 ± 1 |
| **EGFR** | 1010 ± 14<br>85 ± 3 | 65 ± 2<br>86 ± 1 |

Capture durations represent mean ± s.e.m. using the fits in **Supplementary Fig. S25**. Values of normalized current blockades are mean ± s.d. from n = 3 independent experiments. These values are determined at concentrations of 65 nM hSUMO1, 50 nM WDR5, and 40 nM EGFR.

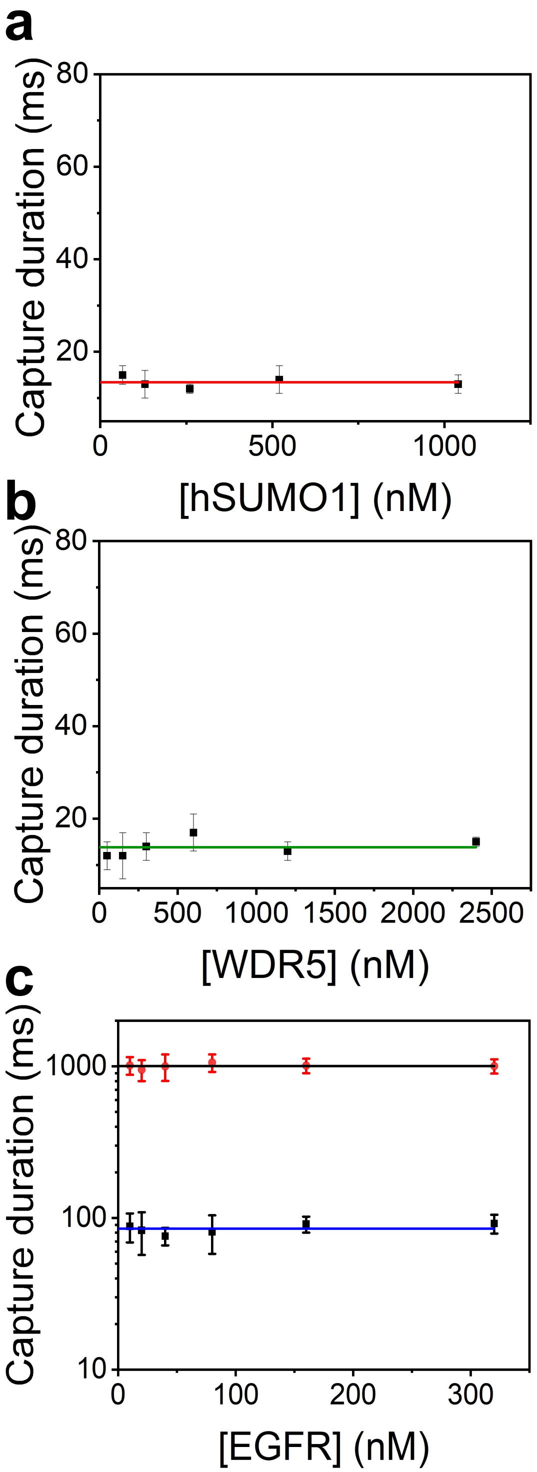


**<u>Supplementary Figure S25.</u> Linear fits of the analyte-captured durations acquired with FN3SUMO-tFhuA, Mb4-tFhuA, and Adnectin1-tFhuA sensors. (a)** hSUMO1. **(b)** WDR5. **(c)** EGFR. The applied transmembrane potentials were +40 mV, +40 mV, and +20 mV, respectively. All the other experimental conditions are the same as those mentioned in **Methods**. Plot values indicate mean ± s.d. from n = 3 independent experiments.

**22. Steady-state FP anisotropy curves of hSUMO1−FN3SUMO-tFhuA and WDR5−Mb4-tFhuA interactions.**

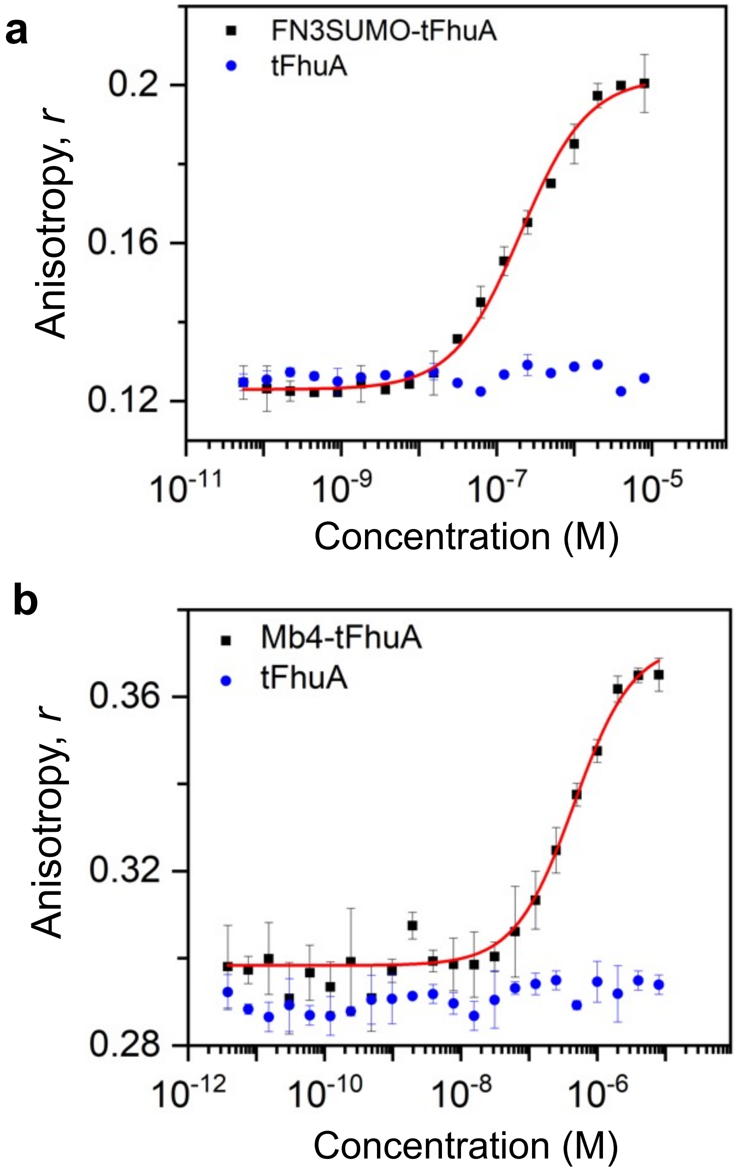


__Supplementary Figure S26.__ **Steady-state FP anisotropy curves of hSUMO1−FN3SUMO-tFhuA and WDR5−Mb4-tFhuA interactions. (a)** hSUMO1 was labelled with fluorescein. The final concentration of the labeled hSUMO1 in each well was 50 nM. **(b)** WDR5 was labelled with rhodamine. The final concentration of the labeled WDR5 in each well was 50 nM. The labeled hSUMO1 and WDR5 were titrated against FN3SUMO-tFhuA and Mb4-tFhuA, respectively. Data indicate mean ± s.d. from n = 3 independent experiments.

*23. List of primers used in this study.*

<u>Supplementary Table S21.</u> **List of primers used in this study.**

| Primer name | Sequences (5′-3′) |
|---|---|
| FN3SUMO_for | CTTTAAGAAGGAGATATACAAATGGGTAGCCCGAGCGTTCC GGGC |
| FN3SUMO_rev | CTGAACTTCTTTCAGGCTGCCGCCGCTGCCGCCGGTGGTAAC GCTAACGCTGC |
| Mb4_for | CTTTAAGAAGGAGATATACAAATGGGATCTTCTGTTCCGAC C |
| Mb4_rev | CTGAACTTCTTTCAGGCTGCCGCCGCTGCCGCCGGTACGGTA GTTAATCGAG |
| Adnectin1_for | CTTTAAGAAGGAGATATACCATGGGGGTATCTGACGTG |
| Adnectin1_rev | CAGGCTGCCGCCGGATCCGCCTTGCGAAGGTTTGTCGATTTC |
| tFhuA_for | CCGGCGGATCCGGCGGCAGCCTGAAAGAAG |
| tFhuA_rev | GATCCTCGAGTTAAAAACGAAAGGTTGCGGTGGC |
| Egfr_ecd_for | TCCGCTAGCGCCACCATGGTGCGACCCTCCGGGACG |
| Egfr_ecd_rev | TCGAGATCTTTAGTGATGATGATGATGATGGGACGGGATCT TAGGCCCATTCGTTGG |
| tFhuA_cys_for | TGTGAAGGTAGCAGCGGTCCGTATCG |
| tFhuA_cys_rev | TGAATTATAGCCAAACCAGGCATTAATATCATTGCGC |

*24. SDS-PAGE analysis of purified EGFR.*

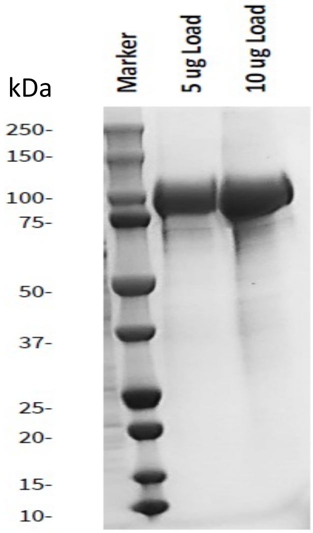


<u>Supplementary Figure S27.</u> **SDS-PAGE gel of purified EGFR.** The EGFR was purified using a polyhistidine-tag column. The protein purity was checked on a 4-20%-gradient SDS-PAGE gel. The expected molecular weight (MW) of EGFR is 69.6 kDa. However, we observed a higher apparent MW due to glycosylation.

***25. PDB entries used for visualization and molecular graphics of all protein structures employed in this study.***

**<u>Supplementary Table S22.</u> PDB entries used for visualization and molecular graphics of all protein structures employed in this study.**

| Protein/Protein complex | PDB accession code | Source's URL |
|---|---|---|
| FhuA | 1BY3[22] | https://www.rcsb.org/structure/1by3 |
| Parent FN3 | 1FNF[23] | https://www.rcsb.org/structure/1FNF |
| ySMB9-hSUMO1 | 3RZW[24] | https://www.rcsb.org/structure/3RZW |
| Mb4-WDR5 | 6BYN[25] | https://www.rcsb.org/structure/6BYN |
| EGF-EGFR | 1NQL[18] | https://www.rcsb.org/structure/1NQL |
| Adnectin1-EGFR | 3QWQ[17] | https://www.rcsb.org/structure/3QWQ |

***26. Supplementary references.***
